# Temporal organization of shared brain states characterizes heterogeneity in major depressive disorder

**DOI:** 10.64898/2026.09.28.754910

**Authors:** Chunyu Pan, Yue Tu, Fei Wang, Kangguang Lin, Shulan Qiu, Weixiong Zhang, the REST-meta-MDD Consortium

## Abstract

Major depressive disorder (MDD) is biologically heterogeneous, but how this variability is expressed in the temporal organization of brain activity remains unclear. Using resting-state functional MRI from 2,225 participants, we applied hidden Markov modeling to identify recurrent brain states and quantify their temporal organization. Five states formed a shared repertoire across individuals with MDD and healthy controls, while MDD-related alterations were distributed across states, functional systems, and temporal properties. Within MDD, unsupervised clustering of state occupancy, persistence, and switching delineated two prototype neurodynamic architectures characterized by preferential deployment of ACS/VSS or MAS/DMS states within a preserved state repertoire and broadly similar transition backbone. The architectures showed different within-group brain–symptom associations and symptom-network patterns. Connectome transcriptome analyses organized architecture–state imaging patterns into three transcriptomic state families shared across architectures, and contrasting group-level deployment patterns were recovered in an independent MDD cohort. These findings suggest that MDD heterogeneity is expressed not only through spatially distributed neural alterations, but also through differences in how a common repertoire of recurrent brain states is organized over time.

## 1. Introduction

Biological heterogeneity represents a fundamental challenge in understanding psychiatric disorders. Major depressive disorder (MDD), a prevalent disorder and major contributor to the global burden of disease [1-3], provides a prominent example that individuals meeting the same diagnostic criteria can differ substantially in symptom profiles, brain organization, and treatment response [4-6]. Despite advances in neuroimaging and molecular neuroscience, the dimensions of brain organization underlying this variability remain incompletely understood. Diagnostic categories define clinically recognizable populations but inevitably average across potentially meaningful differences between individuals [7]. A central question is therefore not only which neural abnormalities characterize MDD on average, but also along which dimensions brain organization varies within MDD and whether this variation forms reproducible organizational patterns.

Resting-state functional MRI (rs-fMRI) studies have provided substantial evidence that MDD is associated with distributed alterations in large-scale brain organization, including the default-mode, salience, executive, and sensory networks [8, 9]. These findings have established that the neural alterations associated with MDD extend across multiple functional systems rather than being confined to a single brain region or circuit. Brain function arises from the coordinated activity of these systems, which can combine into persistent or recurrent configurations. Individuals with MDD may therefore differ not only in the magnitude of alterations within particular regions or networks, but also in how distributed systems are jointly configured into recurring patterns of brain activity [5, 6, 10]. Determining whether these configurations exhibit structured and reproducible variation across individuals may help clarify the dimensions of brain organization underlying MDD heterogeneity [4, 11].

Neuroimaging-based stratification approaches have begun to address this challenge by identifying patient profiles based on functional connectivity, regional activity, and individualized deviations from normative brain organization [12-14]. Many identified profiles involve large-scale functional systems, particularly fronto-limbic and default-mode networks [15-17]. However, these approaches have predominantly relied on static or time-averaged representations of brain organization. Although these representations characterize relatively stable spatial properties, they compress the temporal structure through which brain activity configurations emerge, persist, recur, and transition over time [18]. Studies using dynamic functional connectivity and co-activation patterns have further demonstrated the relevance of temporal variation to neuroimaging-based stratification [19, 20]. However, whether the occupancy, persistence, switching, and transition of recurrent brain configurations collectively form a structured dimension of individual variation within MDD remains unclear [21].

Brain activity can be conceptualized as a continuous reconfiguration among recurrent functional states, in which distributed neural systems are coordinated into transient but reproducible patterns of activity [22-24]. These brain states represent organized configurations of large-scale networks rather than isolated activation of individual regions, reflecting coordinated interactions that support cognitive flexibility, adaptive behavior, and the balance between functional integration and segregation [25-27]. Importantly, brain-state organization contains two complementary but distinct components: the functional configuration of individual states, which describes the systems jointly engaged within a recurrent state, and the temporal organization of the state repertoire, which describes how states are dynamically deployed across time, including their occupancy, persistence, variability, and transitions [28, 29]. From this perspective, heterogeneity does not necessarily require different state repertoires or discrete disease entities; it may arise from different temporal organizations of a shared dynamical repertoire. We therefore hypothesized that MDD heterogeneity may partly arise from differences in the temporal organization of shared recurrent brain-state repertoires, resulting in distinct neurodynamic architectures across individuals [30, 31].

Here, we tested these hypotheses using a large multisite resting-state fMRI dataset comprising individuals with MDD and healthy controls. We first identified a common repertoire of recurrent brain states and characterized their functional configurations and temporal properties. After examining group-level alterations in MDD, we assessed whether individual differences in state occupancy, persistence, switching, and transitions formed structured patterns within the disorder. We refer to the resulting prototype patterns of temporal state deployment as neurodynamic architectures. We then characterized their state-preference and transition profiles, examined architecture-stratified brain–symptom associations and symptom-network patterns, and used connectome-transcriptome analyses to provide a molecular context for the recurrent state configurations. Finally, we applied the discovery-derived state framework to an independent MDD cohort to assess whether the principal temporal organization could be recovered across datasets. Together, these analyses tested whether the temporal organization of shared brain states characterizes a structured and cross-cohort dimension of heterogeneity within MDD.

## 2. Results

### 2.1 Study overview

We analyzed resting-state functional MRI data from 2,225 individuals, including 1,177 patients with MDD and 1,048 healthy controls (HCs), from the REST-meta-MDD consortium (Table 1). The overall study design is illustrated in *Fig. 1*. First, hidden Markov modeling (HMM) was applied to identify a recurrent brain-state repertoire underlying large-scale neural activity and to decode participant-specific state sequences. State-specific functional configurations and temporal features, including fractional occupancy (FO), lifetime (LT), switching rate (SR), and transition probability (TP), were subsequently quantified to characterize individual brain-state organization (*Fig. 1A*). Second, we characterized MDD-related alterations in brain-state features and examined their associations with depressive symptom severity. Individual profiles of state occupancy, persistence, and switching were then analyzed using unsupervised clustering to identify structured patterns of temporal organization within the MDD cohort. These patterns were summarized as prototype neurodynamic architectures and characterized according to their state-preference and transition profiles. Post hoc classification analyses further assessed the separability of each architecture from HCs (*Fig. 1B*). Third, we examined architecture-stratified brain–symptom associations and symptom-network patterns and used connectome-transcriptome analyses to investigate the molecular context of recurrent state configurations. Finally, the discovery-derived brain-state framework was applied to an independent cohort of 65 patients with MDD to evaluate the cross-cohort recovery of the principal temporal organization (*Fig. 1C*). Together, these analyses provided a multilevel characterization of recurrent brain-state organization, its variation within MDD, and its clinical, molecular, and cross-cohort context.

**Table 1.** Demographic information of the study participants.

| Variable | Total | HC | MDD | $\chi^2/t$ | P value |
| --- | --- | --- | --- | --- | --- |
| Number | $n=2225$ | $n=1048$ | $n=1177$ | -- | -- |
| Gender-Male | 38.50% | 41.14% | 36.29% | $\chi^2 = 2.17$ | $p = 0.14$ |
| Gender-Female | 61.50% | 58.86% | 63.71% |  |  |
| Age (Mean±SD) | $36.25 \pm 15.30$ | $36.30 \pm 15.97$ | $36.20 \pm 14.69$ | $\vert t\vert = 0.15$ | $p = 0.88$ |
| HAMD (Mean±SD) | -- | -- | $20.26 \pm 7.14$ | -- | -- |

All P values are two-sided. HC, healthy controls; MDD, major depressive disorder; SD, standard deviation. HAMD-17 scores were available for a subset of participants.

**Figure 1.**
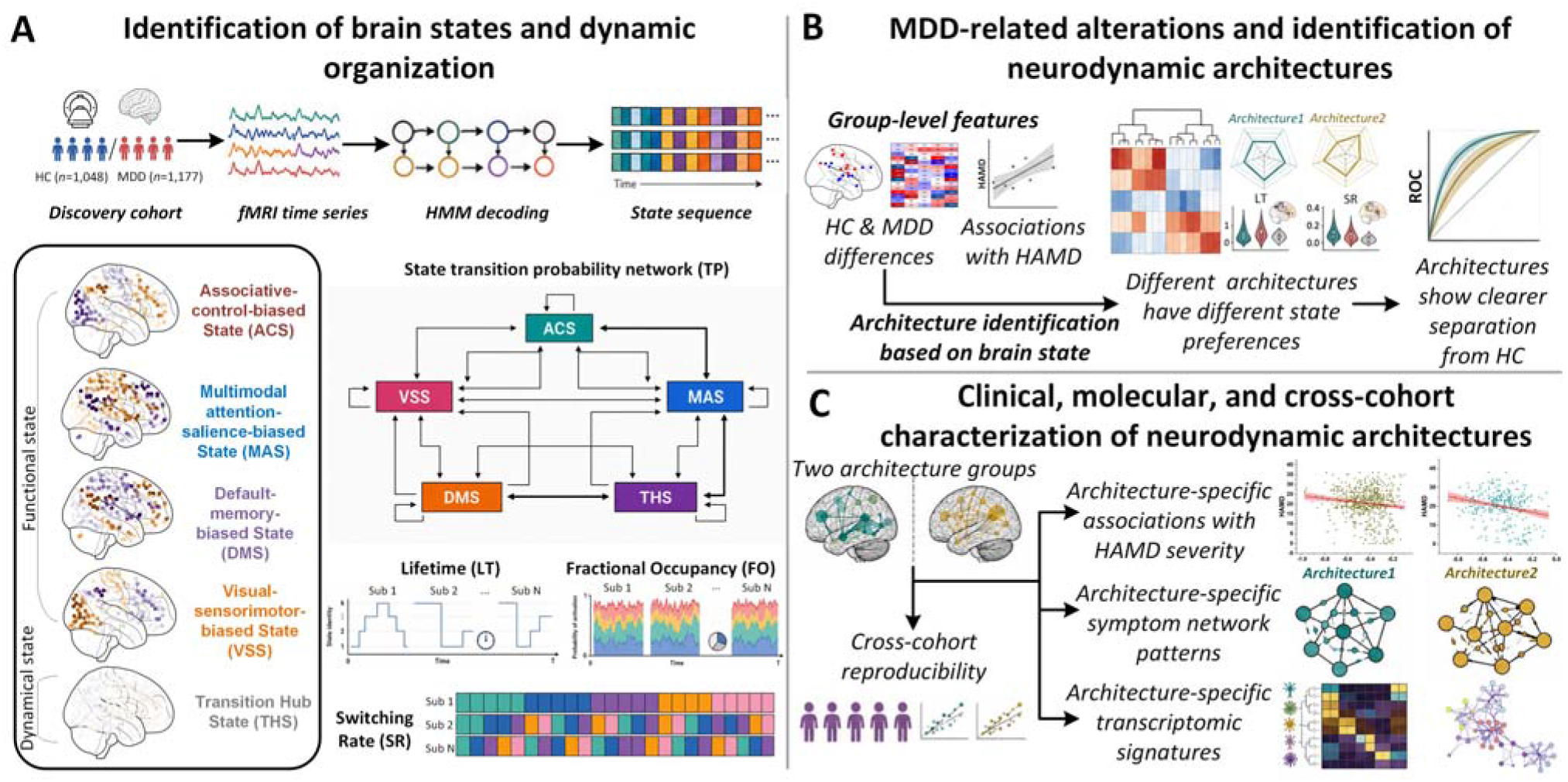
Overview of the study. (A) Identification of brain states and their dynamic organization. Resting-state functional MRI data were modeled using hidden Markov modeling (HMM) to identify recurrent states of spontaneous brain activity and decode participant-specific state sequences. State-specific functional configurations and temporal properties were characterized using fractional occupancy (FO), lifetime (LT), switching rate (SR), and transition probability (TP). (B) MDD-related alterations and identification of neurodynamic architectures. Brain-state features were compared between individuals with MDD and healthy controls (HCs), and their associations with Hamilton Depression Rating Scale (HAMD) severity were examined. Unsupervised clustering of individual temporal profiles was then performed within the MDD cohort to identify prototype neurodynamic architectures. The identified architectures were characterized according to their state-preference and transition profiles, followed by post hoc evaluation of their separability from HCs. (C) Clinical, molecular, and cross-cohort characterization. Brain–symptom associations and symptom-network patterns were examined separately within each architecture. Connectome-transcriptome analyses were used to characterize the molecular context of recurrent state configurations, and the discovery-derived brain-state framework was applied to an independent MDD cohort to assess the cross-cohort recovery of the principal temporal organization.

### 2.2 A repertoire of recurrent brain states captures functional specialization and temporal organization

Across the entire cohort, hidden Markov modeling identified five recurrent brain states that together constituted a compact repertoire of large-scale spontaneous brain activity (*see methods*). Each state was defined by a distinct regional activity configuration and temporal profile. These states captured both the functional configuration and dynamical organization of brain activity and served as the foundation for all subsequent analyses.

Four states exhibited distinct brain activity configurations (*Fig. 2A*). The Associative-control-biased State (ACS) was characterized by increased activity in frontoparietal, salience, and memory-retrieval systems, accompanied by reduced activity in visual and sensorimotor systems. The Multimodal attention-salience-biased State (MAS) showed increased activity across auditory, cingulo-opercular, dorsal-attention, salience, and sensorimotor systems, together with reduced activity in default-mode and memory-retrieval systems. In contrast, the Default-memory-biased State (DMS) exhibited preferential activity in default-mode and memory-retrieval systems and reduced activity across attention-related and sensorimotor systems. The Visual-sensorimotor-biased State (VSS) was distinguished by increased activity in visual and sensorimotor systems, accompanied by reduced activity in frontoparietal, salience, and memory-retrieval systems. Unlike these four states, the Transition Hub State (THS) showed a comparatively balanced activity configuration, with activity across most functional systems remaining close to the across-state average.

**Figure 2.**
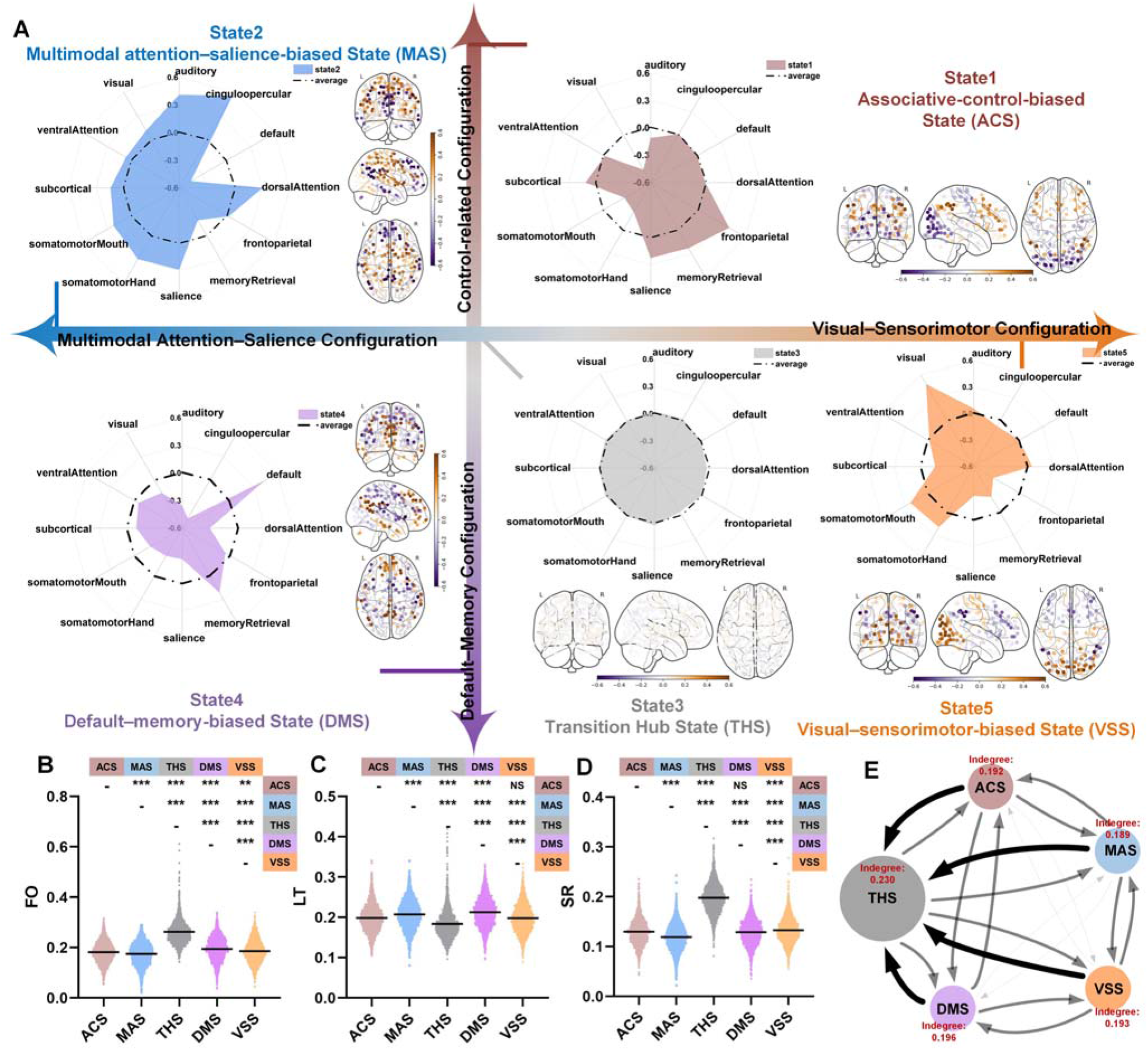
A common recurrent state space captures functional specialization and temporal organization. (A) Activity configurations of the five recurrent brain states. Hidden Markov modeling identified the Associative-control-biased State (ACS), Multimodal attention–salience-biased State (MAS), Transition Hub State (THS), Default–memory-biased State (DMS), and Visual–sensorimotor-biased State (VSS). Radar plots show state-specific mean activity across 12 large-scale functional systems, with dashed contours indicating the corresponding across-state means; brain maps display the regional activity configuration of each state. The state labels and schematic ACS–DMS and MAS–VSS contrasts summarize the dominant relative activity configurations rather than directly measured psychological processes or statistically derived dimensions. THS is positioned between the four configuration-biased states because of its comparatively balanced activity profile. (B–D) Temporal properties of the five brain states. Violin plots show fractional occupancy (FO), lifetime (LT), and switching rate (SR). FO represents the proportion of time occupied by each state, LT represents the mean duration of consecutive visits to a state, and SR represents the mean temporal change in posterior state probability between adjacent time points. Matrices above the distributions summarize pairwise comparisons between states. (E) Transition structure of the recurrent brain-state repertoire. The directed transition probability (TP) network illustrates transitions among the five states. Node size represents fractional occupancy, arrows indicate transition direction, edge width represents transition probability, and red labels denote weighted indegree. THS showed the greatest weighted incoming transition strength and, together with its high occupancy, short lifetime, and high switching rate, occupied a hub-like position within the observed transition structure. \**P* < 0.05, \*\**P* < 0.01, and \*\*\**P* < 0.001, FDR-corrected for multiple comparisons.

The THS was distinguished primarily by its temporal properties. It exhibited the highest fractional occupancy (FO), shortest lifetime (LT), and highest switching rate (SR) among the five states (*Fig. 2B–D* and *Supplementary Table S1*). Transition probability analysis further showed that the THS occupied a central position within the state-transition architecture and received the greatest incoming transition strength (*Fig. 2E* and *Supplementary Table S2*). Its high occupancy, short persistence, rapid switching, and prominent incoming transitions collectively supported its characterization as a transition hub within the recurrent state repertoire.

Together, these findings indicate that large-scale spontaneous brain activity was organized into a structured state repertoire. These brain states and their associated temporal features were subsequently used to examine MDD-related alterations and inter-individual variability in brain-state organization.

### 2.3 Major depressive disorder is associated with distributed alterations in brain state organization

To characterize MDD-related differences in brain-state organization, we compared state-specific brain activity configurations and temporal properties between individuals with MDD and HCs (*Fig. 3A–E* and *Supplementary Table S3*). Differences were distributed across multiple states and functional systems, involving both regional activity configurations and temporal features. Within ACS, individuals with MDD showed higher state-specific mean sensorimotor activity accompanied by lower salience and subcortical activity. MAS exhibited lower visual activity together with higher default-mode and ventral-attention activity, whereas DMS showed the opposite pattern, with increased visual activity and lower default-mode and ventral-attention activity. VSS exhibited comparatively widespread differences involving auditory, dorsal -attention, sensorimotor, salience, and subcortical systems. By comparison, THS showed relatively limited differences between groups. Alterations in temporal properties, including FO, LT, and SR, were also distributed across states and did not indicate a uniform shift in the temporal organization of the recurrent state repertoire.

**Figure 3.**
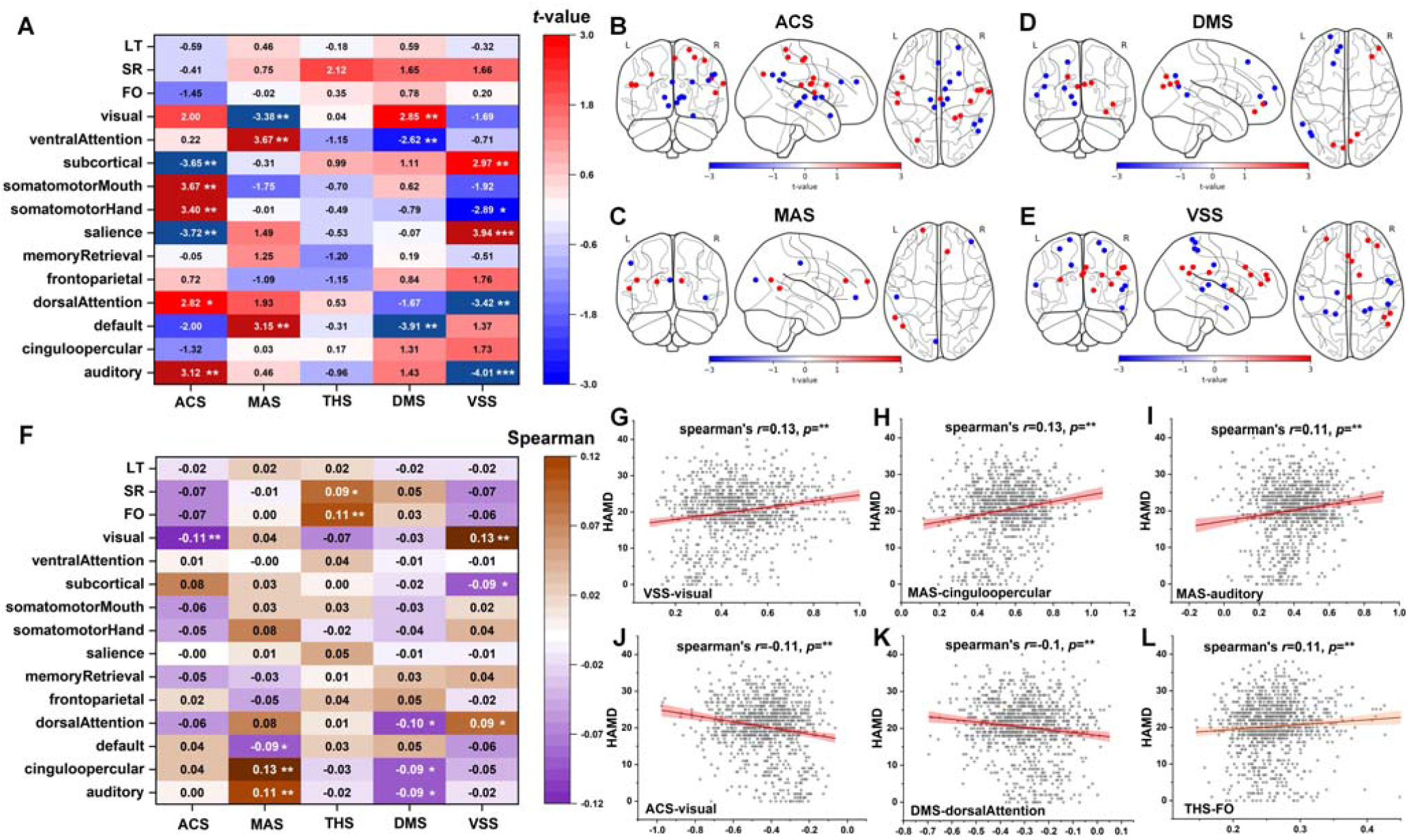
Distributed state-configuration alterations and symptom associations in pooled MDD. **(A)** Group differences in brain-state features between individuals with MDD and HC. Group differences in state-specific functional configurations and temporal properties were assessed using two-sample *t-tests* between MDD and HC groups. Analyses were performed across regional activity patterns, functional community representations, and dynamic features of the five recurrent brain states; **(B-E)** Spatial distribution of MDD-related alterations in state-specific functional configurations. Significant group differences ROI were visualized across brain; **(F)** Associations between brain-state features and depressive symptom severity within the MDD cohort. Associations between state-specific functional features, temporal dynamic measures, and depressive symptom severity assessed by the HAMD were evaluated using Spearman correlation analyses; **(G-L)** Representative brain state and symptom associations. Scatter plots illustrate representative significant associations between state-specific brain-state features and HAMD scores within individuals with MDD; *\*P* < 0.05, *\*\*P* < 0.01, and *\*\*\*P* < 0.001. All statistical tests were corrected for multiple comparisons using FDR.

We next examined associations between brain-state features and depressive symptom severity within the MDD cohort (*Fig. 3F–L* and *Supplementary Table S4*). Significant associations with HAMD scores were identified across both state-specific activity configurations and temporal features. Greater depressive symptom severity was associated with higher visual activity in VSS (*r* = 0.13), higher cingulo-opercular and auditory activity in MAS (*r* = 0.13 and 0.11, respectively), and higher fractional occupancy of THS (*r* = 0.11). Conversely, HAMD scores were negatively associated with visual activity in ACS (*r* = -0.11) and dorsal-attention activity in DMS (*r* = -0.10). Additional significant associations were observed across multiple state–system combinations and temporal features in the full correlation matrix (*Fig. 3F*). Although the individual effect sizes were small, the associations were distributed across different states and features rather than concentrated in a single component of brain-state organization.

Together, these findings show that MDD-related alterations in activity configurations are distributed across a shared recurrent brain-state repertoire, with symptom associations extending across both activity configurations and temporal properties. This state-dependent and directionally heterogeneous pattern further suggests that diagnostic group-level comparisons may capture only part of the inter-individual variation in brain-state organization within MDD. Accordingly, we next examined whether individual differences in the temporal deployment of the shared state repertoire formed structured patterns within MDD.

### 2.4 Neurodynamic stratification reveals two brain state organizational architectures in major depressive disorder

To determine whether interindividual variability in brain-state organization reflected structured heterogeneity within MDD, we first used exploratory sparse principal component analysis to characterize the major sources of variation across the 75 candidate brain-state features. Unsupervised clustering subsequently identified two neurodynamic architectures, whose stability was further evaluated across 1,000 repeated 80% subsampling iterations (see *Methods* and *Supplementary Material S2*). The two groups should be interpreted as prototype expressions along a dominant dimension of temporal organization rather than as sharply separated categories. Demographic and clinical characteristics of the two architectures are summarized in *Table 2*.

**Table 2.**
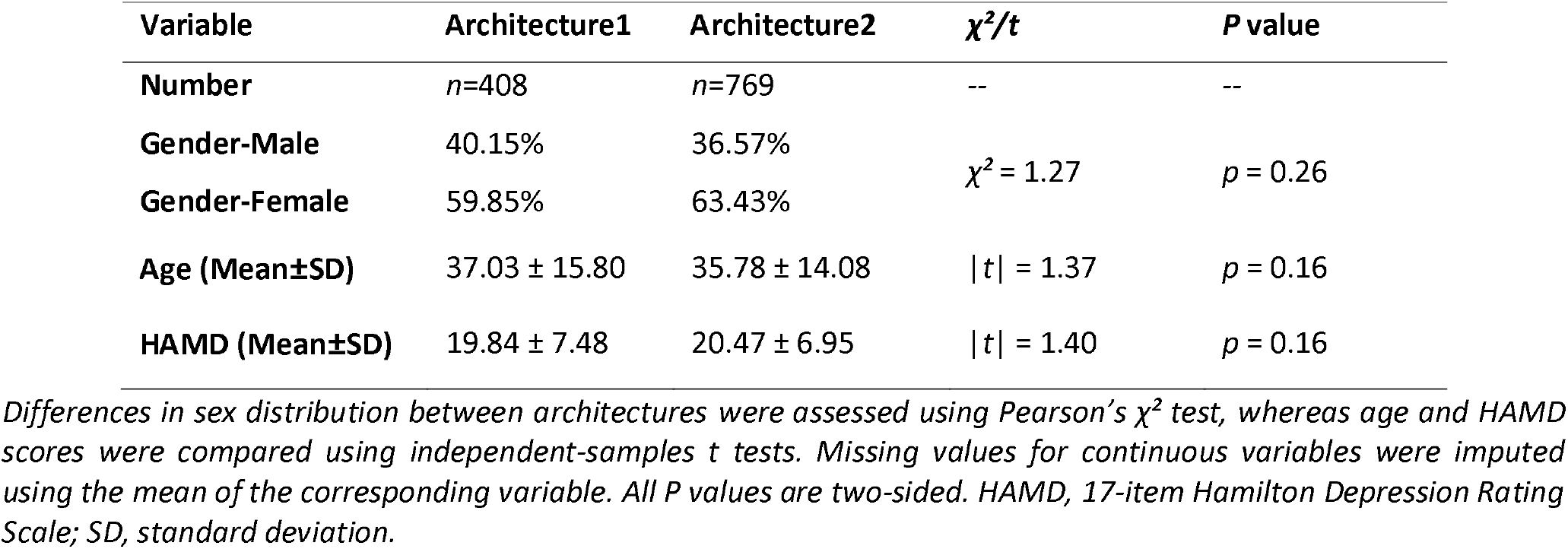
Demographic and clinical characteristics of the neurodynamic architectures in major depressive disorder.

At the global level, the two architectures showed contrasting state-preference profiles reflected by FO patterns (*Fig. 4A* and *Supplementary Table S5*). Relative to healthy controls, Architecture 1 showed preferential engagement of the ACS and VSS, together with reduced engagement of the MAS and DMS. In contrast, Architecture 2 showed the opposite pattern, with greater engagement of the MAS and DMS and reduced engagement of the ACS and VSS. The THS showed a smaller but significant FO difference between architectures, with higher FO in Architecture 1. The dominant state-preference contrast therefore followed an ACS/VSS–MAS/DMS axis, with THS contributing less strongly to the separation.

**Figure 4.**
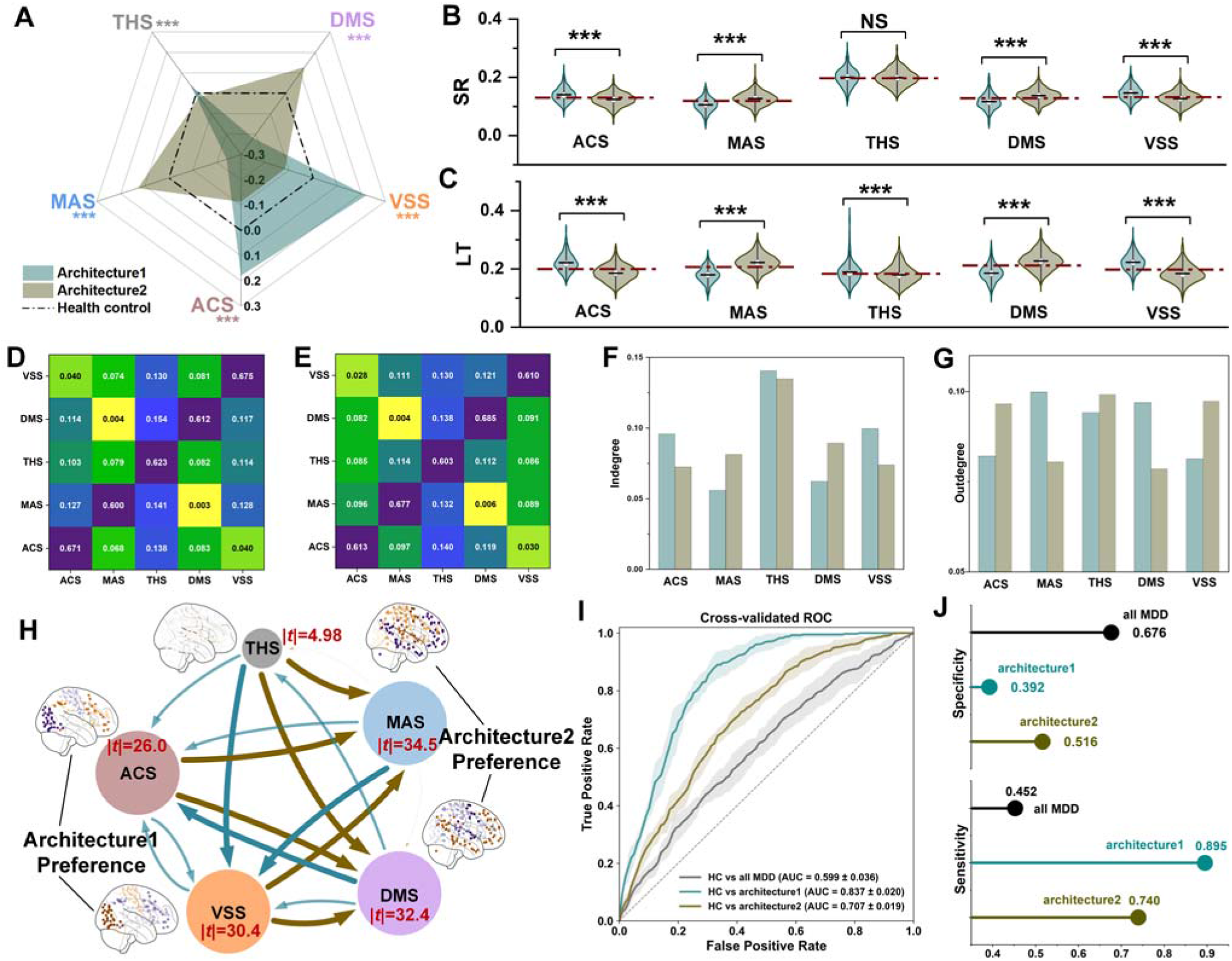
Neurodynamic architectures reveal divergent organizations of a shared brain-state repertoire in MDD. **(A)** State-preference profiles of the two neurodynamic architectures. Relative state preferences were characterized using FO across the five recurrent brain states. Values represent deviations relative to healthy controls, illustrating contrasting state-engagement patterns between the two neurodynamic architectures; **(B–C)** Architecture differences in temporal dynamics. Architecture-specific differences in SR and LT across the five recurrent brain states are shown. Red dashed lines indicate the corresponding mean values in HCs; **(D–E)** Transition probability matrix of neurodynamic architectures. TP matrices show state-transition organization in architecture1 and architecture2. Both architectures preserved a similar transition backbone while exhibiting architecture-specific transition preferences; **(F–G)** Degree centrality of state-specific TP. Indegree and outdegree derived from transition probability matrices were used to characterize architecture differences in state-specific transition involvement; **(H)** Differential transition network between neurodynamic architectures. The differential network represents between-architecture differences in fractional occupancy and transition probabilities rather than the absolute transition probabilities of either architecture. Node size represents the absolute *t* statistic from FO comparisons, edge width represents the absolute *t* statistic from transition probability comparisons, and edge color indicates the architecture showing higher transition probability; **(I–J)** Cross-validated separability from HCs. ROC curves illustrate the extent to which individuals in the pool ed MDD group, Architecture 1, and Architecture 2 could be distinguished from HCs using the same brain-state features. Sensitivity and specificity at the selected operating thresholds are shown in (J). Separability from HCs was evaluated using five-fold cross-validation; *P* < 0.05, P < 0.01, and \*\**P* < 0.001.

Despite these differences in state preference, both architectures largely preserved the canonical activation configurations defining each brain state. Across ACS, MAS, THS, DMS, and VSS, the average functional configurations of both architectures remained broadly consistent with the corresponding canonical state templates, with only modest state-specific regional differences (*Supplementary Material S2*). Both architectures therefore retained a common recurrent state space while differing primarily in how its constituent states were temporally deployed.

Beyond differences in state occupancy, the two architectures exhibited corresponding differences in persistence and switching across recurrent brain states (*Fig. 4B–C* and *Supplementary Table S5*). Architecture 1 showed higher switching rates within ACS and VSS but lower switching rates within MAS and DMS compared with Architecture 2, whereas THS demonstrated minimal architecture differences. Lifetime exhibited a complementary state-dependent pattern, with Architecture 1 showing prolonged persistence in ACS and VSS, while Architecture 2 showed longer persistence in MAS and DMS. Together with the FO differences, these findings indicate that architecture differentiation involved coordinated alterations in state occupancy, switching behavior, and temporal persistence rather than isolated changes in a single dynamic feature.

We further examined architecture-related differences in transition probabilities. Both architectures retained a broadly similar transition backbone characterized by prominent diagonal self-transitions and comparable overall state-transition organization. Within this shared backbone, architecture-specific transition preferences were observed. Architecture 1 showed stronger self-transitions and incoming transition involvement for ACS and VSS, whereas Architecture 2 showed corresponding preferences for MAS and DMS. Weighted indegree and outdegree further revealed complementary state-dependent patterns of transition involvement (*Fig. 4F–G*). THS maintained a relatively similar topological position across architectures, consistent with its shared role as a transition-dominant state. Differential transition-network analysis further revealed coordinated architecture differences in state occupancy and transition probabilities (*Fig. 4H* and *Supplementary Table S6*), with ACS- and VSS-related components showing greater expression in Architecture 1 and MAS- and DMS-related components showing greater expression in Architecture 2.

Finally, we evaluated whether resolving MDD according to neurodynamic architecture affected its differentiation from HCs using the same brain-state features (*Fig. 4I–J*). When individuals with MDD were considered as a single group, their cross-validated separability from HCs was limited (AUC = 0.599 ± 0.036). In contrast, architecture-resolved comparisons showed greater separability, with AUCs of 0.837 ± 0.020 for Architecture 1 and 0.707 ± 0.019 for Architecture 2. At the selected operating thresholds, Architecture 1 and Architecture 2 showed higher sensitivity than the pooled MDD group (0.895 and 0.740 versus 0.452), accompanied by lower specificity (0.392 and 0.516 versus 0.676). These findings indicate that neurodynamic stratification exposed brain-state patterns that were less apparent when MDD was treated as a single diagnostic group. However, the increased separability involved different sensitivity–specificity trade-offs and should be interpreted as greater differentiation of brain-state patterns rather than as uniformly improved diagnostic classification performance.

Together, these findings indicate that MDD heterogeneity can be characterized by two neurodynamic architectures arising from different temporal organizations of a shared recurrent brain-state repertoire. Architecture1 was characterized by preferential engagement and persistence of ACS/VSS states, whereas architecture2 showed preferential organization of MAS/DMS states. These divergent architectures were consistently expressed across state occupancy, switching dynamics, persistence, and transition organization.

### 2.5 Neurodynamic architectures contextualize brain–symptom associations and symptom-network patterns

Mean HAMD severity did not differ significantly between Architecture 1 and Architecture 2 (19.84 ± 7.48 versus 20.47 ± 6.95; *P* = 0.16; *Table 2*), indicating that the two architectures were not separated by overall symptom burden. We therefore examined architecture-specific brain–symptom associations and symptom network structures. Compared with the broadly distributed associations observed in the pooled MDD cohort (*Fig. 3F*), architecture-resolved analyses revealed distinct patterns linking recurrent brain state organization with depressive symptom severity (*Fig. 5A–B* and *Supplementary Table S7*).

**Figure 5.**
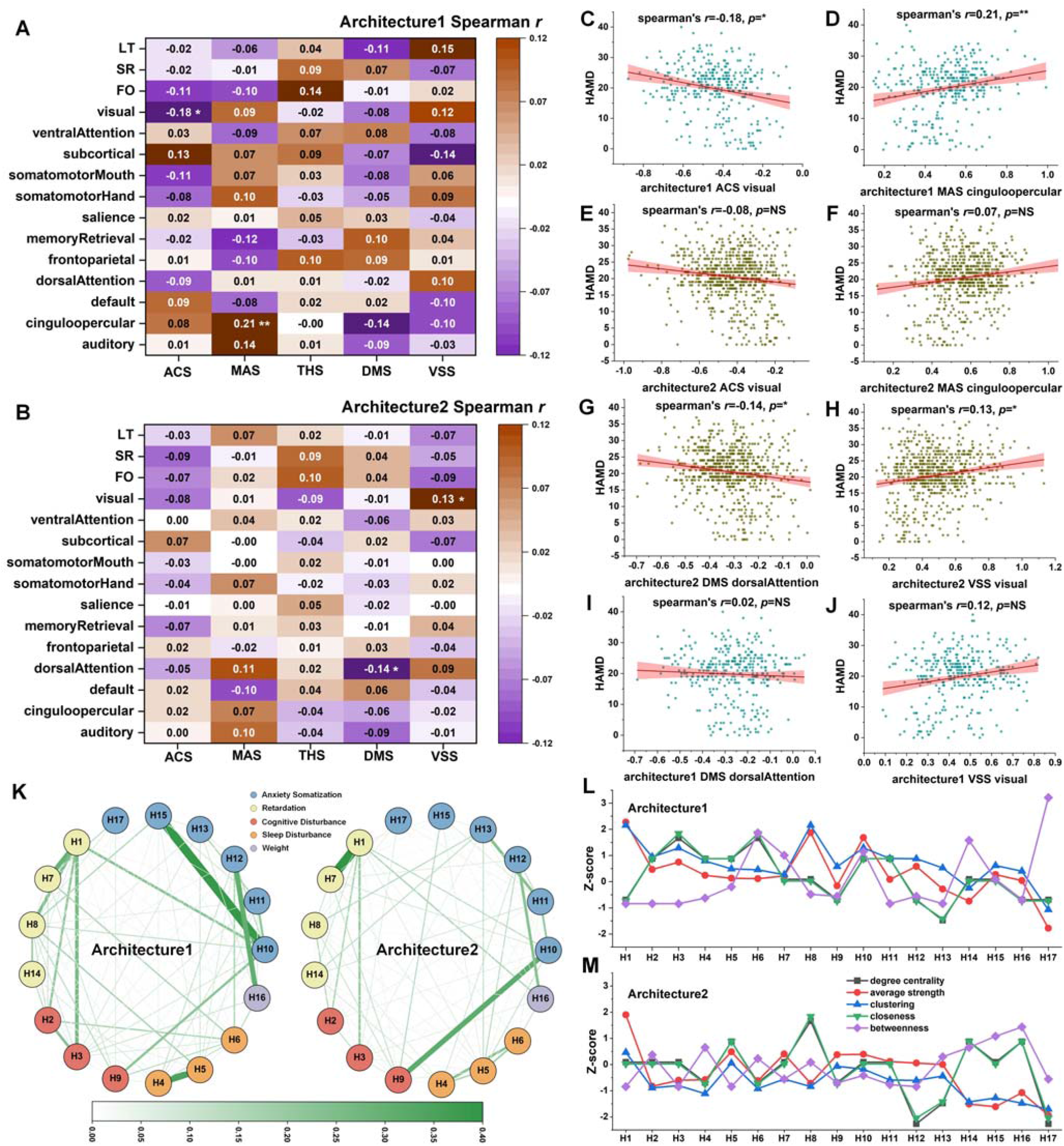
Brain–symptom associations and symptom-network structures across neurodynamic architectures. **(A–B)** architecture-specific associations between brain state features and depressive symptom severity. Heatmaps show Spearman correlation coefficients between HAMD scores and state-specific functional community features and dynamic features for (A) architecture1 and (B) architecture2; **(C–D)** Representative significant brain–symptom associations identified in architecture1. Scatter plots show associations between HAMD scores and visual-network activity in ACS (C) and cingulo-opercular activity in MAS (D), representing the strongest significant associations observed in architecture1; **(E–F)** Corresponding brain-state features shown in (C–D) plotted in architecture2 for comparison; **(G–H)** Representative significant brain–symptom associations identified in architecture2. Scatter plots show associations between HAMD scores and dorsal-attention activity in DMS (G) and visual-network activity in VSS (H), representing the strongest significant associations observed in architecture2; **(I–J)** Corresponding brain-state features shown in (G–H) plotted in architecture1 for comparison; **(K)** Symptom network structures of the two neurodynamic architectures estimated from HAMD items. Nodes represent individual HAMD symptoms, edges represent conditional associations between symptoms, and node colors indicate symptom-factor categories; **(L–M)** Centrality profiles of symptom networks for (L) architecture1 and (M) architecture2, summarizing topological properties of individual symptoms across network metrics; *\*P* < 0.05, *\*\*P* < 0.01, and *\*\*\*P* < 0.001. All statistical tests were corrected for multiple comparisons using FDR.

In Architecture 1, HAMD severity was significantly associated with lower visual-network activity in the ACS (*r* = -0.181) and higher cingulo-opercular activity in the MAS (*r* = 0.207; *Fig. 5C–D*). In Architecture 2, symptom severity was significantly associated with lower dorsal-attention activity in the DMS (*r* = -0.141) and higher visual-network activity in the VSS (*r* = 0.129; *Fig. 5G–H*). The corresponding associations of these brain-state features in another architecture showed weaker or non-significant effects (Fig. *5E–F* and *I–J*). Together, these findings indicate that depressive symptom severity was associated with different state-dependent neural features across neurodynamic architectures, suggesting that similar levels of symptom burden may be expressed through partially distinct patterns of brain-state organization.

Symptom networks were estimated separately for Architecture 1 and Architecture 2 using the 17 HAMD items (*Fig. 5K*). Bootstrap analyses supported the stability of the estimated network structures and centrality profiles across resamples (see *Methods*). The strongest edge within Architecture 1 connected psychic anxiety and hypochondriasis (H10-H15; weight = 0.388). Other comparatively prominent edges involved insomnia, somatic symptoms, anxiety-related symptoms, and work/activity impairment. Within Architecture 2, the strongest edge connected depressed mood and work/activity impairment (H1-H7; weight = 0.452), with other comparatively prominent edges involving anxiety, agitation, somatic symptoms, and sleep disturbance. The complete edge-weight distributions for both networks are provided in *Supplementary Table S8*.

Centrality measures were subsequently calculated for each symptom network (*Fig. 5L-M* and *Supplementary Table S8*). Depressed mood (H1) showed the highest average strength in both Architecture 1 and Architecture 2, providing a shared feature across the two networks. In Architecture 1, psychomotor retardation (H8) and psychic anxiety (H10) also showed relatively high average strength. Insight (H17) had the highest betweenness centrality, suicide (H3) had the highest degree centrality, and late insomnia (H6) had the highest closeness centrality. Within Architecture2, psychomotor retardation (H8) showed the highest degree and closeness centrality, whereas weight loss (H16) and hypochondriasis (H15) showed relatively high betweenness centrality. Together, these profiles characterized the prominent symptom relationships and node-level topology within each architecture.

Overall, these findings demonstrate that neurodynamic architectures were associated with distinct clinical architectures of depression. While depressed mood represented a shared central component across architectures, the organization of symptom relationships and their coupling with recurrent brain states differed between groups. Together with architecture-specific brain state organization, these findings suggest that similar levels of depressive symptoms may emerge from partially distinct neural and clinical configurations.

### 2.6 Transcriptomic organization distinguishes recurrent state families shared across neurodynamic architectures

To characterize the transcriptomic context of recurrent brain states, we linked architecture-state-specific regional activity patterns with regional gene-expression profiles derived from the Allen Human Brain Atlas (AHBA). Separate architecture–state-specific Partial Least Squares (PLS) models were constructed for each combination of neurodynamic architecture and brain state, resulting in ten transcriptomic models that characterized associations between brain-state patterns and regional gene-expression profiles (see *Methods*).

Across all architecture–state combinations, the first latent component (PLS1) consistently explained the largest proportion of imaging-related covariance among successive components (*Fig. 6A–B, Supplementary Table S9*). Permutation testing demonstrated significant PLS1 associations across architecture–state combinations, and regional activity patterns showed significant correlations with the corresponding PLS1-derived transcriptomic patterns (*Fig. 6C–D*; *Supplementary Table S10*), indicating that PLS1 captured the dominant component linking state-dependent neuroimaging patterns with gene-expression profiles.

**Figure 6.**
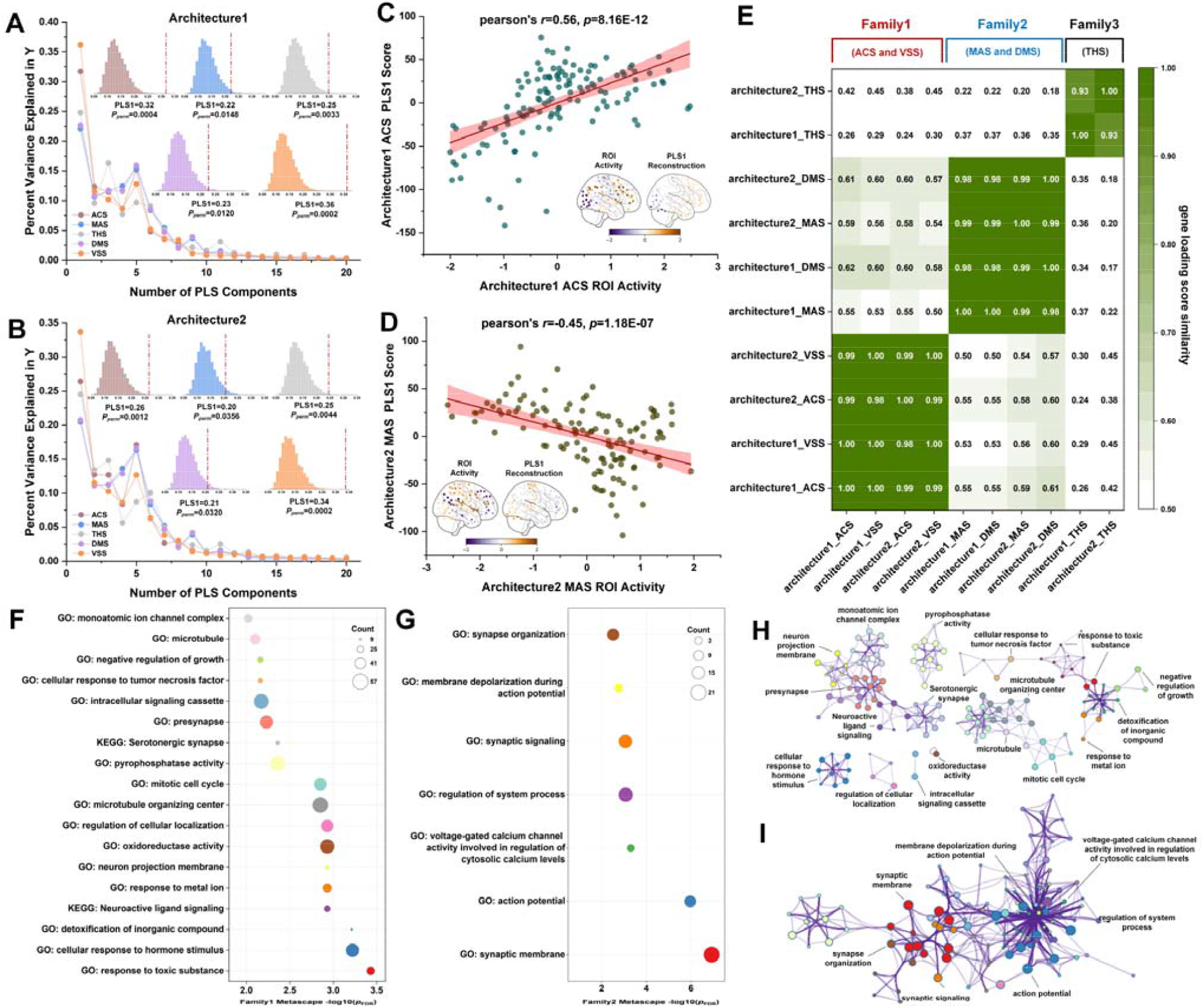
Transcriptomic organization distinguishes recurrent state families shared across neurodynamic architectures. **(A–B)** Variance explained by successive PLS components for architecture–state-specific transcriptomic models. Lines indicate the proportion of imaging-related covariance explained by each PLS component, and bars represent the corresponding cumulative variance explained. PLS1 accounted for the largest proportion of explained covariance across architecture–state combinations. Insets show permutation distributions of PLS1 performance, demonstrating the significance of the observed brain–transcriptome associations; **(C–D)** Representative spatial associations between state-specific neuroimaging patterns and PLS1-derived transcriptomic patterns. Scatter plots illustrate relationships between regional imaging patterns and PLS1 scores, with corresponding brain maps showing spatial distributions of imaging patterns and transcriptomic gradients; **(E)** Similarity structure of PLS1-derived transcriptomic patterns across architecture–state combinations. Heatmaps show pairwise similarity based on PLS1 scores and gene-loading profiles, identifying three transcriptomic state families: Family 1 (ACS and VSS), Family 2 (MAS and DMS), and Family 3 (THS); **(F–G)** Functional enrichment analyses of the major transcriptomic families. Family-specific gene sets were derived from PLS1-associated genes, and significantly enriched GO and KEGG terms are shown for Family 1 and Family 2; **(H–I)** PPI networks of the major transcriptomic families. Nodes represent genes included in the PPI networks, and edges indicate known protein–protein interactions, illustrating the interaction structures of genes associated with Family 1 and Family 2.

To examine how transcriptomic patterns were organized across recurrent states and architectures, we compared the similarity of PLS1 scores and gene-loading profiles across all architecture–state combinations (see *Methods*). Gene-loading similarity revealed a structured organization, with higher similarity within specific groups of recurrent states than between groups (*Fig. 6E*); PLS1 score similarity showed a broadly concordant organization (*Supplementary Fig. S11*). These patterns defined three transcriptomic state families shared across architectures (*Supplementary Material S3*). Family 1 consisted of ACS and VSS from both neurodynamic architectures and showed highly consistent gene-loading profiles (within-family average similarity = 0.99). Family 2 consisted of MAS and DMS from both architectures and demonstrated similarly high within-family similarity (average similarity = 0.99). Family 3 consisted of THS across both architectures and showed a distinct but internally consistent transcriptomic profile (average similarity = 0.97). Notably, this transcriptomic organization mirrored the neurodynamic architecture identified at the behavioral level, ACS/VSS represented the state configuration preferentially engaged by architecture1, whereas MAS/DMS represented the configuration preferentially engaged by architecture2. These findings suggest that different neurodynamic architectures are embedded within partially distinct molecular contexts while relying on a shared repertoire of recurrent brain states.

We next characterized the biological functions associated with the major transcriptomic families using GO and KEGG enrichment analyses applied to family-specific gene sets derived from PLS1 bootstrap ratios (see *Methods*). The family 1 gene set was primarily enriched in pathways related to cellular regulation, stress-response processes, and structural organization, including response to toxic substances, detoxification of inorganic compounds, cellular response to hormone stimuli, regulation of cellular localization, mitotic cell cycle, intracellular signaling cascades, microtubule organization, neuroactive ligand signaling, and serotonergic synapse pathways (*Fig. 6F*). In contrast, the Family 2 gene set was predominantly enriched in neuronal and synaptic processes, including synaptic membrane organization, synaptic signaling, action potential generation, membrane depolarization during action potential, synapse organization, regulation of neurotransmitter secretion, regulation of synaptic plasticity, and voltage-gated calcium-channel activity (*Fig. 6G*). No significantly enriched pathways survived multiple-comparison correction for Family 3 (*Supplementary Table S11*).

Protein–protein interaction (PPI) analyses characterized the interaction structures of genes associated with the two major transcriptomic families (*Fig. 6H–I*). Family 1 exhibited interconnected modules involving cellular regulation, intracellular signaling, stress-response pathways, and structural organization, whereas Family 2 showed densely connected modules centered on synaptic organization, neuronal communication, neurotransmission, and calcium-dependent signaling. Together, these findings distinguish the transcriptomic contexts of recurrent state families shared across neurodynamic architectures and provide a molecular background for the states preferentially deployed along the principal temporal organization within MDD.

### 2.7 Neurodynamic organization is recovered in an independent MDD cohort

To assess the cross-cohort consistency of architecture-related temporal organization, we evaluated an independent validation cohort comprising 65 individuals with MDD using the same brain-state framework (see *Methods*). Demographic and clinical characteristics of the validation cohort are summarized in *Table 3*. The discovery-derived HMM was applied directly to the validation data to decode participant-specific state sequences, the same 15 dynamic features, comprising FO, SR, and LT for each state, were then extracted. Validation participants were then independently re-clustered using the same dynamic-feature representation and hierarchical clustering strategy, without reference to their architecture assignments in the discovery cohort.

**Table 3.** Demographic and clinical characteristics of the neurodynamic architectures in validation dataset.

| Variable | Total | Architecture1 | Architecture2 | $\chi^2/t$ | P value |
| --- | --- | --- | --- | --- | --- |
| **Number** | *n* = 65 | *n* = 18 | *n* = 47 | -- | -- |
| **Gender-Male** | 36.92% | 33.33% | 38.30% | $\chi^2 = 0.007$ | *p* = 0.93 |
| **Gender-Female** | 63.08% | 66.67% | 61.70% |  |  |
| **Age (Mean±SD)** | 26.00 ± 7.48 | 24.39 ± 7.49 | 26.62 ± 7.46 | \|*t*\| = 1.08 | *p* = 0.29 |
| **HAMD (Mean±SD)** | 29.12 ± 6.22 | 29.28 ± 6.05 | 29.06 ± 6.35 | \|*t*\| = 0.12 | *p* = 0.90 |

*Differences in sex distribution between architectures were assessed using Pearson's $\chi^2$ test, whereas age and HAMD scores were compared using independent-samples *t* tests. Missing values for continuous variables were imputed using the mean of the corresponding variable. All *P* values are two-sided. HAMD, 17-item Hamilton Depression Rating Scale; SD, standard deviation.*

Independent re-clustering delineated two validation groups with contrasting state-preference patterns broadly consistent with those identified in the discovery cohort. One group showed preferential engagement of ACS and VSS, accompanied by reduced engagement of MAS and DMS, whereas the other showed the complementary pattern, characterized by greater engagement of MAS and DMS and reduced engagement of ACS and VSS (*Supplementary Figures S12-S13*). These state-preference patterns reproduced the principal distinction between the two discovery architectures, and corresponding SR and LT differences broadly followed the same organization (*Supplementary Figure S14*). Although the magnitude of individual feature differences varied between cohorts, the directionality of the principal architecture-related differences was broadly preserved.

As a complementary transfer-based analysis, the architecture centroids derived from the discovery cohort were used to assign validation participants directly to Architecture 1 or Architecture 2. After alignment of the architecture labels, the centroid-based assignments agreed with the independently re-clustered solution for 42 of 65 participants (*Supplementary Table S17*).

Together, these analyses indicate that the principal ACS/VSS–MAS/DMS organizational axis was recoverable in an independent cohort at the group-pattern level, whereas the moderate agreement between transfer-based and independently re-clustered assignments suggests that individual-level architecture membership remained partly sensitive to the assignment strategy.

## 3. Discussion

MDD is characterized by substantial clinical and biological heterogeneity, yet the dimensions of brain organization underlying this variability remain poorly understood. Here, we show that individual differences in the temporal organization of recurrent brain states represent an additional dimension of neurobiological heterogeneity in MDD. By distinguishing recurrent activity configurations from their temporal organization, we identified a shared repertoire of five brain states with distinct spatial and temporal properties across individuals with MDD and healthy controls. At the diagnostic-group level, MDD-related alterations were distributed across multiple states, functional systems, and temporal properties rather than converging on a single common pattern. Neurodynamic stratification further delineated two prototype architectures that were distinguished not by different state repertoires or the emergence of novel activity configurations, but by coordinated differences in state occupancy, persistence, switching, together with corresponding differences in transition patterns. The two architectures showed different within-group brain–symptom association patterns and symptom-network structures, while transcriptomic analyses organized architecture–state imaging patterns into state-related molecular families shared across architectures. Importantly, contrasting group-level patterns characterized by preferential deployment of ACS/VSS or MAS/DMS states were also recovered in an independent MDD cohort. Together, these findings suggest that MDD heterogeneity may arise not only from variation in neural abnormalities, but also from different temporal organizations of a shared recurrent brain-state repertoire.

### 3.1 Brain-state dynamics provide a complementary dimension for understanding MDD heterogeneity

MDD has traditionally been investigated using static or time-averaged measures of brain organization, which have revealed spatially distributed alterations across large-scale functional systems. However, such representations provide limited information about how recurrent activity configurations are deployed and reorganized over time. By resolving spontaneous brain activity into five recurrent states, our analysis distinguished two complementary aspects of brain organization: spatial configurations, which describe the activity pattern defining each state, and temporal properties, which describe how individuals occupy, maintain, switch among, and revisit these states [32]. This distinction makes it possible to examine whether individuals who share a common state repertoire nevertheless differ in how that repertoire is organized over time.

The identified repertoire comprised four states with differentiated system-level activity biases and a transition-oriented state with a comparatively balanced activity profile. ACS and DMS preferentially weighted associative-control and default-memory systems, respectively, whereas MAS and VSS preferentially weighted multimodal attention–salience and visual–sensorimotor systems. Together, these states represented recurrent configurations spanning different combinations of large-scale functional systems. THS was distinguished less by a strongly specialized spatial profile than by its temporal position within the repertoire: it was frequently occupied, briefly maintained, and received strong incoming transitions. These properties support its operational interpretation as a recurrent intermediate configuration within the observed transition structure [33], broadly consistent with accounts of metastability in which brain activity moves among differentiated configurations through transient intermediate patterns [34, 35]. They do not, however, establish that THS causally coordinates transitions or maintains neural flexibility.

At the diagnostic-group level, MDD-related differences did not converge on a single state or a uniform temporal shift. Instead, they were distributed across multiple state–system combinations and temporal properties, rather than reflecting a uniform disruption of the state repertoire. Case–control comparisons therefore capture population-level alterations but may compress differently organized individual patterns within the diagnostic group [36]. Examining the temporal organization of recurrent states consequently adds an important level of analysis: beyond identifying where MDD-related alterations occur, it provides a framework for determining how a common repertoire is differentially deployed across individuals with MDD.

### 3.2 Prototype neurodynamic architectures reflect divergent deployment of a shared brain-state repertoire

The two neurodynamic architectures identified in this study were not defined by different brain-state repertoires or architecture-specific activity configurations. Both expressed the same spatial profiles of the five recurrent states and a broadly similar transition backbone. Their defining differences comprised coordinated variation in state occupancy, persistence, and switching, while subsequent transition analyses revealed corresponding differences in transition preferences. The central distinction between the architectures therefore lay not in which states were expressed, but in how the shared states were deployed and coordinated over time.

Architecture 1 showed preferential engagement and greater persistence of ACS and VSS, whereas Architecture 2 preferentially engaged and maintained MAS and DMS. Corresponding differences were observed in switching and transition patterns, while THS showed comparatively limited differentiation between the two architectures. The architectures thus primarily reflected different relative weightings of the four activity-biased states against a largely preserved repertoire and transition structure. Rather than defining qualitatively distinct pathological systems, they summarize recurring prototype patterns within broader neurodynamic variation in MDD. Their value therefore lies in representing structured organizational tendencies, rather than imposing categorical boundaries between patients, consistent with dimensional accounts of psychiatric heterogeneity [15].

The contrasting state preferences of the two architectures may also help explain why some neurodynamic differences are attenuated when individuals with MDD are pooled into a single diagnostic group. When ACS/VSS-preferring and MAS/DMS-preferring individuals are combined, differently directed patterns may partially offset one another, attenuating group-level effects. Consistent with this possibility, post hoc classification analyses showed clearer separation from HCs for each architecture than for the pooled MDD group. However, because the architectures were derived from brain-state dynamics and the classifiers incorporated overlapping brain-state information, these analyses do not provide an independent test of the validity of the stratification. The architecture-specific AUCs should therefore be interpreted as a descriptive illustration of the patterns exposed after stratification, rather than evidence of improved diagnostic performance. Together, these findings suggest that diagnostic averaging may obscure structured differences in how a shared brain-state repertoire is temporally organized within MDD.

### 3.3 Clinical patterns are organized within different neurodynamic contexts

The two neurodynamic architectures showed comparable overall depressive severity, indicating that their differentiation did not simply recapitulate differences in symptom burden. In the pooled MDD cohort, associations between brain-state features and depressive severity were relatively modest and broadly distributed. Following neurodynamic stratification, architecture-resolved analyses revealed more differentiated state-dependent relationships between brain organization and symptom severity. In Architecture 1, the most prominent associations involved visual-system activity in ACS and cingulo-opercular activity in MAS, whereas those in Architecture 2 involved dorsal-attention activity in DMS and visual-system activity in VSS. This distribution is consistent with the possibility that pooling individuals with different neurodynamic organizations attenuates context-dependent brain–symptom relationships [37]. Importantly, stratification did not uniformly strengthen all brain–symptom associations; rather, it revealed architecture-specific relationships that were diluted in the pooled cohort. This suggests that similar depressive severity may be associated with different configurations of dynamic brain organization rather than from a single common neural correlate.

Symptom-network analyses provided a complementary view of clinical organization within each neurodynamic context. Although depressed mood was a highly connected symptom across both architectures, indicating a shared prominent feature of their symptom networks, the surrounding patterns of symptom relationships were descriptively different. Architecture 1 showed broader coupling among anxiety, somatic symptoms, sleep disturbance, and functional impairment, whereas Architecture 2 exhibited a more concentrated organization involving depressed mood, functional impairment, anxiety, agitation, and somatic symptoms. These findings suggest that comparable symptom severity may reflect different configurations of symptom interactions.

Together, the convergence between neurodynamic organization, brain–symptom coupling, and symptom-network architecture indicates that MDD heterogeneity is expressed across multiple organizational levels. Neurodynamic architectures therefore represent distinct biological contexts through which depressive symptoms are organized rather than separate psychiatric disorders. This perspective may also provide a conceptual basis for future neurodynamic-informed intervention strategies. Rather than implying distinct treatment categories, these architectures may reflect different patterns of brain-state regulation that could influence how individuals respond to interventions targeting cognitive flexibility, self-referential processing, or adaptive network reconfiguration. Future longitudinal studies incorporating treatment-response data will be required to determine whether neurodynamic profiles can predict therapeutic outcomes or guide personalized intervention strategies.

### 3.4 Transcriptomic organization provides molecular context for recurrent state families across neurodynamic architectures

Connectome-transcriptome analyses revealed a structured similarity organization among the PLS-derived gene-expression signatures associated with architecture–state imaging patterns. Across architecture–state combinations, PLS-derived signatures converged into three transcriptomic families, organized primarily according to recurrent state grouping rather than architecture membership. ACS and VSS formed one family, MAS and DMS formed a second family, and THS constituted a third, with each family including the corresponding states from both neurodynamic architectures. This cross-architecture convergence indicates that transcriptomic similarity was more closely aligned with shared state families than with the patient grouping itself.

The organization of these families nevertheless corresponded to the state preferences distinguishing the two architectures, ACS and VSS were preferentially deployed in Architecture 1, whereas MAS and DMS were preferentially deployed in Architecture 2. Importantly, because both transcriptomic families were represented in both architectures, these findings do not indicate that the two patient groups possess distinct molecular profiles. Rather, they suggest that the architectures differ in the temporal deployment of recurrent states embedded within partially differentiated transcriptomic contexts. This correspondence therefore extends the central organizational principle of the study: the architectures share both the recurrent state repertoire and its associated transcriptomic organization, while differing in how these common states are deployed over time.

Functional enrichment analyses further supported this interpretation, with ACS/VSS-associated signatures showing enrichment for cellular regulation, stress-response processes, and intracellular signaling, whereas MAS/DMS-associated signatures were characterized by pathways related to synaptic organization, neuronal signaling, neurotransmission, and membrane excitability. However, these findings should not be interpreted as direct molecular mechanisms driving neurodynamic architectures. AHBA-based analyses primarily capture spatial correspondence between regional gene-expression patterns and neuroimaging phenotypes rather than individual-level molecular determinants [38]. The absence of strong pathway enrichment for THS may reflect its transition-oriented role. Unlike specialized states, THS may depend on distributed biological processes rather than a single dominant molecular program. This interpretation is consistent with its proposed role as a coordination state within the brain-state repertoire.

Together, these findings provide a molecular context for recurrent state families shared across neurodynamic architectures. The transcriptomic organization was associated primarily with common state configurations, whereas architecture-level heterogeneity was expressed through differences in how these states were temporally deployed.

### 3.5 Limitations and future directions

Several limitations should be considered. First, although the two-cluster solution showed the highest silhouette coefficient, its absolute separation was modest, and resampling showed moderate global partition agreement despite relatively high subject-level stability. The architectures should therefore be interpreted as organizational prototypes rather than definitive biological subtypes. Moreover, the independent cohort was relatively small, the discovery-derived HMM was transferred without re-estimating the state repertoire, and agreement between the two assignment strategies was moderate. The validation therefore supports recovery of the group-level organizational patterns more strongly than stable transfer of individual architecture membership.

Second, the multisite discovery dataset contained variation in acquisition protocols, temporal resolution, and clinical composition. Standardized preprocessing and site-wise characterization improved comparability but could not eliminate these influences on brain-state dynamics. Further validation using harmonized datasets and explicit modelling of site effects is needed.

Third, the clinical analyses were cross-sectional, and the architecture-resolved correlations and symptom networks were not directly compared between groups. These results therefore provide descriptive clinical context rather than evidence for distinct clinical phenotypes. Incomplete information on medication exposure, illness duration, episode history, and comorbidity may also have contributed to neurodynamic variation. Longitudinal studies are needed to determine whether the architectures remain stable, change with illness progression, or relate to treatment response.

Fourth, AHBA-based transcriptomic mapping reflects spatial correspondence between group-level neuroimaging patterns and gene expression in healthy post-mortem tissue. It does not establish patient-specific molecular differences or molecular mechanisms underlying state dynamics.

Finally, future studies integrating longitudinal imaging, treatment-response data, multimodal biological measurements, and computational modelling may further clarify the significance of neurodynamic architectures. Advances in brain network analysis may improve the characterization of multiscale and time-varying brain organization [39-41], whereas network control theory may provide a framework for examining how state transitions are constrained or altered by perturbation [42-45].

## 4. Conclusion

In conclusion, this study demonstrates that the temporal organization of recurrent brain states represents a biologically meaningful dimension of heterogeneity in MDD. By characterizing both the functional configurations and dynamic properties of a shared brain-state repertoire, we identified distinct neurodynamic architectures that differentiated individuals with MDD according to how recurrent states were engaged, maintained, switched, and transitioned over time. These architectures were associated with architecture-specific brain–symptom relationships, symptom-network organizations, and transcriptomic profiles, and were reproducible in an independent cohort. Our findings suggest that heterogeneity in MDD is not solely expressed through differences in neural abnormalities, but also through individual differences in the dynamic coordination of distributed brain systems. This neurodynamic perspective provides a framework for moving beyond diagnostic averaging toward a more mechanistic understanding of biological variability in depression.

## 5. Methods

### 5.1 Participants

Two independent cohorts were included in the present study, a large-scale discovery cohort from the REST-meta-MDD consortium and an independent validation cohort from the Affiliated Brain Hospital of Guangzhou Medical University.

The discovery cohort consisted of 2,225 participants, including 1,177 individuals with MDD and 1,048 HCs, from the REST-meta-MDD dataset (https://rfmri.org/REST-meta-MDD-V1) [46, 47]. The sample size was determined by the availability of eligible rs-fMRI data after quality control. Demographic and clinical information was available for subsets of participants and was included when available. Missing values were only imputed by average for descriptive comparisons of demographic and clinical variables and were not used in imaging analyses. Depressive symptoms were assessed using the 17-item Hamilton Depression Rating Scale (HAMD-17) [48]. An independent validation cohort consisting of 65 individuals with MDD was recruited from Affiliated Brain Hospital of Guangzhou Medical University, with no overlap with participants in the discovery cohort. Available demographic and clinical characteristics of the discovery and validation cohorts are summarized in *Table 1* and *Table 3*, respectively.

All participants provided written informed consent before participation. The original studies contributing to the REST-meta-MDD consortium and the validation cohort [*ethic approval number: 2021-008*] were approved by their respective institutional review boards or ethics committees and were conducted in accordance with the Declaration of Helsinki.

### 5.2 MRI acquisition and preprocessing

Resting-state fMRI (rs-fMRI) data for the discovery cohort were obtained from multiple sites participating in the REST-meta-MDD consortium [46]. The consortium implemented standardized imaging acquisition and preprocessing procedures to improve cross-site comparability across participating datasets. Detailed information regarding scanner vendors, acquisition parameters, and site-specific quality control procedures is available from the REST-meta-MDD database (https://rfmri.org/REST-meta-MDD-V1). Across sites, imaging data were acquired using 3.0 T MRI scanners with gradient-echo echo-planar imaging sequences using comparable acquisition parameters across sites [46].

The rs-fMRI data were preprocessed using the standardized preprocessing pipeline established by the REST-meta-MDD consortium [46, 47]. Following preprocessing, regional time series were extracted using the Power functional atlas [49]. Quality control was performed on the extracted ROI -level time series before subsequent brain-state analyses. Participants with invalid ROI time-series matrices, including all-zero rows, all-zero columns, missing values, or missing ROI signals, were excluded. After quality control, 2,225 participants mentioned in the previous section were retained as the discovery cohort for subsequent analyses.

For the independent validation cohort, resting-state fMRI data were acquired using an independent imaging protocol. Functional images consisted of 64 × 64 × 33 voxels with 240 time points (TR = 1,000 ms). The validation data were preprocessed using the same preprocessing strategy and Power functional atlas as the discovery cohort to ensure comparability across cohorts.

### 5.3 Hidden Markov modeling of brain states and dynamic features

Resting-state brain activity was modeled using a Gaussian Hidden Markov Model (HMM) to identify recurrent latent brain states underlying large-scale functional organization [50-52]. HMM assumes that observed multivariate neural activity arises from a sequence of unobserved states that evolve according to a first-order Markov process. Let *x*_t_ denote the regional activity vector at time point *t*, and *z*_t_ denote the corresponding latent state. Conditional on each state, neural activity was modeled using a multivariate Gaussian distribution [53]:

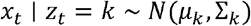

where *μ*_k_ represents the state-specific spatial activity pattern and Σ_k_represents the covariance structure of each state. The model estimated posterior state probabilities across time, providing a probabilistic representation of state expression, together with the most probable state sequence obtained from decoding.

HMM estimation was performed using the Python GLHMM toolbox (https://glhmm.readthedocs.io/en/latest/) [50]. Individual regional time series were concatenated across participants before model fitting. Candidate models with different numbers of states (K = 2–15) were evaluated to identify the most appropriate model order. For each candidate K, the model was independently fitted across 10 runs with different random seeds. Within each run, multiple initializations were performed using the default GLHMM procedure, and the solution with the best initial objective value was retained for subsequent variational expectation–maximization optimization. The final number of states was selected based on complementary criteria, including variational free energy (FE), Akaike information criterion (AIC), Bayesian information criterion (BIC), and across-run stability of inferred state representations. The five-state solution was selected as the final model and used for all subsequent analyses. Detailed *K* number selection procedures and results are provided in *Supplementary Material S1*.

The inferred brain states were characterized using both spatial and temporal properties. Each inferred state was represented by a 264-dimensional state-specific mean activity pattern (*μ*_k_), reflecting the spatial configuration of large-scale brain activity associated with that state. For functional interpretation, ROI-level state patterns were aggregated according to the community assignments provided by the Power atlas, and the mean activity of all ROIs within each community was calculated to generate community-level representations of each brain state. Among the 14 communities defined in the Power functional atlas [49], 12 cortical and subcortical communities were retained for subsequent interpretation, whereas the “None” and “Cerebellar” communities were excluded. Individual-level temporal dynamics were quantified using fractional occupancy (FO), lifetime (LT), switching rate (SR), and transition probability (TP). FO was defined as the mean posterior probability of each state across all time points, representing the estimated occupancy of each state. LT was calculated from the decoded state sequence as the mean duration of consecutive visits to each state and subsequently normalized within individuals. SR was defined as the mean absolute change in posterior state probability between adjacent time points, reflecting temporal variability in state expression. TP was calculated as the probability of transitioning from state *i*to state *j*, generating a directed transition matrix describing the temporal organization of the brain-state repertoire.

### 5.4 Identification of neurodynamic architectures in major depressive disorder

Brain state-based architecture identification was performed exclusively within the MDD cohort. For each participant, a candidate feature vector was constructed to characterize individual variability in both functional configuration and temporal organization of recurrent brain states. State-specific functional features consisted of the mean activity of 12 large-scale functional communities within each of the five brain states, resulting in 60 functional features. Dynamic features included FO, SR, and LT for each state, yielding 15 temporal features. Thus, each participant was represented by a 75-dimensional candidate feature space integrating functional and dynamic properties of brain-state organization.

All candidate features were standardized across participants using z-score normalization. Sparse principal component analysis (sPCA) [54] was applied as an exploratory approach to characterize the dominant dimensions of inter-individual variation within the candidate feature space. Importantly, sPCA was not used as a clustering algorithm or as a criterion for selecting architecture assignments, but rather to characterize the major sources of variability underlying brain-state heterogeneity. Five sparse components were extracted, and the leading component showed predominant contributions from dynamic features, indicating that temporal organization represented a major source of inter-individual variation in MDD. Given the objective of identifying neurodynamic architectures, the complete set of dynamic features (FO, SR, and LT across five states) was retained, rather than selecting only features associated with specific sparse components, as the prespecified feature space for subsequent architecture identification. This strategy preserved the multidimensional organization of brain-state dynamics without relying on individual feature selection. Detailed sPCA parameters and component loadings are provided in *Supplementary Material S2*.

Hierarchical clustering was subsequently performed using the retained dynamic feature space. Dynamic features were standardized using z-score normalization before clustering. Pairwise Euclidean distances were calculated, and agglomerative hierarchical clustering was performed using Ward’s linkage criterion. Candidate clustering solutions ranging from K = 2 to K = 9 were evaluated by cutting the hierarchical tree at different levels without imposing predefined architecture labels. The optimal cluster number was determined based on the average silhouette coefficient, and the resulting solution was used for subsequent architectures characterization.

The stability of the clustering solution was evaluated using repeated subsampling. In each of 1,000 iterations, 80% of MDD participants were randomly sampled without replacement, and hierarchical clustering was repeated using the same dynamic feature set, z-score standardization, Euclidean distance, and Ward’s linkage criterion. Global partition stability was assessed by comparing each subsampled solution with the full-sample two-cluster solution using the adjusted Rand index (ARI), normalized mutual information (NMI), and assignment-matching rate. Subject-level stability was quantified as the proportion of iterations in which each participant retained the same architecture assignment among all iterations in which that participant was included. Pairwise co-clustering stability was further evaluated using a consensus matrix, in which each element represented the proportion of co-sampled iterations in which a given pair of participants was assigned to the same architecture. Together, these analyses assessed the reproducibility of the two-architecture solution at the levels of overall partition structure, individual architecture assignment, and pairwise clustering consistency. Detailed results are provided in the *Supplementary Material S2*.

Finally, to characterize site-level composition and potential site-related heterogeneity, the final architecture assignments were summarized separately across contributing sites, together with sex, age, and HAMD scores (Supplementary Table S16).

### 5.5 Discriminative evaluation of MDD neurodynamic architectures

Following architecture identification, the final cluster labels were used to characterize architecture-specific patterns of brain-state organization. Individual-level state features and transition measures derived from the HMM analysis were summarized separately within each architecture, and architecture differences were evaluated using the statistical procedures described below. Group-averaged transition matrices and corresponding node-level transition metrics were calculated for each architecture. A differential transition network was visualized by mapping between-architecture test statistics of state occupancy onto network nodes and test statistics of directed transition probabilities onto network edges, thereby representing the magnitude and direction of architecture differences in state occupancy and transition organization (*Fig. 4H*).

To evaluate whether neurodynamic stratification resulted in more distinguishable brain-state organization relative to healthy controls, support vector machine (SVM) classifiers [54] were constructed for three binary classification tasks, HC versus all MDD participants, HC versus architecture1, and HC versus architecture2. Classification was performed independently from architecture identification, using the full 75-dimensional brain-state feature representation integrating functional and dynamic features. Missing values were imputed using the mean value of the corresponding feature, and all preprocessing steps, including feature normalization, were performed within each training fold to prevent information leakage. SVM models with radial basis function kernels and probabilistic output were implemented using identical settings across classification tasks.

Model performance was evaluated using stratified five-fold cross-validation with shuffled data partitions and a fixed random seed. In each fold, the model was trained on four-fifths of participants and evaluated on the held-out fifth. Receiver operating characteristic (ROC) curves and areas under the curve (AUCs) were calculated from predicted class probabilities. Mean AUC and standard deviation were calculated across five folds. Out-of-fold predictions were further generated to ensure that each participant was evaluated only when serving as a held-out observation. These predictions were used to calculate accuracy, precision, sensitivity, specificity, F1 score, negative predictive value, and cross-validated confusion matrices. Sensitivity was defined as the proportion of positive cases correctly identified, whereas specificity represented the proportion of negative cases correctly classified. Corresponding confusion matrices are provided in *Supplementary Figure S7*.

### 5.6 Symptom network construction and analyses

Architecture-specific symptom networks were constructed to characterize the organization of depressive symptoms associated with different neurodynamic architectures. Network analyses were performed separately within architecture1 and architecture2 using item-level scores from the HAMD-17. Participants were included when individual HAMD item scores were available and the total HAMD score was greater than zero. When necessary, missing values in individual HAMD items were handled using architecture-specific mean imputation. The resulting symptom matrices were standardized using z-score normalization before network estimation. Each HAMD item was treated as an individual symptom node.

We employed the Graphical LASSO (Glasso) [55] method to estimate architecture1-specific depressive symptom networks based on the 17 HAMD items. This approach estimates a Gaussian Graphical Model (GGM) by applying regularization through the Least Absolute Shrinkage and Selection Operator (LASSO), resulting in a sparse inverse covariance matrix [56]. The LASSO penalty (L1 penalty) eliminates potential bias to create a sparse network, a method widely used in psychological research. The mathematical formulation for the estimator is as follows [57]:

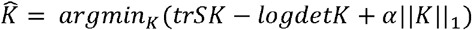

Where *K* is the sparse inverse covariance matrix to be estimated; *S* is the covariance matrix of the sample; and ∥*K*∥_1_ is the sum of the absolute values of the off-diagonal coefficients of *K. α* is the sparsity parameter. To improve analytical accuracy, we are setting the diagonal elements to zero. This approach focuses solely on the association strength between different nodes, disregarding self-association. In the constructed network, an edge was defined between any node pair *(i, j*) if the corresponding element in the matrix was non-zero, with the edge weight determined by the matrix value. The Graphical LASSO estimator was implemented using the scikit-learn library in Python [58].

To facilitate interpretation of symptom-network organization, HAMD items were further grouped into five symptom-factor categories based on the established HAMD-17 factor structure, including Anxiety/Somatization (H10, H11, H12, H13, H15, H17), Cognitive Disturbance (H2, H3, H9), Retardation (H1, H7, H8, H14), Sleep Disturbance (H4, H5, H6), and Weight (H16). These factor assignments were used for visualization of symptom communities and interpretation of network patterns, but were not involved in network estimation.

Network topology was characterized using node strength, degree centrality, closeness centrality, and betweenness centrality [59]. Strength quantified the overall connectivity of each symptom by summing the absolute weights of its connected edges. Degree centrality represented the number of direct symptom connections, whereas closeness and betweenness centrality characterized the relative position of each symptom within the overall network structure.

To evaluate network stability, we performed non-parametric bootstrapping [60]. We randomly resampled 90% of the full dataset and re-estimated the network structure [61, 62]. This process was repeated 1,000 times to obtain distributions, 95% confidence intervals, and mean values for each metric (*Supplementary Figure S9*).

### 5.7 Transcription-neuroimaging association analysis and gene enrichment

Brain-wide gene-expression data were obtained from the Allen Human Brain Atlas (AHBA; http://human.brain-map.org) [63] and preprocessed using the *abagen* toolbox (https://abagen.readthedocs.io) [64]. Given the limited sampling of the right hemisphere in the AHBA dataset, subsequent analyses were restricted to the left hemisphere of the Power264 atlas, yielding expression profiles for 15,632 genes across 125 cortical regions.

For each architecture-state combination, partial least squares (PLS) regression was performed to identify associations between regional neuroimaging abnormalities and transcriptional profiles [65]. Regional gene-expression levels were treated as predictor variables, whereas ROI activity values were treated as response variables. Prior to analysis, both gene-expression data and imaging phenotypes were standardized using z-score normalization. The first latent component (PLS1), which explained the greatest proportion of imaging-related variance, was retained for subsequent analyses.

The statistical significance of PLS1 was assessed using 5,000 permutations. Gene contributions to PLS1 were evaluated using 1,000 bootstrap resamples. To account for sign indeterminacy in PLS solutions, bootstrap-derived gene weights were aligned to the original PLS1 weight vector before estimating standard errors. A Z-score was calculated for each gene by dividing its PLS1 weight by the corresponding bootstrap standard error. Genes were ranked according to their bootstrap ratios, and genes with | Z| > 5 were retained as PLS1-associated genes for downstream analyses.

We next examined transcriptomic similarity across all architecture-state combinations using two complementary analyses. First, pairwise Pearson correlations were calculated between PLS1 scores across the ten architecture-state models (*Supplementary Material S3*). Second, pairwise Pearson correlations were calculated between whole-brain PLS1 gene-loading vectors across the same ten models to evaluate the similarity of transcriptomic signatures. Absolute Pearson correlation coefficients were used in all similarity analyses to emphasize the strength of shared transcriptomic organization irrespective of directionality.

To characterize the biological functions associated with the identified transcriptomic families (*Supplementary Material S3*), family-specific gene lists were constructed. For each transcriptomic family, the final family-specific gene list was defined as the intersection of PLS1-associated genes across all architecture-state members belonging to that family. The resulting family-specific gene sets were subsequently subjected to Metascape analysis [66] to identify enriched Gene Ontology (GO) biological processes and Kyoto Encyclopedia of Genes and Genomes (KEGG) pathways. Only pathways surviving false discovery rate correction (*P*_*FDR*_ < 0.01) were retained for further interpretation and visualization.

### 5.8 External validation of neurodynamic architectures

An independent validation cohort of 65 participants with MDD was analyzed to evaluate the reproducibility of the neurodynamic architectures identified in the discovery cohort. Resting-state fMRI data were preprocessed using procedures consistent with those applied to the discovery dataset. Given the relatively limited sample size of the validation cohort and to preserve cross-cohort comparability, the discovery-derived HMM was applied directly to the validation data without retraining or parameter optimization. The number of states and all state-specific model parameters were fixed, and each participant’s resting-state time series was decoded within the discovery-derived state space. Posterior state probabilities and discrete state sequences were subsequently obtained. Subject-level FO, SR, and LT were calculated for each of the five states using the same procedures as in the discovery cohort, yielding 15 homologous dynamic features across datasets.

The primary validation analysis examined whether a similar neurodynamic organization could be independently recovered without imposing discovery-derived architecture assignments. The 15 dynamic features were standardized within the validation cohort, and hierarchical clustering was performed using Euclidean distance, Ward’s linkage, and a prespecified two-cluster solution consistent with the discovery analysis, resulting in two independently derived validation subgroups.

As a supplementary transfer-based analysis, the discovery-derived architecture definition was applied directly to the validation participants. Validation features were standardized using the means and standard deviations estimated exclusively from the discovery cohort. Centroids for Architecture 1 and Architecture 2 were calculated in the standardized discovery feature space, and each participant was assigned to the architecture with the smaller Euclidean distance. No standardization parameters or centroids were re-estimated in the validation cohort. After label alignment, centroid-based assignments were compared with those obtained from the primary independent re-clustering analysis.

## Acknowledgments

This work was supported by the Hong Kong RGC Strategic Target Grant (RGC grant STG1/M-501/23-N), the Hong Kong Health and Medical Research Fund (HMRF grant 10211696), the Hong Kong Global STEM Scholar Scheme, and the Hong Kong Jockey Club Charities Trust.

## Author contributions

Conceptualization: W.Z., S.Q and F.W. Methodology: C.P., and Y.T. Software: C.P. Validation: C.P. Formal Analysis: C.P. Investigation: C.P., and Y.T. Resources: W.Z., and S.Q. Data Collection & Curation: Y.T., K.L., and R. C. Writing – Original Draft: C.P. Writing – Review & Editing: All authors. Visualization: C.P. Supervision: W.Z. and S.Q. Project Administration: W.Z. and S.Q. Funding Acquisition: W.Z.

## Reference

[1] W. Marx et al., “Major depressive disorder,” Nature Reviews Disease Primers, vol. 9, no. 1, p. 44, 2023.

[2] C. J. Murray, “The global burden of disease study at 30 years,” Nature medicine, vol. 28, no. 10, pp. 2019–2026, 2022.

[3] C. J. Murray, “Findings from the global burden of disease study 2021,” The Lancet, vol. 403, no. 10440, pp. 2259–2262, 2024.

[4] L. Tozzi et al., “Personalized brain circuit scores identify clinically distinct biotypes in depression and anxiety,” Nature medicine, vol. 30, no. 7, pp. 2076–2087, 2024.

[5] Q. Liu et al., “Multilevel brain functional connectivity and task-based representations explaining heterogeneity in major depressive disorder,” Translational Psychiatry, vol. 15, no. 1, p. 199, 2025.

[6] S. Marek et al., “Reproducible brain-wide association studies require thousands of individuals,” Nature, vol. 603, no. 7902, pp. 654–660, 2022.

[7] M. A. Di Biase et al., “Biochemical and brain heterogeneity characterizes psychiatric and non-psychiatric illness,” Nature Communications, 2026.

[8] P. C. Mulders, P. F. van Eijndhoven, A. H. Schene, C. F. Beckmann, and I. Tendolkar, “Resting-state functional connectivity in major depressive disorder: a review,” Neuroscience & Biobehavioral Reviews, vol. 56, pp. 330–344, 2015.

[9] B. B. Biswal and L. Q. Uddin, “The history and future of resting-state functional magnetic resonance imaging,” Nature, vol. 641, no. 8065, pp. 1121–1131, 2025.

[10] J. Li et al., “Major depressive disorder on a neuromorphic continuum,” Nature communications, vol. 16, no. 1, p. 2405, 2025.

[11] L. M. Williams, “Precision psychiatry: a neural circuit taxonomy for depression and anxiety,” The Lancet Psychiatry, vol. 3, no. 5, pp. 472–480, 2016.

[12] Z. Luo et al., “Identifying neurophenotypes of major depressive disorder through normative model of regional homogeneity,” Translational Psychiatry, 2026.

[13] C. Chen et al., “Stable depression subtypes identified using functional connectome normative deviation models and their response to rTMS,” Molecular Psychiatry, pp. 1–10, 2026.

[14] X. Wang et al., “Unveiling diverse clinical symptom patterns and neural activity profiles in major depressive disorder subtypes,” EBioMedicine, vol. 116, 2025.

[15] A. T. Drysdale et al., “Resting-state connectivity biomarkers define neurophysiological subtypes of depression,” Nature medicine, vol. 23, no. 1, pp. 28–38, 2017.

[16] K. Fang et al., “Individualized resting-state functional connectivity abnormalities unveil two major depressive disorder subtypes with contrasting abnormal patterns of abnormality,” Translational Psychiatry, vol. 15, no. 1, p. 45, 2025.

[17] Q. Li et al., “Heterogeneity of brain functional connectivity, transcriptome, and neurotransmitter profiles in major depressive disorder,” Psychological Medicine, vol. 55, p. e341, 2025.

[18] B. Ü. Kilic, J. Jubeir, P. Balchandani, J. W. Murrough, L. S. Morris, and Y. Jacob, “Spatiotemporal asymmetries on brain energy landscape uncover system entrapment related to depression severity,” Nature Communications, 2026.

[19] A. Cai et al., “Unraveling spatiotemporal dynamics in transdiagnosis subtypes of major depressive disorder and bipolar disorder: insights from co-activation patterns and treatment response,” Molecular Psychiatry, vol. 31, no. 3, pp. 1810–1822, 2026.

[20] B. Klugah-Brown et al., “Dynamic brain connectivity patterns induced by oxytocin: An fMRI Co-Activation pattern analysis study,” Molecular Psychiatry, pp. 1–13, 2026.

[21] X. Liu et al., “Subtypes of adolescent major depressive disorder characterized by divergent information dynamics in sensory-association cortices,” Nature Communications, 2026.

[22] K. Finc, I. Adamska-Stolarczyk, and D. S. Bassett, “Flexible modularity in the human brain: How network architecture reconfigures over time,” Neuroscience & Biobehavioral Reviews, vol. 188, p. 106784, 2026.

[23] X. Chang et al., “Neurocognitive characterization of behaviour and mental illness through time-varying brain network analysis,” Nature Communications, 2026.

[24] C. V. Cocuzza et al., “Brain network dynamics reflect psychiatric illness status and transdiagnostic symptom profiles across health and disease,” Nature Communications, vol. 17, no. 1, p. 6678, 2026/07/21 2026, doi: 10.1038/s41467-026-75585-6.

[25] M. W. van Es, C. Higgins, C. Gohil, A. J. Quinn, D. Vidaurre, and M. W. Woolrich, “Large-scale cortical functional networks are organized in structured cycles,” Nature Neuroscience, vol. 28, no. 10, pp. 2118–2128, 2025.

[26] J. Li et al., “Hierarchical neural variability reveals adaptive and maladaptive mechanisms of non-suicidal self-injury,” Molecular Psychiatry, 2026/06/05 2026, doi: 10.1038/s41380-026-03671-8.

[27] T. H. Alderson et al., “Shared spatial and temporal principles govern connectome dynamics across timescales,” Proceedings of the National Academy of Sciences, vol. 123, no. 25, p. e2535464123, 2026.

[28] C. Xue and G. Okazawa, “The ‘neat’and ‘messy’in task-dependent neural geometry and computation,” Trends in Neurosciences, 2026.

[29] A. S. Greene, C. Horien, D. Barson, D. Scheinost, and R. T. Constable, “Why is everyone talking about brain state?,” Trends in neurosciences, vol. 46, no. 7, pp. 508–524, 2023.

[30] Q. Liu et al., “Energy inefficiency underpinning brain state dysregulation in individuals with major depressive disorder,” Nature Mental Health, vol. 4, no. 3, pp. 400–415, 2026.

[31] T. Liu et al., “Heterogeneous functional state dynamics and its structural substrates in male individuals with autism spectrum disorder,” Molecular Psychiatry, pp. 1–14, 2026.

[32] L. Geng et al., “Depression links to unstable resting-state brain dynamics: insights from hidden markov models and functional network variability,” Psychological Medicine, vol. 55, p. e200, 2025.

[33] I. Rentzeperis, J. Li, R. Bauer, and C. van Leeuwen, “Opening the black box of neural variability: from noise to mechanisms,” Neuroscience & Biobehavioral Reviews, p. 106661, 2026.

[34] F. Hancock et al., “Metastability demystified—the foundational past, the pragmatic present and the promising future,” Nature Reviews Neuroscience, vol. 26, no. 2, pp. 82–100, 2025.

[35] A. Wolff and G. Dumas, “Coupled neural timescales in social interaction,” Trends in Cognitive Sciences, 2026.

[36] Y. Shi et al., “Dynamic functional connectivity and transcriptomic signatures reveal stage-dependent brain network dysfunction in Alzheimer’s disease spectrum,” Alzheimer’s Research & Therapy, vol. 17, no. 1, p. 247, 2025.

[37] J. Shao et al., “Neuroimaging of Heterogeneity in Neuropsychiatric Disorders: Toward Disease Progression Modeling,” Biological Psychiatry, 2026.

[38] J. Li et al., “Cortical structural differences in major depressive disorder correlate with cell type-specific transcriptional signatures,” Nature communications, vol. 12, no. 1, p. 1647, 2021.

[39] C. Pan, Y. Ma, L. Wang, Y. Zhang, F. Wang, and X. Zhang, “From connectivity to controllability: Unraveling the brain biomarkers of major depressive disorder,” Brain Sciences, vol. 14, no. 5, p. 509, 2024.

[40] Y. Chen et al., “State-and trait-related dysfunctions in bipolar disorder across different mood states: a graph theory study,” Journal of Psychiatry and Neuroscience, vol. 49, no. 1, pp. E11-E22, 2024.

[41] L. Tang et al., “Epigenetic molecular underpinnings of brain structural-functional connectivity decoupling in patients with major depressive disorder,” Journal of Affective Disorders, vol. 363, pp. 249–257, 2024.

[42] X. Zhang et al., “Cancer-keeper genes as therapeutic targets,” Iscience, vol. 26, no. 8, 2023.

[43] X. Wei, C. Pan, X. Zhang, and W. Zhang, “Total network controllability analysis discovers explainable drugs for Covid-19 treatment,” Biology direct, vol. 18, no. 1, p. 55, 2023.

[44] X. Zhang, C. Pan, and W. Zhang, “Control hubs of complex networks and a polynomial-time identification algorithm,” arXiv preprint arXiv:2206.01188, 2022.

[45] C. Pan et al., “Adaptive control of dynamic networks,” IEEE Transactions on Network Science and Engineering, 2025.

[46] C.-G. Yan et al., “Reduced default mode network functional connectivity in patients with recurrent major depressive disorder,” Proceedings of the National Academy of Sciences, vol. 116, no. 18, pp. 9078–9083, 2019.

[47] X. Chen et al., “The DIRECT consortium and the REST-meta-MDD project: towards neuroimaging biomarkers of major depressive disorder,” Psychoradiology, vol. 2, no. 1, pp. 32–42, 2022.

[48] J. B. Williams, “A structured interview guide for the Hamilton Depression Rating Scale,” Archives of general psychiatry, vol. 45, no. 8, pp. 742–747, 1988.

[49] J. D. Power et al., “Functional network organization of the human brain,” Neuron, vol. 72, no. 4, pp. 665–678, 2011.

[50] D. Vidaurre et al., “The Gaussian-linear hidden Markov model: A Python package,” Imaging Neuroscience, vol. 3, p. pimag_a_00460, 2025.

[51] X. Zhang et al., “Reconfiguration of brain network dynamics in bipolar disorder: a hidden Markov model approach,” Translational Psychiatry, vol. 14, no. 1, p. 507, 2024.

[52] N. Javaheripour et al., “Altered brain dynamic in major depressive disorder: state and trait features,” Translational Psychiatry, vol. 13, no. 1, p. 261, 2023.

[53] D. Vidaurre, S. M. Smith, and M. W. Woolrich, “Brain network dynamics are hierarchically organized in time,” Proceedings of the National Academy of Sciences, vol. 114, no. 48, pp. 12827–12832, 2017.

[54] W. S. Noble, “What is a support vector machine?,” Nature biotechnology, vol. 24, no. 12, pp. 1565–1567, 2006.

[55] J. Friedman, T. Hastie, and R. Tibshirani, “Sparse inverse covariance estimation with the graphical lasso,” Biostatistics, vol. 9, no. 3, pp. 432–441, 2008.

[56] D. Borsboom et al., “Network analysis of multivariate data in psychological science,” Nature reviews methods primers, vol. 1, no. 1, p. 58, 2021.

[57] C. Pan et al., “Module control of network analysis in psychopathology,” Iscience, vol. 27, no. 7, 2024.

[58] F. Pedregosa et al., “Scikit-learn: Machine learning in Python,” the Journal of machine Learning research, vol. 12, pp. 2825–2830, 2011.

[59] L. C. Freeman, “Centrality in social networks conceptual clarification,” Social networks, vol. 1, no. 3, pp. 215–239, 1978.

[60] Z. Fei et al., “Reproducible psychosocial–emotional profiles reveal population-level heterogeneity in young adults,” PREPRINT (Version 1) available at Research Square, 2026, doi: 10.21203/rs.3.rs-10317962/v1.

[61] M. R. Chernick, Bootstrap methods: A guide for practitioners and researchers. John Wiley & Sons, 2011.

[62] S. Epskamp, D. Borsboom, and E. I. Fried, “Estimating psychological networks and their accuracy: A tutorial paper,” Behavior Research Methods, vol. 50, no. 1, p. 195, 2017.

[63] M. J. Hawrylycz et al., “An anatomically comprehensive atlas of the adult human brain transcriptome,” Nature, vol. 489, no. 7416, pp. 391–399, 2012.

[64] R. Markello, G. Shafiei, Y.-Q. Zheng, and B. Mišić, “abagen: A toolbox for the Allen Brain Atlas genetics data,” Zenodo, 2020.

[65] H. Abdi, “Partial least squares regression and projection on latent structure regression (PLS Regression),” Wiley interdisciplinary reviews: computational statistics, vol. 2, no. 1, pp. 97–106, 2010.

[66] Y. Zhou et al., “Metascape provides a biologist-oriented resource for the analysis of systems-level datasets,” Nature communications, vol. 10, no. 1, p. 1523, 2019.

