## Supplementary Material for "Temporal organization of shared brain states characterizes heterogeneity in major depressive disorder"

**– Supplementary information**

**SUPPLEMENTARY MATERIAL**

- ***Supplementary Material S1.*** *Selection of the final number of hidden states*
- ***Supplementary Material S2.*** *Identification and stability assessment of neurodynamic architectures*
- ***Supplementary Material S3.*** *Identification of transcriptomic similarity structures and state families*

**SUPPLEMENTARY FIGURES**

- ***Supplementary Figure S1.*** *Evaluation of model fit across different numbers of hidden states*
- ***Supplementary Figure S2.*** *Across-run stability of HMM state trajectories*
- ***Supplementary Figure S3.*** *Spatial configurations of the five inferred brain states*
- ***Supplementary Figure S4.*** *sPCA identifies dominant dimensions of brain-state heterogeneity*
- ***Supplementary Figure S5.*** *Hierarchical clustering and consensus structure of neurodynamic architectures*
- ***Supplementary Figure S6.*** *Stability distribution of architecture assignments across repeated subsampling*
- ***Supplementary Figure S7.*** *Cross-validated classification of MDD architectures versus controls*
- ***Supplementary Figure S8.*** *State-specific functional configurations of neurodynamic architectures*
- ***Supplementary Figure S9****. Bootstrap stability of symptom-network across neurodynamic architectures*
- ***Supplementary Figure S10****. PLS1 associations with neuroimaging phenotypes*
- ***Supplementary Figure S11.*** *Similarity of PLS1 spatial score patterns across architecture–state combinations*
- ***Supplementary Figure S12****. Brain-state configurations in the validation cohort*
- ***Supplementary Figure S13****. FO profiles of neurodynamic architectures in the independent validation cohort*
- ***Supplementary Figure S14.*** *Temporal dynamics of validation neurodynamic architectures*

**SUPPLEMENTARY TABLES**

- ***Supplementary Table S1.*** *Pairwise comparisons of all samples among the five recurrent brain states*
- ***Supplementary Table S2.*** *Transition probability matrix of the five recurrent brain states under all samples*
- ***Supplementary Table S3.*** *HC–MDD group comparison of brain-state features*
- ***Supplementary Table S4.*** *HAMD associations with brain-state features*
- ***Supplementary Table S5.*** *Statistical test of brain-state dynamic features between neurodynamic architectures*
- ***Supplementary Table S6.*** *Architecture differences in brain-state transition probabilities*
- ***Supplementary Table S7.*** *neurodynamic architecture features associations with HAMD scores*
- ***Supplementary Table S8.*** *Symptom network centrality profiles of neurodynamic architectures*
- ***Supplementary Table S9.*** *PLS component variance explained in neurodynamic architecture–states*
- ***Supplementary Table S10.*** *Spatial correlations between ROI activity patterns and PLS1 scores*
- ***Supplementary Table S11.*** *Enrichment analysis results of transcriptomic Family 3*
- ***Supplementary Table S12.*** *Jaccard similarity of PLS1-associated gene sets across architecture–state combinations*
- ***Supplementary Table S13.*** *Overlap coefficient of PLS1-associated genes across architecture–state combinations*
- ***Supplementary Table S14.*** *Sparse principal component loadings of brain-state features*
- ***Supplementary Table S15.*** *Silhouette coefficients across candidate cluster numbers*
- ***Supplementary Table S16.*** *Distribution of neurodynamic architecture composition across sites*
- ***Supplementary Table S17.*** *Comparison of two clustering strategies in the validation cohort*

**SUPPLEMENTARY MATERIAL**

***Supplementary Material* *S1. Selection of the final number of hidden states***

To determine the optimal number of hidden states for the Gaussian hidden Markov model (HMM), models with different numbers of states (K = 2–15) were systematically evaluated. For each candidate K, the HMM was independently fitted across 10 runs using different random seeds. Model fitting was performed using the GLHMM toolbox with state-specific mean parameters and covariance structures.

Model fit and complexity were evaluated using variational free energy, Akaike information criterion (AIC), and Bayesian information criterion (BIC). Although all three criteria were calculated during model evaluation, AIC and BIC showed highly consistent patterns across candidate state numbers. Therefore, only BIC and variational free energy were visualized for presentation (*Supplementary Figure S1*). To identify candidate solutions with an optimal balance between model fit and model complexity, the second-order differences of BIC and variational free energy were examined. Both criteria showed the smallest changes around K = 5 and K = 6, indicating these solutions as candidate state numbers.

Because model fit alone may favor additional states that do not represent reproducible neural configurations, the stability of inferred brain states was further assessed across repeated model estimations (Supplementary Figure S2). For each candidate K, posterior state probability trajectories obtained from different runs were compared using Pearson correlation after state matching with the Hungarian algorithm. The average matched correlation across all pairwise comparisons was used as an index of state reproducibility. The K = 5 solution demonstrated consistently high across-run similarity, whereas the K = 6 solution showed substantially reduced similarity, indicating less stable state decomposition. Therefore, integrating model fit, complexity, and reproducibility, the five-state solution was selected as the optimal model order and used for all subsequent analyses. The spatial configurations of the five inferred brain states across the 264 Power atlas regions are illustrated in Supplementary Figure S3.

***Supplementary Material* *S2. Identification and stability assessment of neurodynamic architectures***

To determine whether individual variability in brain-state organization reflected reproducible neurodynamic patterns rather than arbitrary partitioning, we systematically evaluated feature representation, clustering solutions, and architecture stability within the MDD cohort.

All candidate features were standardized across participants using z-score normalization before sparse principal component analysis (sPCA). sPCA was applied to characterize the dominant dimensions underlying inter-individual variability in brain-state organization rather than to directly select features for clustering. To determine an appropriate sparse representation, the number of sparse components and sparsity penalty were systematically evaluated. Candidate models were tested with the number of components ranging from 5 to 15 and sparsity parameter α ranging from 0.1 to 5.0. Models were evaluated according to clustering separation in the resulting latent space, quantified by the silhouette coefficient, together with component sparsity and reconstruction performance. The optimal sPCA configuration consisted of five sparse components with α = 5.0. The loading patterns of the five sparse components were subsequently examined. The first sparse component showed predominant contributions from dynamic features, indicating that temporal organization of recurrent brain states represented a major source of heterogeneity within MDD. Based on this observation, the complete set of dynamic features, including FO, SR, and LT across all five states, was retained for subsequent architecture identification. Detailed sPCA optimization results and loading patterns are shown in *Supplementary Figure S4* and *Supplementary Table S14*.

Hierarchical clustering was subsequently performed using the retained dynamic feature space. Before clustering, dynamic features were again standardized using z-score normalization. Pairwise Euclidean distances were calculated, and agglomerative hierarchical clustering was performed using Ward’s linkage criterion. Candidate solutions ranging from K = 2 to K = 9 were generated by cutting the hierarchical tree at different cluster numbers. The optimal number of clusters was determined based on the average silhouette coefficient (*Supplementary Table S15*), and the resulting cluster assignments were used as the final architecture labels. The hierarchical clustering structure and consensus matrix are presented in *Supplementary Figure S5*.

The reproducibility of the identified architecture solution was evaluated using repeated subsampling analyses. In each of 1,000 iterations, 80% of MDD participants were randomly sampled without replacement, and hierarchical clustering was repeated using the same dynamic feature space and clustering procedure. The global stability of the partition structure was assessed by comparing each subsampled solution with the full-sample solution using adjusted Rand index (ARI, 0.42 ± 0.18), normalized mutual information (NMI, 0.39 ± 0.10), and assignment matching rate after cluster label alignment. In addition, subject-level assignment stability was calculated as the proportion of iterations in which each participant retained the same architecture assignment among all iterations in which that participant was included. The mean and median subject-level stability were 0.81 and 0.90, respectively, with 800 and 592 participants showing stability values ≥0.80 and ≥0.90, respectively. Pairwise clustering consistency was further evaluated using a consensus matrix, where each element represented the proportion of iterations in which two participants were assigned to the same architecture. The distribution of assignment stability across repeated subsampling is shown in *Supplementary Figure S6*.

After architectureidentification, state-specific functional configurations were further examined to determine whether architecture differences reflected distinct brain-state repertoires or differential organization of shared states. For each architecture, ROI-level activity patterns across the five recurrent brain states were averaged within the 12 Power atlas functional communities and visualized as radar plots (*Supplementary Figure S8*). The spatial configurations of the five brain states were highly similar between the two neurodynamic architectures and the overall MDD cohort, indicating that architecture differentiation was not driven by the emergence of distinct functional states. Instead, the identified architectures primarily reflected differences in the temporal organization of a shared repertoire of recurrent brain states.

***Supplementary Material* *S3. Identification of transcriptomic similarity structures and state families***

To determine whether the transcriptomic state families identified from PLS1 gene-loading similarity represented robust molecular organizations, we performed complementary similarity analyses across all architecture–state-specific PLS models. Each architecture–state combination generated a PLS1 component that characterized the spatial relationship between regional imaging patterns and whole-brain gene-expression profiles. Similarity among these PLS1-derived transcriptomic patterns was evaluated from three complementary perspectives: spatial organization of transcriptomic gradients, whole-transcriptome gene-loading patterns, and overlap of PLS1-associated core genes.

For each architecture–state-specific PLS model, two continuous representations were obtained from the PLS1 component. First, PLS1 spatial scores were extracted across the 125 cortical regions included in the analysis. These spatial scores characterize the anatomical distribution of the transcriptomic gradient associated with each brain-state pattern. Pairwise Pearson correlations were calculated between PLS1 spatial score vectors across the ten architecture–state combinations to evaluate the similarity of spatial organization of transcriptomic gradients (Supplementary Figure S11). These analyses assessed whether different brain-state configurations shared similar regional distributions of transcriptomic patterns.

Second, PLS1 gene-loading vectors were extracted across all analyzed genes. Each vector contained the loading scores of 15,632 genes, representing the contribution of individual genes to the corresponding transcriptomic component. Pairwise Pearson correlations between whole-transcriptome gene-loading vectors were calculated to quantify molecular similarity among architecture–state combinations. The resulting similarity matrix was used to characterize the transcriptomic organization shown in Fig. 6E and identify candidate transcriptomic state families.

To further evaluate whether the identified similarity structure was driven by shared high-contribution genes rather than global correlation patterns across the entire transcriptome, we performed a complementary core gene overlap analysis. For each architecture–state-specific PLS model, genes with strong contributions to PLS1 were identified based on bootstrap ratios (|Z| > 5), consistent with the gene-selection procedure described in Methods. The resulting PLS1-associated gene sets were converted into binary gene membership profiles, where genes retained in each model were coded as 1 and other genes were coded as 0. Pairwise similarity between gene sets was then quantified using two complementary overlap measures: the Jaccard coefficient, which reflects the proportion of shared genes relative to the union of two gene sets (*Supplementary Table S12*), and the overlap coefficient, which evaluates the proportion of shared genes relative to the smaller gene set (*Supplementary Table S13*). These analyses were used to determine whether architecture–state combinations assigned to the same transcriptomic family shared convergent molecular components.

Across the three complementary similarity analyses, architecture–state combinations demonstrated a convergent transcriptomic organization. Brain states preferentially engaged by architecture1 (ACS and VSS) showed highly similar transcriptomic patterns across spatial gradients, whole-transcriptome loading profiles, and core gene composition, forming a shared transcriptomic family. Similarly, MAS and DMS across both architectures exhibited consistent transcriptomic similarity and formed a second transcriptomic family. THS displayed a distinct transcriptomic organization compared with the other states and formed a separate family. Together, these convergent analyses support that the identified transcriptomic state families reflect stable molecular pattern rather than similarity patterns arising from a single analytical representation.

**SUPPLEMENTARY FIGURES**

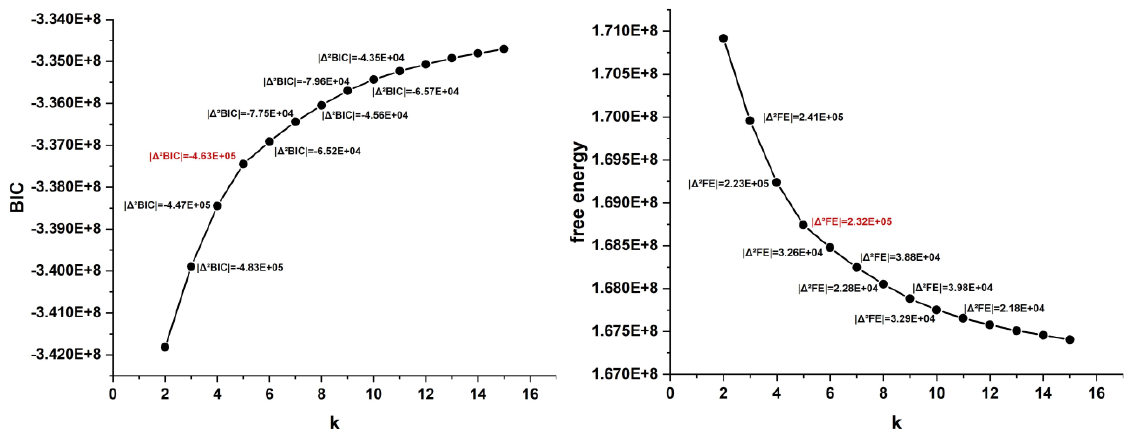

**Supplementary Figure S1. Evaluation of model fit across different numbers of hidden states.** Bayesian information criterion (BIC; left panel) and variational free energy (right panel) were calculated for Gaussian HMMs with different numbers of hidden states (K = 2–15). Each model was fitted across 10 independent runs, and the mean values across runs were used for visualization. The second-order differences of BIC and free energy were calculated to identify points where increasing model complexity yielded diminishing improvements. Both criteria indicated that K = 5 and K = 6 represented candidate solutions with optimal model fit and complexity balance.

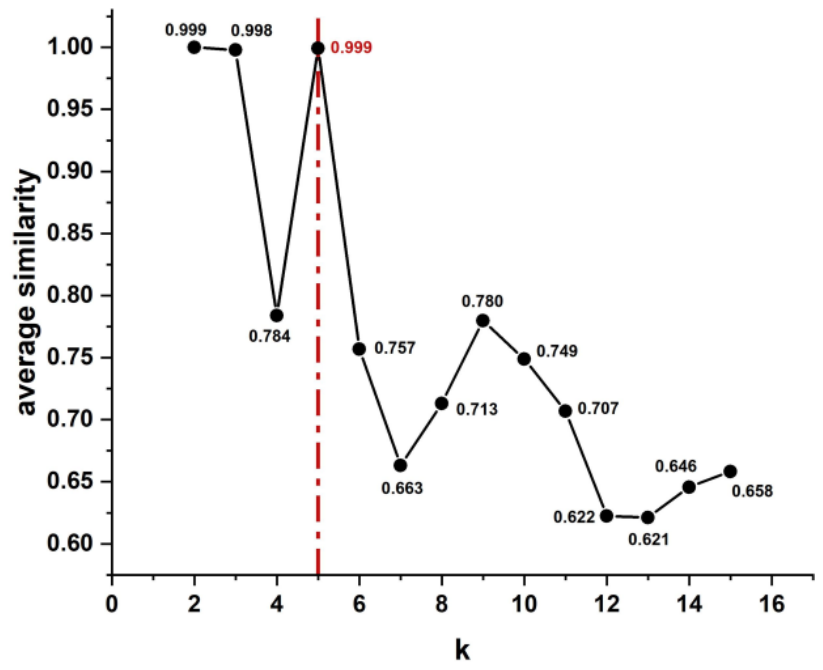

**Supplementary Figure S2. Across-run stability of HMM state trajectories.** Model stability was evaluated by comparing posterior state probability trajectories (Gamma matrices) across repeated HMM estimations. For each candidate number of states (K = 2–15), pairwise correlations between state-specific Gamma time courses were calculated after state matching using the Hungarian algorithm. The average matched correlation across all run pairs was used as the stability metric. The five-state solution showed high reproducibility across independent runs, whereas the six-state solution exhibited reduced stability, supporting the selection of K = 5 as the optimal state number.

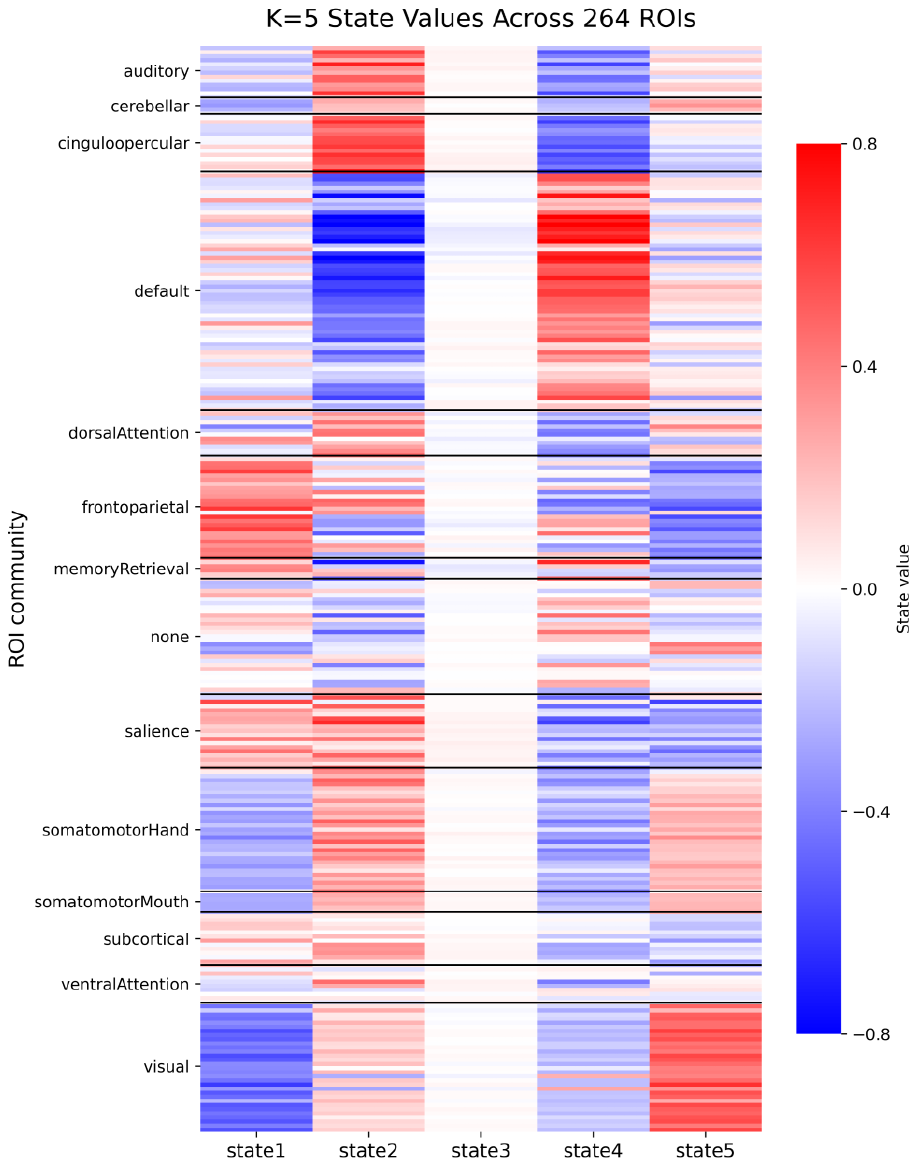

**Supplementary Figure S3. Spatial configurations of the five inferred brain states.** The heatmap illustrates the state-specific mean activity patterns estimated by the HMM with five hidden states (K = 5). Each column represents one inferred brain state, and each row represents one region of interest (ROI) from the Power264 functional atlas. Color intensity indicates the normalized state-specific activity value of each ROI, reflecting the spatial configuration of functional engagement associated with each brain state. Horizontal lines indicate the boundaries between functional communities defined by the Power atlas. state1: ACS; state2: MAS; state3: THS; state4: DMS; state5: VSS;

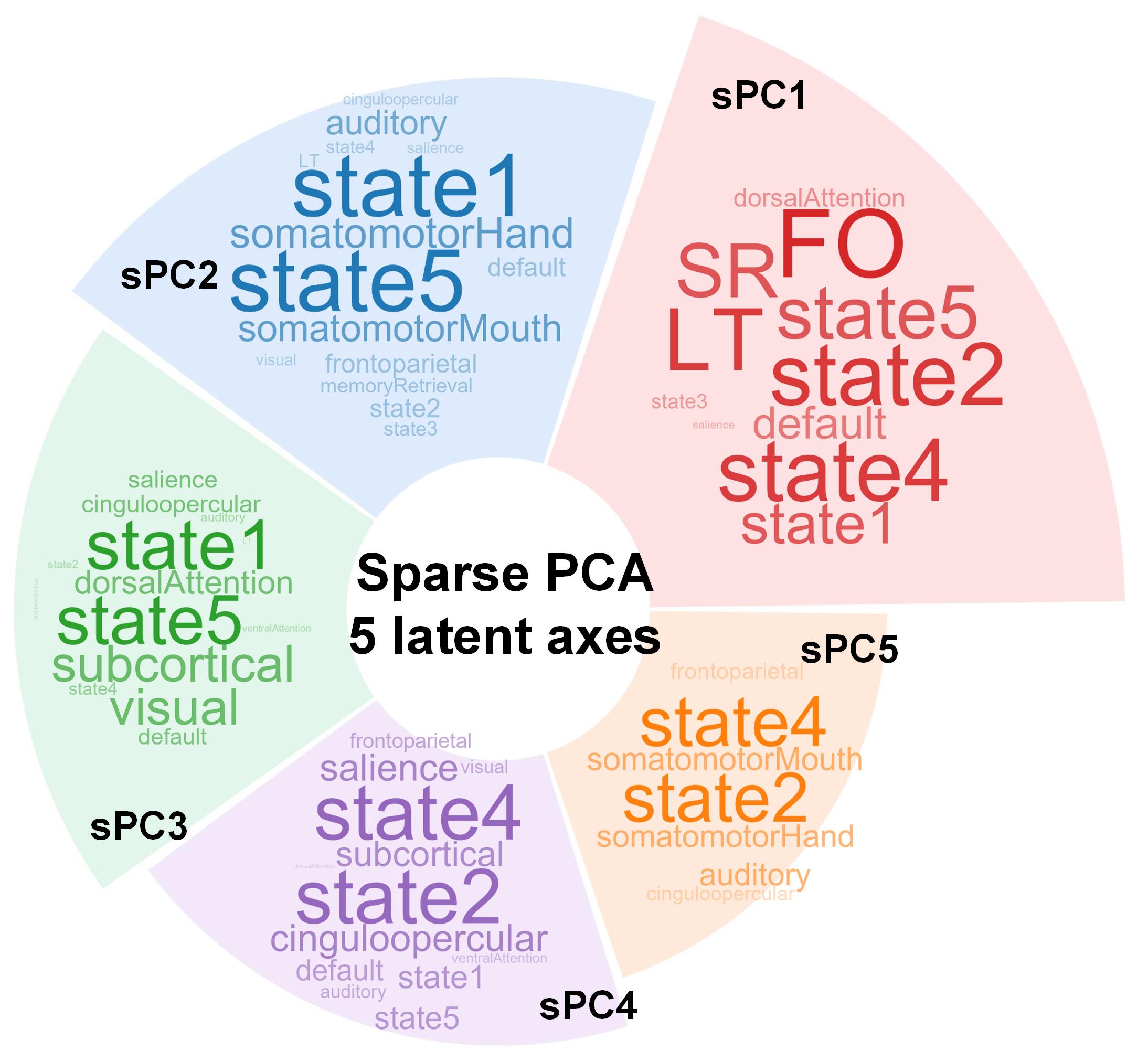

**Supplementary Figure S4. sPCA identifies dominant dimensions of brain-state heterogeneity.** Sparse principal component analysis (sPCA) was applied to the 75-dimensional brain-state feature space consisting of state-specific functional and dynamic features. The five sparse components represent major latent axes of inter-individual variation within the MDD cohort. Feature contributions and loading patterns indicate that the first sparse component was predominantly characterized by dynamic features, supporting the use of temporal brain-state features for subsequent architecture identification. state1: ACS; state2: MAS; state3: THS; state4: DMS; state5: VSS;

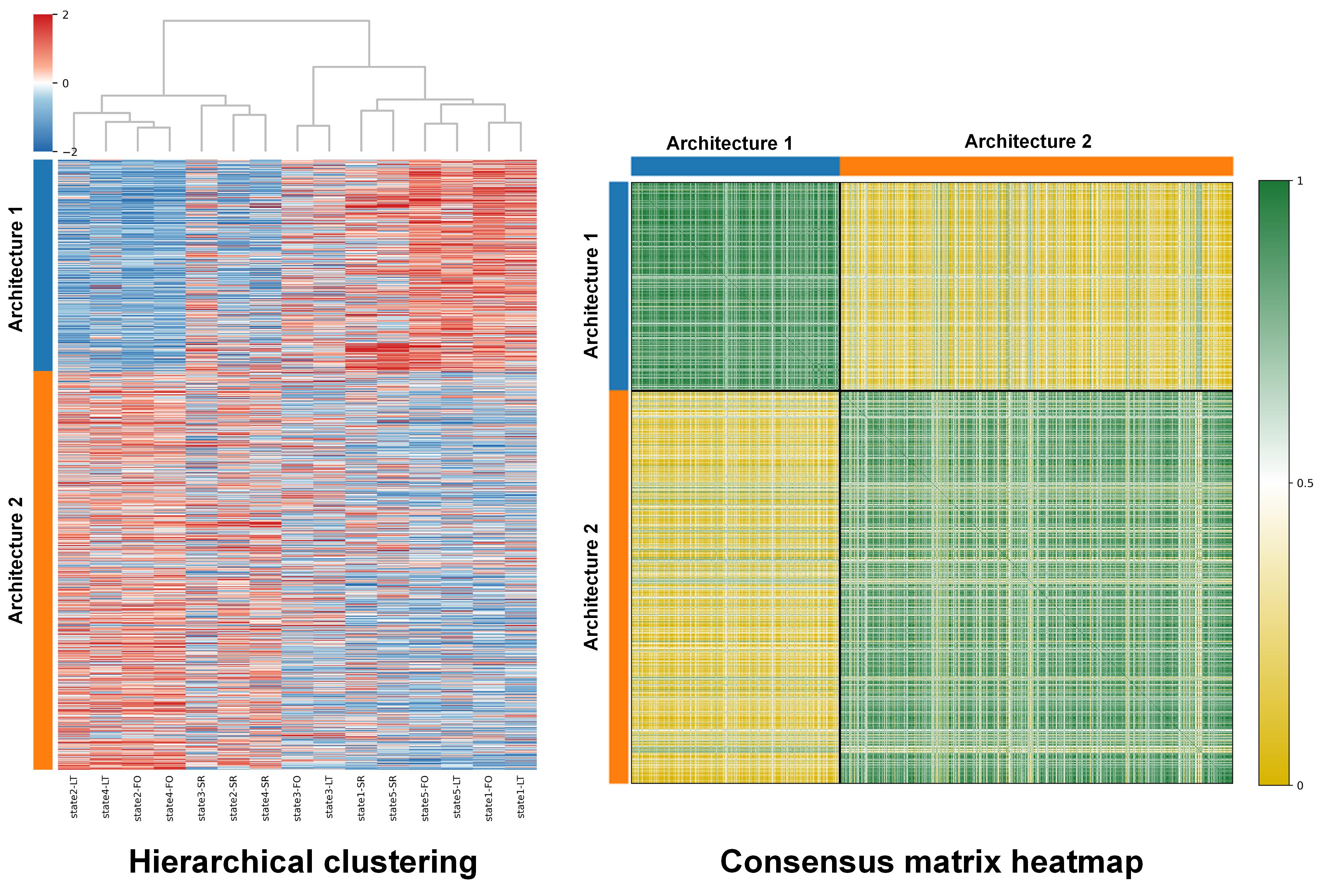

**Supplementary Figure S5. Hierarchical clustering and consensus structure of neurodynamic architectures.** Hierarchical clustering was performed using the dynamic feature space consisting of fractional occupancy, switching rate, and lifetime across the five brain states. The left panel shows the clustering structure derived from Ward linkage after z-score normalization of dynamic features. The right panel presents the consensus matrix obtained from repeated clustering, where each element represents the proportion of iterations in which a participant pair was assigned to the same architecture. The distinct block structure supports the separation of two neurodynamic architectures. state1: ACS; state2: MAS; state3: THS; state4: DMS; state5: VSS;

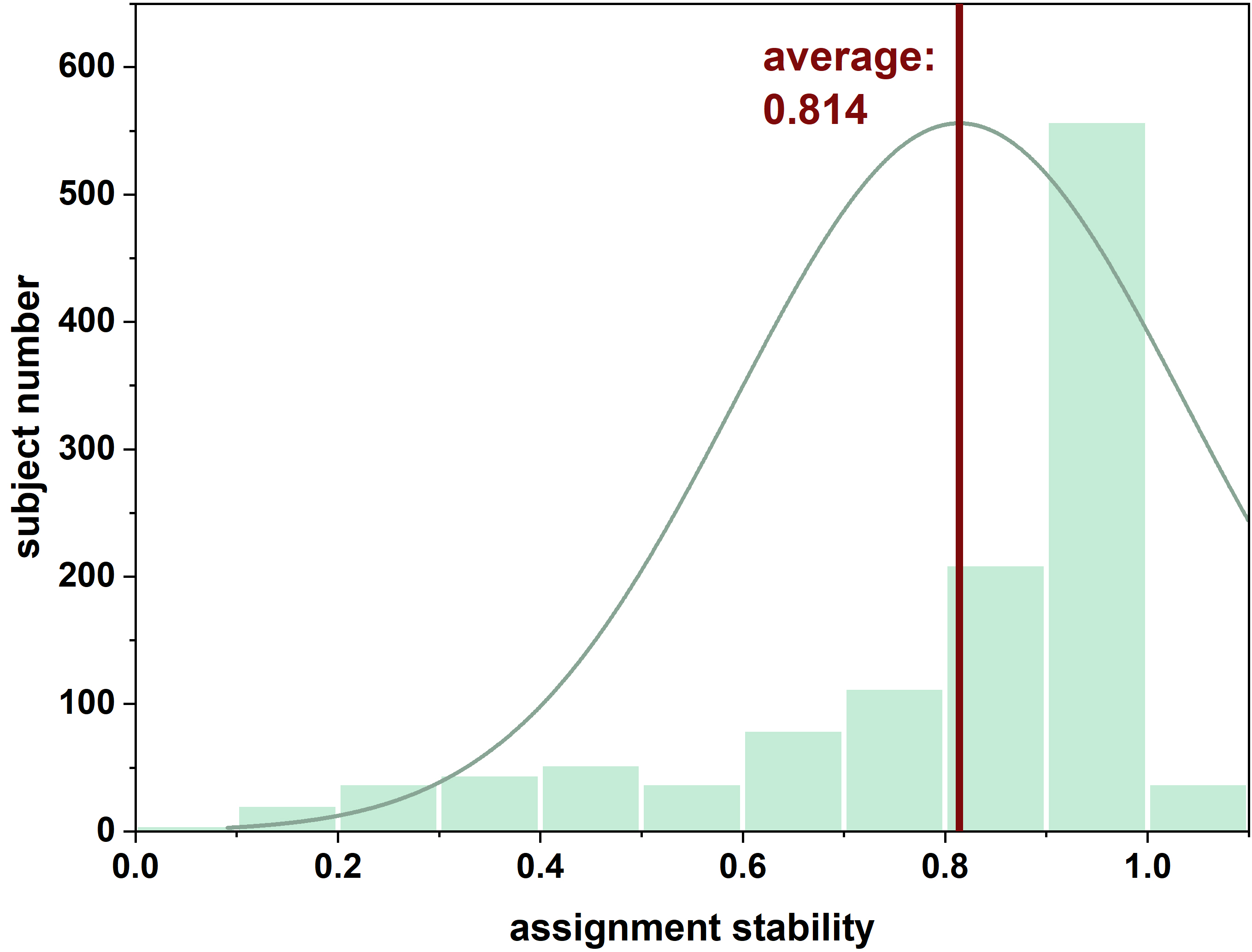

**Supplementary Figure S6. Stability distribution of architectures assignments across repeated subsampling.** Architecture stability was assessed through 1,000 repeated subsampling analyses, in which 80% of MDD participants were randomly selected in each iteration and re-clustered using the same dynamic feature space and clustering procedure. The histogram shows the distribution of subject-level assignment stability, defined as the proportion of included iterations in which each participant retained the same architecture assignment as in the full-sample solution. The high average stability indicates robust reproducibility of the identified two- architecture solution.

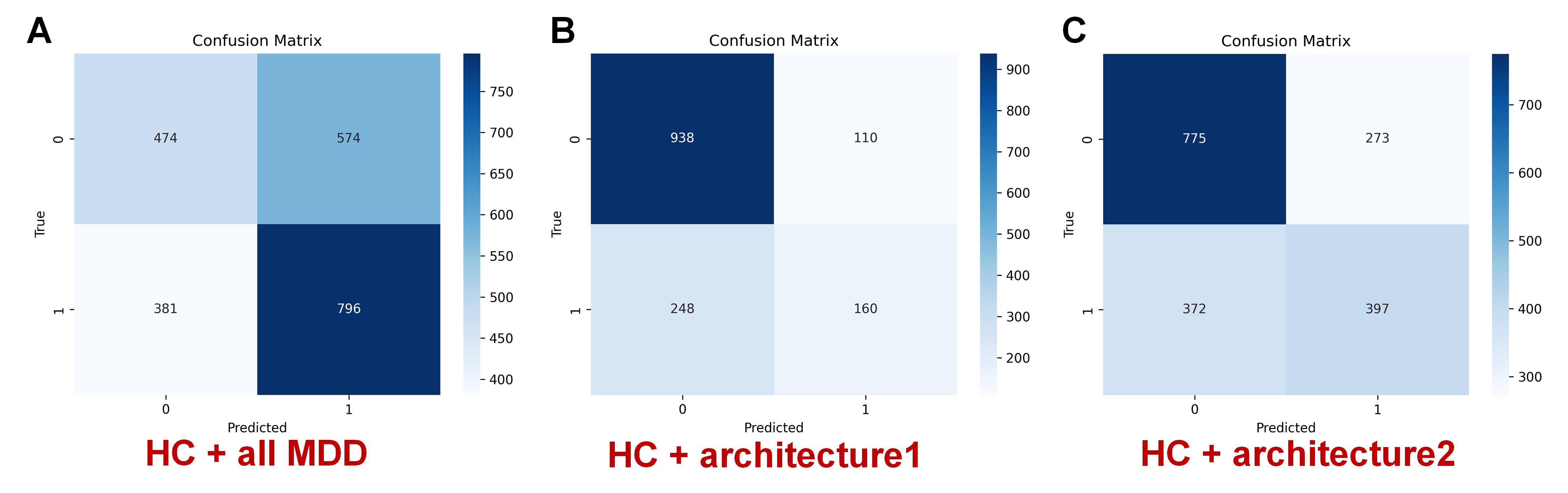

**Supplementary Figure S7. Cross-validated classification of MDD architectures versus controls.** Confusion matrices from cross-validated support vector machine classification are shown for three classification tasks: (A) healthy controls versus all MDD participants, (B) healthy controls versus architecture1, and (C) healthy controls versus architecture2. Each matrix displays the number of correctly and incorrectly classified participants based on out-of-fold predictions. These results complement ROC-based performance evaluation by illustrating threshold-dependent classification outcomes.

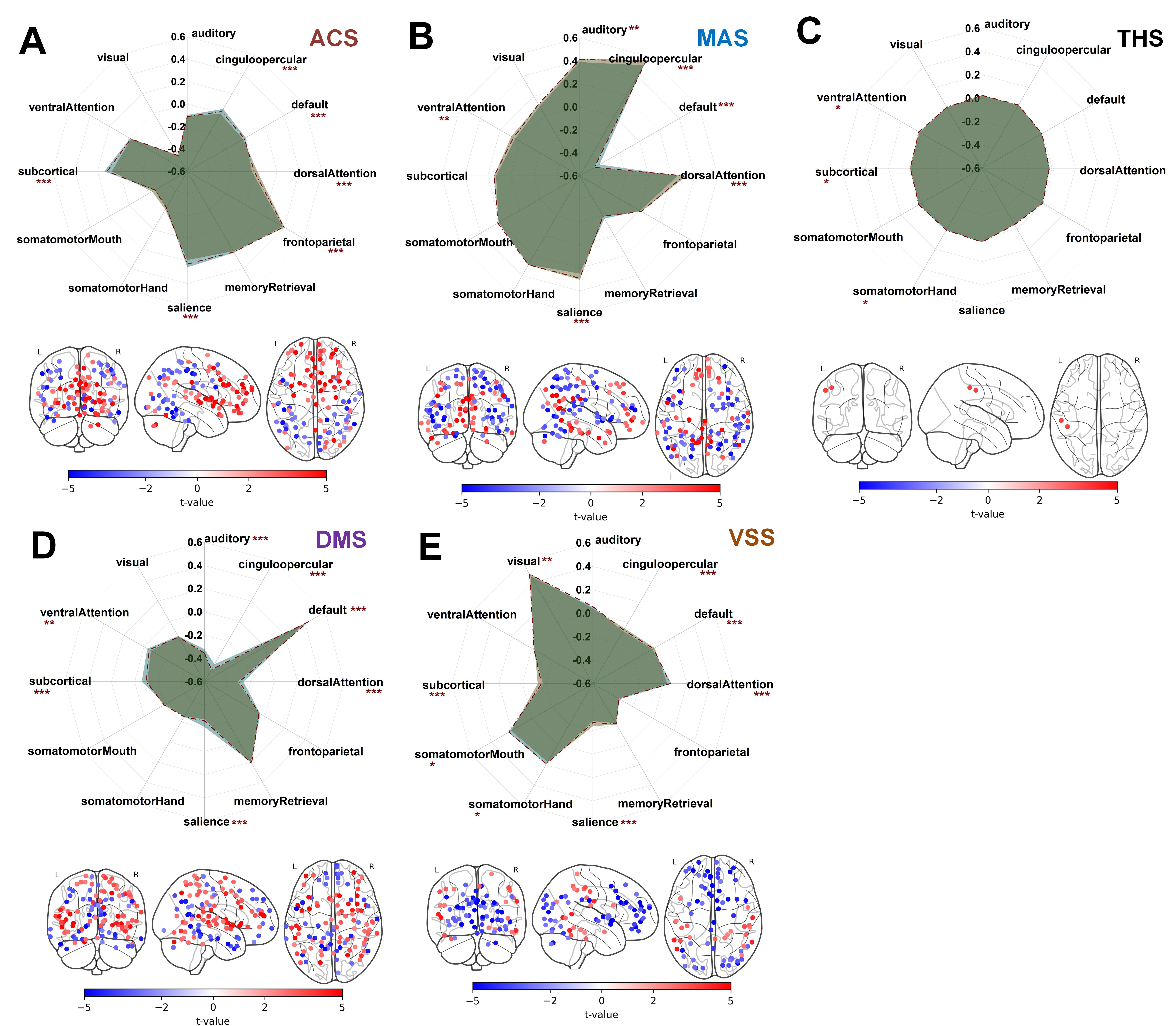

**Supplementary Figure S8. State-specific functional configurations of neurodynamic architectures.** Radar plots and brain maps illustrate the spatial functional configurations of the five recurrent brain states across the two neurodynamic architectures. (A–E) For each brain state, radar plots show the mean activity patterns across the 12 functional communities of the Power264 atlas in architecture 1 and architecture 2. Corresponding brain maps display ROI-level differences in state-specific activity between the two architectures, with color-coded t-values from voxel/community-level comparisons. Although some regional differences were observed within individual brain states, the overall spatial configurations of the five recurrent states were highly similar between the two architectures. These findings indicate that the identified neurodynamic architectures do not represent distinct functional state repertoires, but rather reflect different temporal organizations of a shared set of recurrent brain states.

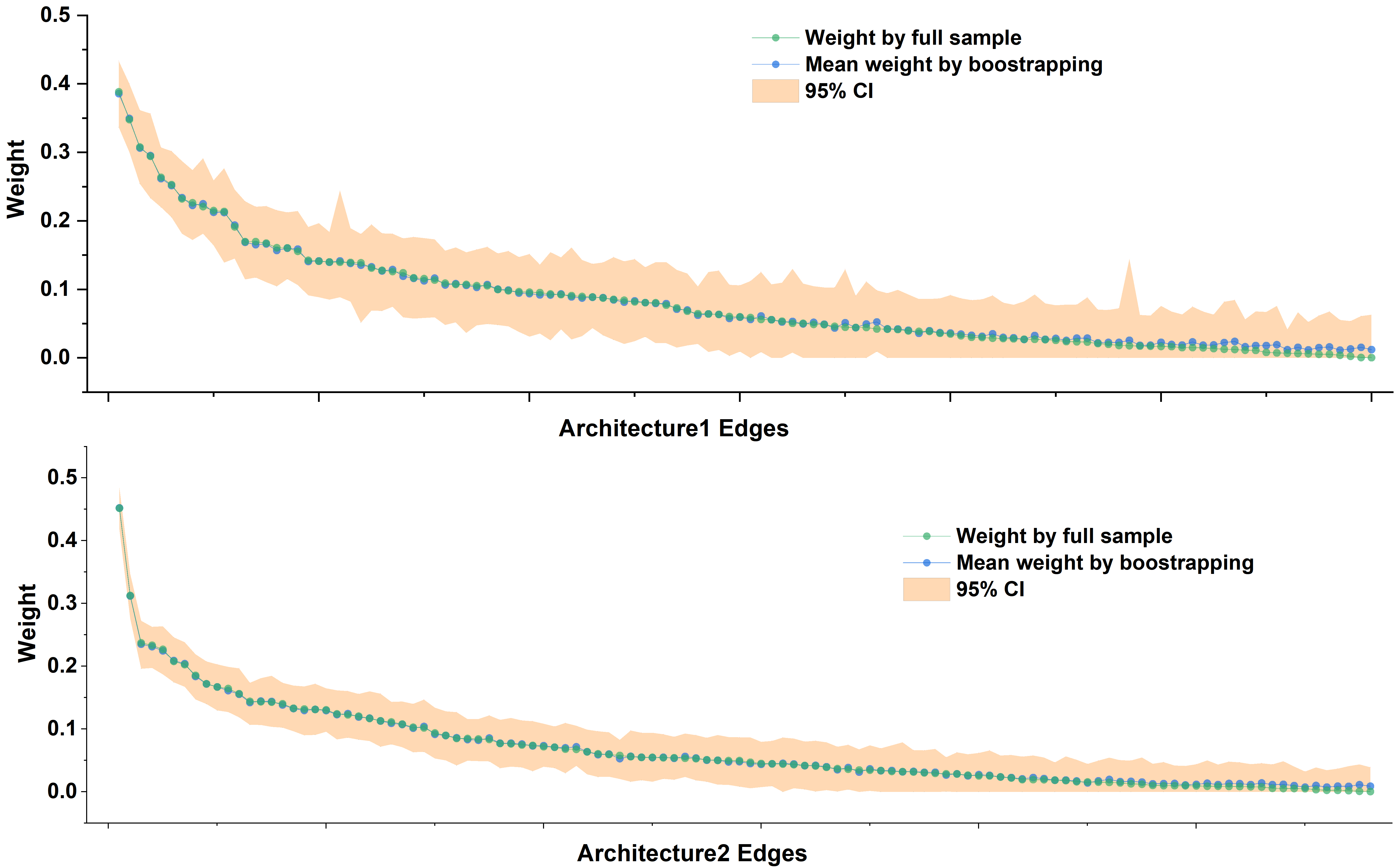

**Supplementary Figure S9. Bootstrap stability of symptom-network across neurodynamic architectures.** Bootstrap validation of architecture-specific symptom networks. Green points indicate the edge weights estimated from the full sample, whereas blue points represent the mean edge weights obtained across 1,000 bootstrap iterations. The orange shaded areas indicate the 95% confidence intervals derived from bootstrap resampling. Greater overlap between the full-sample edge weights and bootstrap-derived mean estimates indicates higher stability of the estimated edge weights. For both neurodynamic architectures, all observed edge weights from the original networks were contained within the bootstrap-derived 95% confidence intervals, supporting the robustness and reproducibility of the estimated symptom-network structures.

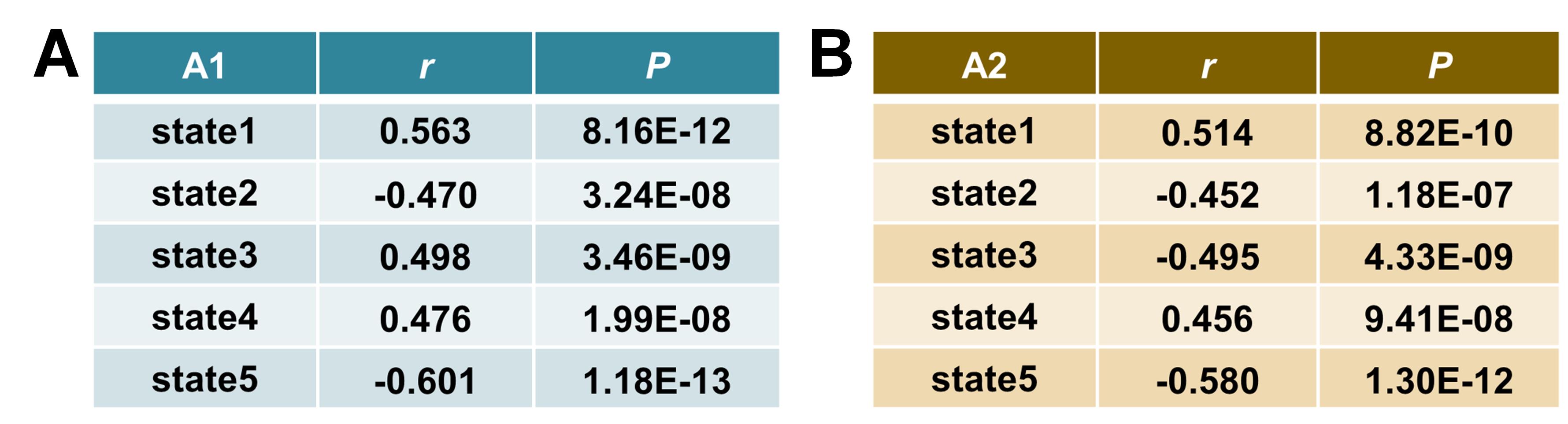

**Supplementary Figure S10. PLS1 associations with neuroimaging phenotypes. (A)** Correlations between PLS1 spatial scores and corresponding neuroimaging phenotypes across the five brain states in architecture 1; **(B)** Correlations between PLS1 spatial scores and corresponding neuroimaging phenotypes across the five brain states in architecture 2. Pearson correlation coefficients ($r$) quantify the spatial correspondence between PLS1-derived transcriptomic patterns and regional neuroimaging alterations. All architecture–state combinations showed significant associations between PLS1 spatial scores and imaging phenotypes after false discovery rate (FDR) correction, supporting the validity of PLS1 components for capturing transcriptomic patterns associated with state-specific neuroimaging variations. A1:architecture1; A2: architecture2; state1: ACS; state2: MAS; state3: THS; state4: DMS; state5: VSS;

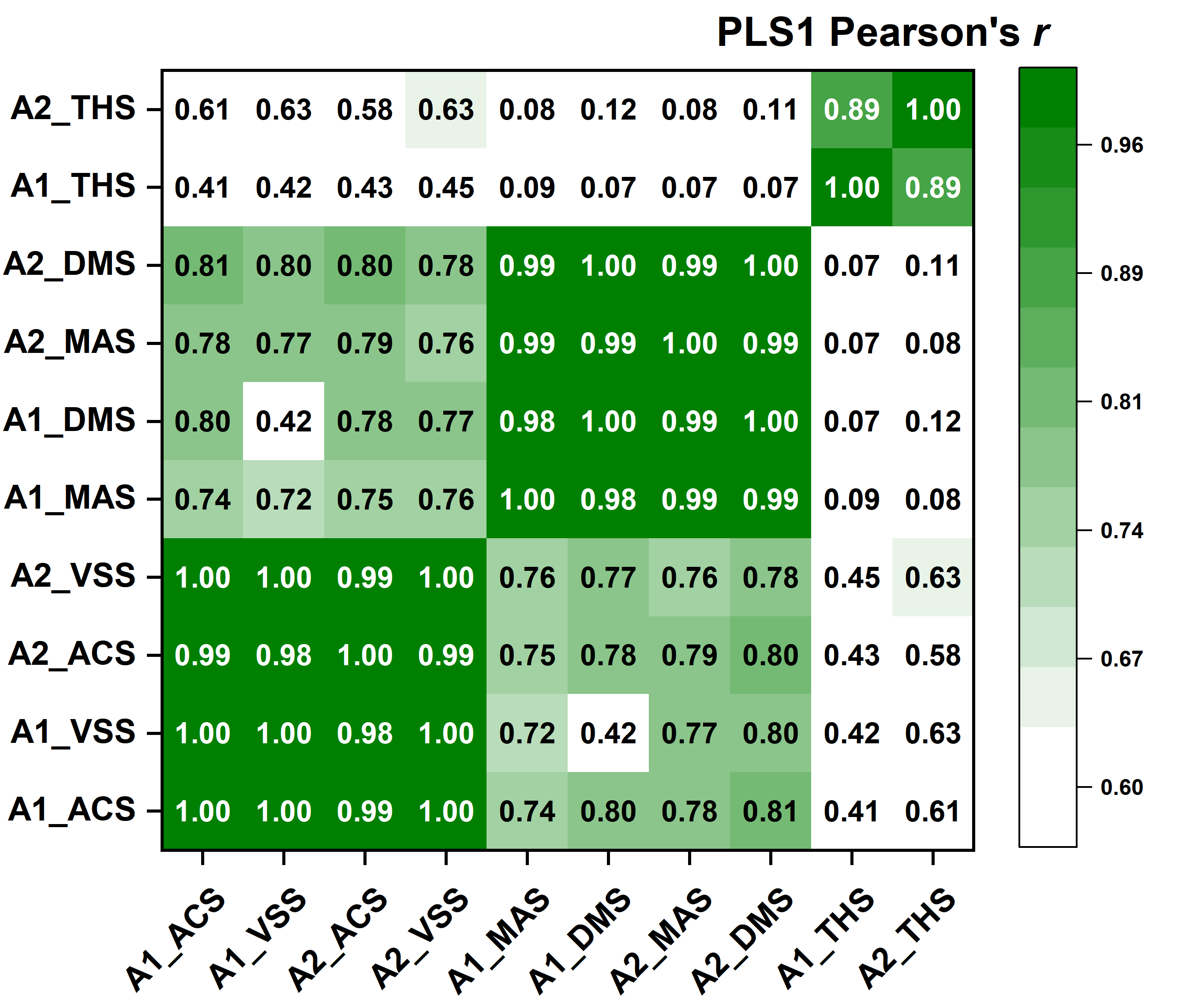

**Supplementary Figure S11. Similarity of PLS1 spatial score patterns across architecture–state combinations.** Pairwise similarity matrix of PLS1 spatial scores across the ten architecture–state transcriptomic models. Similarity was quantified using the absolute Pearson correlation coefficients between regional PLS1 score vectors derived from each architecture–state combination. The spatial transcriptomic similarity structure recapitulated the three transcriptomic families identified from PLS1 gene-loading profiles, with ACS/VSS-related states, MAS/DMS-related states, and THS-related states forming distinct clusters across architectures. These findings indicate that the identified transcriptomic state families were supported not only by gene-loading patterns but also by the corresponding spatial organization of PLS1-associated transcriptomic gradients. A1: Architecture 1; A2: Architecture 2;

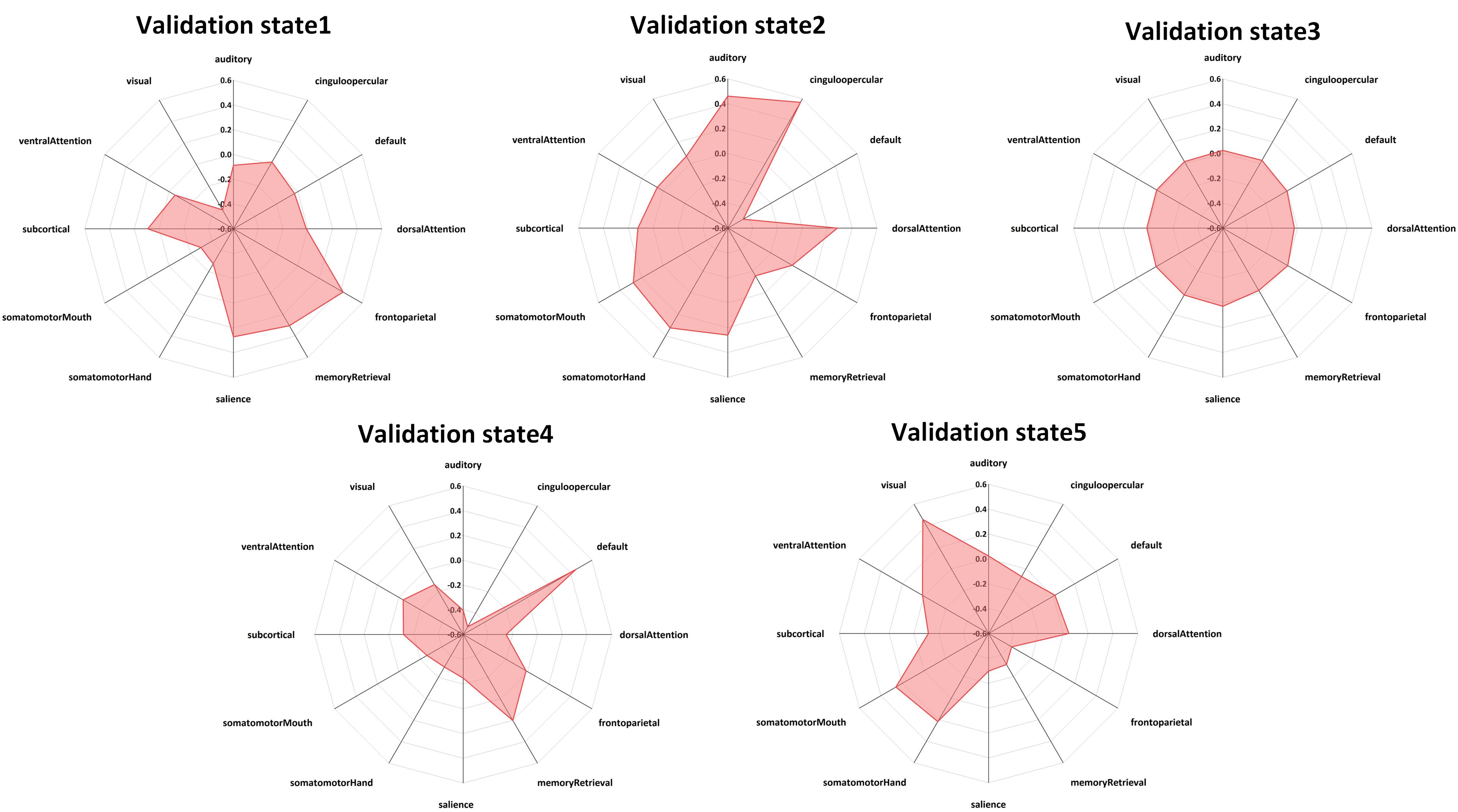

**Supplementary Figure S12. Brain-state configurations in the validation cohort.** Radar plots illustrate the network-level functional configurations of the five recurrent brain states inferred from the validation cohort. Each axis represents the mean activity pattern of a Power264 functional network within the corresponding brain state. The five states showed distinct spatial configurations, including functionally specialized patterns and a relatively balanced transition-oriented state, consistent with the brain-state organization identified in the discovery cohort. These results demonstrate the preservation of the recurrent brain-state repertoire in the independent validation dataset.

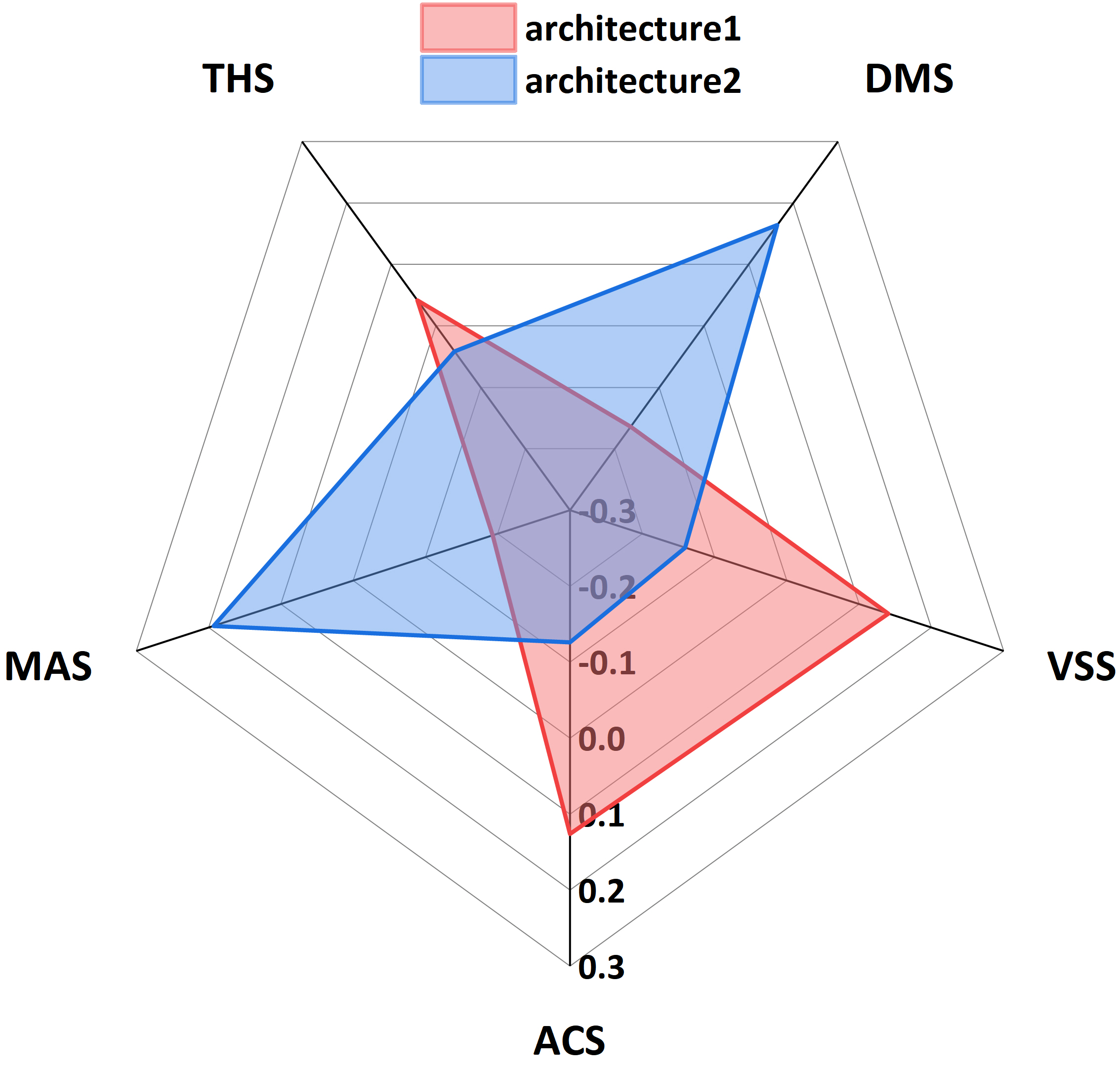

**Supplementary Figure S13. FO profiles of neurodynamic architectures in the independent validation cohort.** The radar plot illustrates the FO profiles of the two neurodynamic subgroups identified in the independent validation cohort across the five recurrent brain states. The two validation subgroups demonstrated complementary patterns of brain-state engagement, with one subgroup showing relatively increased occupancy of ACS and VSS and reduced occupancy of AVS and SMS, whereas the other exhibited the opposite pattern characterized by increased MAS and DMS occupancy and reduced CCS and SPS occupancy. The observed state-preference organization closely resembled the ACS/VSS-dominant and MAS/DMS-dominant neurodynamic architectures identified in the discovery cohort.

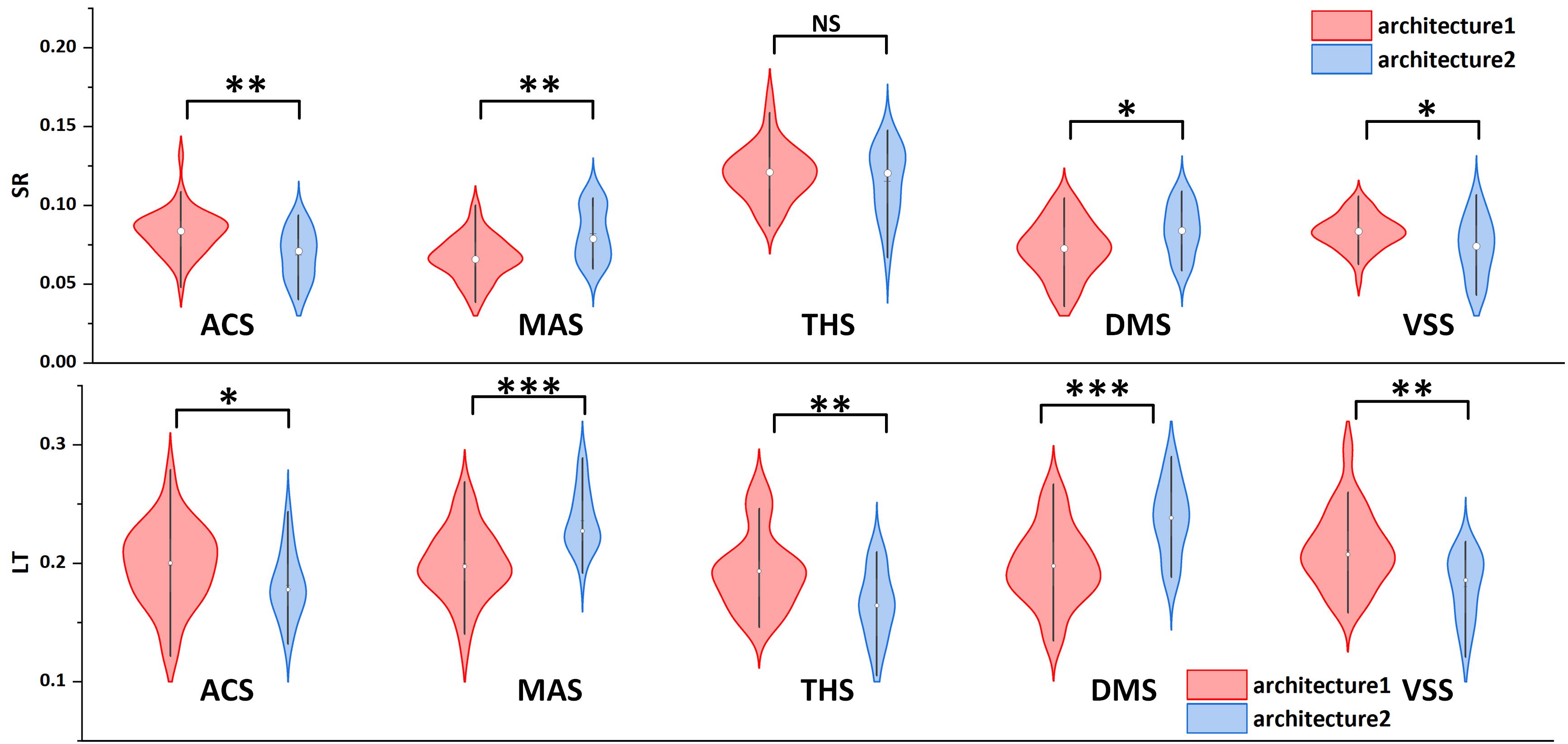

**Supplementary Figure S14. Temporal dynamics of validation neurodynamic architectures.** Violin plots show architecture differences in SR and LT across the five recurrent brain states in the independent validation cohort. Red and blue distributions represent validation architecture 1 and validation architecture 2, respectively. The two neurodynamic subgroups exhibited distinct temporal organization patterns across multiple brain states, with architecture-related differences in SR and LT showing concordant directions with those observed in the discovery cohort. Statistical significance was assessed using two-sample comparisons between validation architectures;*P < 0.05, **P < 0.01, ***P < 0.001; NS, not significant.

**SUPPLEMENTARY TABLES**

***Supplementary Table S1. Pairwise comparisons of all samples among the five recurrent brain states***

|  | *P_FO_* | *P_SR_* | *P_LT_* |
| --- | --- | --- | --- |
| state1 & state2 | 7.66E-08 | 2.32E-44 | 1.87E-17 |
| state1 & state3 | <1E-99 | <1E-99 | 1.42E-47 |
| state1 & state4 | 8.76E-22 | 0.2393 | 1.06E-41 |
| state1 & state5 | 0.0025 | 0.0001 | 0.5249 |
| state2 & state3 | <1E-99 | <1E-99 | <1E-99 |
| state2 & state4 | 6.23E-47 | 5.29E-36 | 1.29E-07 |
| state2 & state5 | 5.82E-16 | 8.88E-68 | 4.49E-19 |
| state3 & state4 | <1E-99 | <1E-99 | 2.6E-152 |
| state3 & state5 | <1E-99 | <1E-99 | 5.73E-42 |
| state4 & state5 | 1.68E-10 | 9.62E-07 | 1.72E-43 |

*Values represent two-sided P values from pairwise comparisons among the five recurrent brain states for fractional occupancy (FO), switching rate (SR), and lifetime (LT). P_FO_, P_SR_, and P_LT_ denote the corresponding P values for FO, SR, and LT, respectively.*

***Supplementary Table S2. Transition probability matrix of the five recurrent brain states under all samples***

|  | state1 | state2 | state3 | state4 | state5 |
| --- | --- | --- | --- | --- | --- |
| state1 | 0.643518 | 0.084122 | 0.137341 | 0.102602 | 0.032417 |
| state2 | 0.103103 | 0.661615 | 0.134547 | 0.004378 | 0.096357 |
| state3 | 0.089209 | 0.099952 | 0.615878 | 0.10122 | 0.093741 |
| state4 | 0.089077 | 0.004223 | 0.138678 | 0.670461 | 0.097561 |
| state5 | 0.033342 | 0.095346 | 0.125836 | 0.101964 | 0.643512 |

*Rows indicate origin states and columns indicate destination states. Values represent the mean transition probability across participants, with standard deviations shown in parentheses.*

***Supplementary Table S3. HC–MDD group comparison of brain-state features***

| Features | *P_state1_* | *P_state2_* | *P_state3_* | *P_state4_* | *P_state5_* |
| --- | --- | --- | --- | --- | --- |
| somatomotorHand | 0.0024 | 0.993529 | 0.7982 | 0.60417 | 0.010888 |
| somatomotorMouth | 0.00124 | 0.224829 | 0.7982 | 0.627793 | 0.12842 |
| cinguloopercular | 0.25998 | 0.993529 | 0.934848 | 0.383891 | 0.128909 |
| auditory | 0.005187 | 0.882889 | 0.7982 | 0.383891 | 0.000585 |
| default | 0.070803 | 0.007777 | 0.885951 | 0.001331 | 0.216894 |
| memoryRetrieval | 0.959273 | 0.367141 | 0.7982 | 0.910873 | 0.648679 |
| ventralAttention | 0.891324 | 0.003417 | 0.7982 | 0.041147 | 0.557943 |
| visual | 0.070803 | 0.005214 | 0.96491 | 0.031018 | 0.128909 |
| frontoparietal | 0.549862 | 0.428962 | 0.7982 | 0.60417 | 0.128909 |
| salience | 0.00124 | 0.271051 | 0.7982 | 0.943197 | 0.000585 |
| subcortical | 0.00124 | 0.882889 | 0.7982 | 0.467009 | 0.010675 |
| dorsalAttention | 0.011338 | 0.189694 | 0.7982 | 0.330593 | 0.00301 |
| FO | 0.554326 | 0.985512 | 0.921484 | 0.921484 | 0.921484 |
| SR | 0.921484 | 0.921484 | 0.495031 | 0.495031 | 0.495031 |
| LT | 0.921484 | 0.921484 | 0.921484 | 0.921484 | 0.921484 |

*P values from two-sample t-tests comparing HCs and MDD patients, corresponding to the t values shown in Fig. 3A. FDR correction was applied for multiple comparisons.*

***Supplementary Table S4. HAMD associations with brain-state features***

| Features | *P_state1_* | *P_state2_* | *P_state3_* | *P_state4_* | *P_state5_* |
| --- | --- | --- | --- | --- | --- |
| somatomotorHand | 0.273639 | 0.083244 | 0.694437 | 0.489635 | 0.409298 |
| somatomotorMouth | 0.247349 | 0.644269 | 0.644269 | 0.550004 | 0.694437 |
| cinguloopercular | 0.495646 | 0.002301 | 0.606258 | 0.034068 | 0.311597 |
| auditory | 0.928489 | 0.006334 | 0.704814 | 0.031691 | 0.694437 |
| default | 0.466385 | 0.031691 | 0.559251 | 0.311597 | 0.247349 |
| memoryRetrieval | 0.303688 | 0.550004 | 0.897883 | 0.53357 | 0.440211 |
| ventralAttention | 0.897883 | 0.960602 | 0.489635 | 0.833563 | 0.913403 |
| visual | 0.006334 | 0.489635 | 0.140241 | 0.524052 | 0.002301 |
| frontoparietal | 0.694437 | 0.303688 | 0.489635 | 0.303688 | 0.694437 |
| salience | 0.948901 | 0.822779 | 0.273639 | 0.833563 | 0.889799 |
| subcortical | 0.069673 | 0.644269 | 0.928489 | 0.750543 | 0.031691 |
| dorsalAttention | 0.190382 | 0.083244 | 0.802277 | 0.018097 | 0.031691 |
| FO | 0.116418 | 0.960602 | 0.006452 | 0.580888 | 0.226255 |
| SR | 0.15157 | 0.833563 | 0.031691 | 0.303688 | 0.15157 |
| LT | 0.694437 | 0.66472 | 0.691363 | 0.67367 | 0.694437 |

*P values from Spearman correlation analyses between brain-state features and HAMD scores in individuals with MDD, corresponding to the correlation coefficients shown in Fig. 3F. FDR correction was applied for multiple comparisons.*

***Supplementary Table S5. Statistical test of brain-state dynamic features between neurodynamic architectures***

| State-feature | *t* | *p* | *p_FDR_* |
| --- | --- | --- | --- |
| state1-FO | 26.00826 | 1.07E-110 | 2.28E-109 |
| state2-FO | -34.5263 | 1.60E-166 | 1.36E-164 |
| state3-FO | 4.979455 | 8.06E-07 | 2.36E-06 |
| state4-FO | -32.3837 | 4.82E-153 | 2.05E-151 |
| state5-FO | 30.39309 | 1.70E-140 | 4.82E-139 |
| state1-SR | 10.86454 | 1.24E-25 | 8.12E-25 |
| state2-SR | -14.8991 | 1.72E-44 | 1.63E-43 |
| state3-SR | 0.566409 | 0.571284 | 0.656204 |
| state4-SR | -13.0553 | 3.86E-35 | 2.98E-34 |
| state5-SR | 13.01724 | 3.24E-35 | 2.76E-34 |
| state1-LT | 19.95693 | 6.71E-72 | 7.13E-71 |
| state2-LT | -25.8014 | 9.00E-110 | 1.53E-108 |
| state3-LT | 4.226596 | 2.69E-05 | 6.52E-05 |
| state4-LT | -22.9404 | 8.10E-92 | 1.15E-90 |
| state5-LT | 21.52509 | 2.07E-82 | 2.51E-81 |

*Results of two-sample t-tests comparing fractional occupancy (FO), switching rate (SR), and lifetime (LT) between the two neurodynamic architectures. t values and corresponding uncorrected and FDR-adjusted P values are provided for each recurrent brain state. State1–5 correspond to CCS, AVS, THS, SMS, and SPS, respectively.*

***Supplementary Table S6. Architecture differences in brain-state transition probabilities***

| Edge | *t* | *p* | *p_FDR_* |
| --- | --- | --- | --- |
| S2_to_S2 | -21.7421 | 1.06E-79 | 2.64E-78 |
| S4_to_S4 | -19.2504 | 6.60E-66 | 8.25E-65 |
| S5_to_S5 | 16.78753 | 4.67E-56 | 3.90E-55 |
| S5_to_S2 | -15.6814 | 2.62E-50 | 1.64E-49 |
| S1_to_S1 | 15.73471 | 8.97E-50 | 4.49E-49 |
| S5_to_S4 | -14.7582 | 4.82E-45 | 2.01E-44 |
| S3_to_S2 | -14.4202 | 7.31E-43 | 2.61E-42 |
| S1_to_S4 | -13.8512 | 2.06E-40 | 6.45E-40 |
| S3_to_S4 | -12.9259 | 3.47E-35 | 9.63E-35 |
| S1_to_S2 | -12.4176 | 3.86E-33 | 9.65E-33 |
| S3_to_S5 | 12.54224 | 6.54E-33 | 1.49E-32 |
| S2_to_S5 | 11.60353 | 3.43E-28 | 7.14E-28 |
| S4_to_S1 | 10.27104 | 6.31E-23 | 1.21E-22 |
| S2_to_S1 | 9.412956 | 1.07E-19 | 1.90E-19 |
| S3_to_S1 | 9.040361 | 1.07E-18 | 1.78E-18 |
| S4_to_S5 | 8.660027 | 3.97E-17 | 6.20E-17 |
| S5_to_S1 | 7.923315 | 8.10E-15 | 1.19E-14 |
| S1_to_S5 | 6.121182 | 1.43E-09 | 1.99E-09 |
| S4_to_S3 | 5.139279 | 3.65E-07 | 4.80E-07 |
| S2_to_S4 | -4.01471 | 6.42E-05 | 8.03E-05 |
| S3_to_S3 | 3.590749 | 0.00035 | 0.000416 |
| S2_to_S3 | 2.638839 | 8.52E-03 | 0.00968 |
| S4_to_S2 | -1.77541 | 0.076215 | 0.082842 |
| S1_to_S3 | -0.9642 | 0.335216 | 0.349183 |
| S5_to_S3 | -0.232 | 0.816592 | 0.816592 |

*t values and P values from two-sample t-tests comparing directed transition probabilities between neurodynamic architectures. Edges indicate transitions from the origin state to the destination state.*

***Supplementary Table S7. neurodynamic architecture features associations with HAMD scores***

| Features | *r _architecture1_* | *p _architecture1_* | *p_FDR_* | *r _architecture2_* | *p _architecture2_* | *p_FDR_* |
| --- | --- | --- | --- | --- | --- | --- |
| state1-FO | -0.10961 | 0.044025 | 0.271518 | -0.06819 | 0.074289 | 0.353667 |
| state2-FO | -0.09858 | 0.070275 | 0.271518 | 0.021464 | 0.574642 | 0.871227 |
| state3-FO | 0.136637 | 0.011919 | 0.147026 | 0.102677 | 0.007114 | 0.141115 |
| state4-FO | -0.01381 | 0.800263 | 0.892134 | 0.044307 | 0.246491 | 0.698391 |
| state5-FO | 0.020689 | 0.704689 | 0.853149 | -0.08751 | 0.021887 | 0.177373 |
| state1-SR | -0.02029 | 0.710084 | 0.853149 | -0.08572 | 0.024752 | 0.177373 |
| state2-SR | -0.01495 | 0.78418 | 0.892134 | -0.0142 | 0.710499 | 0.888124 |
| state3-SR | 0.089088 | 0.102038 | 0.299078 | 0.09063 | 0.017582 | 0.177373 |
| state4-SR | 0.071266 | 0.191208 | 0.362108 | 0.041648 | 0.276015 | 0.728202 |
| state5-SR | -0.07403 | 0.174493 | 0.35314 | -0.05093 | 0.182764 | 0.535688 |
| state1-LT | -0.01955 | 0.72024 | 0.853149 | -0.02696 | 0.480798 | 0.851412 |
| state2-LT | -0.057 | 0.296089 | 0.493482 | 0.066482 | 0.081854 | 0.353667 |
| state3-LT | 0.039447 | 0.469789 | 0.726038 | 0.016433 | 0.667447 | 0.888124 |
| state4-LT | -0.11217 | 0.039297 | 0.271518 | -0.00845 | 0.825154 | 0.95712 |
| state5-LT | 0.145489 | 0.007381 | 0.147026 | -0.06531 | 0.087377 | 0.353667 |
| state1-none | 0.067237 | 0.217594 | 0.37746 | 0.0098 | 0.797781 | 0.95712 |
| state1-somatomotorHand | -0.08311 | 0.127289 | 0.318223 | -0.04035 | 0.291281 | 0.728202 |
| state1-somatomotorMouth | -0.11147 | 0.040546 | 0.271518 | -0.02833 | 0.458829 | 0.851412 |
| state1-cinguloopercular | 0.080802 | 0.138221 | 0.325665 | 0.018207 | 0.634055 | 0.888124 |
| state1-auditory | 0.006091 | 0.911172 | 0.933128 | 0.002493 | 0.948033 | 0.982717 |
| state1-default | 0.093412 | 0.08639 | 0.27271 | 0.017868 | 0.640367 | 0.888124 |
| state1-memoryRetrieval | -0.02414 | 0.658331 | 0.852279 | -0.06535 | 0.087194 | 0.353667 |
| state1-ventralAttention | 0.026051 | 0.63318 | 0.840942 | 0.00055 | 0.988525 | 0.996678 |
| state1-visual | -0.18097 | 0.000831 | 0.035309 | -0.07965 | 0.03701 | 0.24199 |
| state1-frontoparietal | 0.00639 | 0.906825 | 0.933128 | 0.021164 | 0.580017 | 0.871227 |
| state1-salience | 0.019178 | 0.725361 | 0.853149 | -0.0055 | 0.885716 | 0.962992 |
| state1-subcortical | 0.128577 | 0.018034 | 0.191609 | 0.06647 | 0.08191 | 0.353667 |
| state1-cerebellar | -0.08928 | 0.101312 | 0.299078 | -0.03269 | 0.392692 | 0.814117 |
| state1-dorsalAttention | -0.08801 | 0.106253 | 0.301051 | -0.05372 | 0.159901 | 0.515364 |
| state2-none | -0.09678 | 0.075582 | 0.27271 | -0.05402 | 0.157523 | 0.515364 |
| state2-somatomotorHand | 0.099137 | 0.068705 | 0.271518 | 0.066203 | 0.083147 | 0.353667 |
| state2-somatomotorMouth | 0.070397 | 0.196689 | 0.362108 | -0.00277 | 0.942206 | 0.982717 |
| state2-cinguloopercular | 0.207268 | 0.000124 | 0.009534 | 0.07458 | 0.050874 | 0.30888 |
| state2-auditory | 0.141575 | 0.009152 | 0.147026 | 0.095222 | 0.012591 | 0.177373 |
| state2-default | -0.0802 | 0.141189 | 0.325665 | -0.10071 | 0.008301 | 0.141115 |
| state2-memoryRetrieval | -0.11776 | 0.03043 | 0.271518 | 0.014529 | 0.704037 | 0.888124 |
| state2-ventralAttention | -0.08563 | 0.116099 | 0.303747 | 0.036553 | 0.339097 | 0.739526 |
| state2-visual | 0.093343 | 0.086626 | 0.27271 | 0.007064 | 0.853477 | 0.95712 |
| state2-frontoparietal | -0.1036 | 0.057078 | 0.271518 | -0.0226 | 0.554623 | 0.871227 |
| state2-salience | 0.006827 | 0.900481 | 0.933128 | 0.004539 | 0.905546 | 0.962992 |
| state2-subcortical | 0.067907 | 0.213031 | 0.377242 | -0.0045 | 0.906346 | 0.962992 |
| state2-cerebellar | -0.01089 | 0.841896 | 0.89787 | -0.01662 | 0.663946 | 0.888124 |
| state2-dorsalAttention | 0.011728 | 0.829897 | 0.89787 | 0.106088 | 0.005413 | 0.141115 |
| state3-none | -0.06989 | 0.199952 | 0.362108 | -0.00653 | 0.864368 | 0.95712 |
| state3-somatomotorHand | -0.02977 | 0.58542 | 0.83779 | -0.01539 | 0.687351 | 0.888124 |
| state3-somatomotorMouth | 0.029302 | 0.591382 | 0.83779 | 0.024081 | 0.528922 | 0.871227 |
| state3-cinguloopercular | -0.00335 | 0.95112 | 0.962443 | -0.04245 | 0.266819 | 0.728202 |
| state3-auditory | 0.013781 | 0.800708 | 0.892134 | -0.03624 | 0.34324 | 0.739526 |
| state3-default | 0.02388 | 0.66177 | 0.852279 | 0.035884 | 0.348012 | 0.739526 |
| state3-memoryRetrieval | -0.03422 | 0.530706 | 0.777759 | 0.02866 | 0.453596 | 0.851412 |
| state3-ventralAttention | 0.069846 | 0.200224 | 0.362108 | 0.015543 | 0.684455 | 0.888124 |
| state3-visual | -0.02056 | 0.70647 | 0.853149 | -0.08957 | 0.018956 | 0.177373 |
| state3-frontoparietal | 0.100086 | 0.066082 | 0.271518 | 0.006404 | 0.867039 | 0.95712 |
| state3-salience | 0.047378 | 0.385231 | 0.617823 | 0.051663 | 0.176515 | 0.535688 |
| state3-subcortical | 0.085206 | 0.117925 | 0.303747 | -0.03701 | 0.33315 | 0.739526 |
| state3-cerebellar | 0.039945 | 0.464199 | 0.726038 | -0.02932 | 0.443189 | 0.851412 |
| state3-dorsalAttention | 0.010669 | 0.845054 | 0.89787 | 0.017815 | 0.641366 | 0.888124 |
| state4-none | 0.098841 | 0.069542 | 0.271518 | 0.021565 | 0.572847 | 0.871227 |
| state4-somatomotorHand | -0.05283 | 0.33284 | 0.544066 | -0.02812 | 0.462202 | 0.851412 |
| state4-somatomotorMouth | -0.07788 | 0.15308 | 0.325665 | -0.00868 | 0.820416 | 0.95712 |
| state4-cinguloopercular | -0.14019 | 0.009866 | 0.147026 | -0.05531 | 0.147883 | 0.515364 |
| state4-auditory | -0.0941 | 0.084093 | 0.27271 | -0.08567 | 0.024842 | 0.177373 |
| state4-default | 0.018644 | 0.732705 | 0.853149 | 0.064073 | 0.093576 | 0.361544 |
| state4-memoryRetrieval | 0.101318 | 0.062802 | 0.271518 | -0.00665 | 0.861959 | 0.95712 |
| state4-ventralAttention | 0.077134 | 0.157085 | 0.325665 | -0.05866 | 0.12482 | 0.461292 |
| state4-visual | -0.07754 | 0.154916 | 0.325665 | -0.01177 | 0.758369 | 0.934223 |
| state4-frontoparietal | 0.093888 | 0.084794 | 0.27271 | 0.02737 | 0.474175 | 0.851412 |
| state4-salience | 0.026749 | 0.624103 | 0.840942 | -0.02334 | 0.541646 | 0.871227 |
| state4-subcortical | -0.07094 | 0.193257 | 0.362108 | 0.022367 | 0.558659 | 0.871227 |
| state4-cerebellar | 0.001116 | 0.983698 | 0.983698 | 0.016521 | 0.665777 | 0.888124 |
| state4-dorsalAttention | -0.02118 | 0.697995 | 0.853149 | -0.14131 | 0.000205 | 0.017403 |
| state5-none | -0.02612 | 0.632325 | 0.840942 | -0.00752 | 0.844154 | 0.95712 |
| state5-somatomotorHand | 0.085915 | 0.114891 | 0.303747 | 0.020928 | 0.584234 | 0.871227 |
| state5-somatomotorMouth | 0.058462 | 0.283832 | 0.482514 | 0.000159 | 0.996678 | 0.996678 |
| state5-cinguloopercular | -0.09995 | 0.066462 | 0.271518 | -0.02227 | 0.560444 | 0.871227 |
| state5-auditory | -0.02643 | 0.62825 | 0.840942 | -0.01429 | 0.708702 | 0.888124 |
| state5-default | -0.10024 | 0.065657 | 0.271518 | -0.03603 | 0.346013 | 0.739526 |
| state5-memoryRetrieval | 0.037171 | 0.495817 | 0.750717 | 0.040891 | 0.28485 | 0.728202 |
| state5-ventralAttention | -0.07788 | 0.153095 | 0.325665 | 0.030201 | 0.429673 | 0.851412 |
| state5-visual | 0.116194 | 0.032718 | 0.271518 | 0.128899 | 0.000714 | 0.030356 |
| state5-frontoparietal | 0.013255 | 0.808168 | 0.892134 | -0.03735 | 0.328644 | 0.739526 |
| state5-salience | -0.03652 | 0.503422 | 0.750717 | -0.0003 | 0.993761 | 0.996678 |
| state5-subcortical | -0.13634 | 0.012108 | 0.147026 | -0.07335 | 0.054837 | 0.310743 |
| state5-cerebellar | 0.079185 | 0.14631 | 0.325665 | 0.053234 | 0.163704 | 0.515364 |
| state5-dorsalAttention | 0.09952 | 0.067636 | 0.271518 | 0.085552 | 0.025041 | 0.177373 |

***Supplementary Table S8. Symptom network centrality profiles of neurodynamic architectures***

| label | S1_DC_ | S1_AS_ | S1_CLU_ | S1_CLO_ | S1_BET_ | S2_DC_ | S2_AS_ | S2_CLU_ | S2_CLO_ | S2_BET_ |
| --- | --- | --- | --- | --- | --- | --- | --- | --- | --- | --- |
| H1 | -0.693 | 2.272 | 2.156 | -0.736 | -0.841 | 0.092 | 1.900 | 0.464 | 0.026 | -0.841 |
| H2 | 0.878 | 0.465 | 0.947 | 0.878 | -0.841 | 0.092 | -0.828 | -0.893 | 0.026 | 0.368 |
| H3 | 1.663 | 0.746 | 1.284 | 1.836 | -0.841 | 0.092 | -0.603 | -0.802 | 0.026 | -0.841 |
| H4 | 0.878 | 0.243 | 0.789 | 0.878 | -0.628 | -0.693 | -0.577 | -1.113 | -0.736 | 0.653 |
| H5 | 0.878 | 0.128 | 0.487 | 0.878 | -0.201 | 0.878 | 0.485 | 0.057 | 0.878 | -0.841 |
| H6 | 1.663 | 0.113 | 0.455 | 1.836 | 1.862 | -0.693 | -0.618 | -0.921 | -0.736 | 0.226 |
| H7 | 0.092 | 0.221 | 0.267 | 0.026 | 1.008 | 0.092 | 0.403 | -0.557 | 0.026 | -0.556 |
| H8 | 0.092 | 1.870 | 2.153 | 0.026 | -0.485 | 1.663 | -0.720 | -0.838 | 1.836 | 0.084 |
| H9 | -0.693 | -0.159 | 0.571 | -0.736 | -0.556 | -0.693 | 0.374 | -0.066 | -0.736 | -0.699 |
| H10 | 0.878 | 1.681 | 1.291 | 0.878 | 1.151 | 0.092 | 0.391 | -0.177 | 0.026 | -0.414 |
| H11 | 0.878 | 0.085 | 0.883 | 0.878 | -0.841 | 0.092 | 0.115 | -0.591 | 0.026 | -0.770 |
| H12 | -0.693 | 0.580 | 0.876 | -0.736 | -0.556 | -2.263 | 0.056 | -0.607 | -2.042 | -0.841 |
| H13 | -1.478 | -0.283 | 0.523 | -1.422 | -0.841 | -1.478 | 0.003 | -0.440 | -1.422 | 0.297 |
| H14 | 0.092 | -0.746 | -0.247 | 0.026 | 1.577 | 0.878 | -1.508 | -1.429 | 0.878 | 0.653 |
| H15 | 0.092 | 0.268 | 0.615 | 0.026 | 0.084 | 0.092 | -1.609 | -1.275 | 0.026 | 1.079 |
| H16 | -0.693 | 0.044 | 0.398 | -0.736 | -0.699 | 0.878 | -1.077 | -1.479 | 0.878 | 1.435 |
| H17 | -0.693 | -1.781 | -1.075 | -0.736 | 3.213 | -2.263 | -1.932 | -1.705 | -2.042 | -0.556 |

*Standardized centrality measures for HAMD symptom networks in architecture1 and architecture2. S1 and S2 represent architecture1 and architecture2. DC, AS, CLU, CLO, and BET represent degree centrality, average strength, clustering coefficient, closeness centrality, and betweenness centrality, respectively. H1–H17 correspond to individual HAMD symptoms.*

***Supplementary Table S9. PLS component variance explained in neurodynamic architecture–states***

| *component* | *S1_state1_* | *S1_state2_* | *S1_state3_* | *S1_state4_* | *S1_state5_* | *S2_state1_* | *S2_state2_* | *S2_state3_* | *S2_state4_* | *S2_state5_* |
| --- | --- | --- | --- | --- | --- | --- | --- | --- | --- | --- |
| 1 | 0.317 | 0.221 | 0.248 | 0.227 | 0.362 | 0.264 | 0.205 | 0.245 | 0.208 | 0.337 |
| 2 | 0.123 | 0.106 | 0.096 | 0.108 | 0.116 | 0.127 | 0.111 | 0.134 | 0.111 | 0.115 |
| 3 | 0.115 | 0.117 | 0.163 | 0.117 | 0.102 | 0.127 | 0.112 | 0.148 | 0.113 | 0.102 |
| 4 | 0.087 | 0.124 | 0.077 | 0.116 | 0.087 | 0.083 | 0.136 | 0.056 | 0.129 | 0.086 |
| 5 | 0.152 | 0.154 | 0.097 | 0.160 | 0.128 | 0.171 | 0.163 | 0.087 | 0.166 | 0.127 |
| 6 | 0.048 | 0.084 | 0.091 | 0.086 | 0.052 | 0.063 | 0.083 | 0.101 | 0.088 | 0.063 |
| 7 | 0.035 | 0.055 | 0.051 | 0.047 | 0.036 | 0.027 | 0.052 | 0.049 | 0.047 | 0.039 |
| 8 | 0.028 | 0.021 | 0.023 | 0.021 | 0.034 | 0.029 | 0.021 | 0.027 | 0.021 | 0.030 |
| 9 | 0.011 | 0.037 | 0.022 | 0.039 | 0.011 | 0.015 | 0.039 | 0.024 | 0.042 | 0.014 |
| 10 | 0.012 | 0.012 | 0.017 | 0.012 | 0.009 | 0.012 | 0.012 | 0.015 | 0.012 | 0.011 |
| 11 | 0.010 | 0.016 | 0.031 | 0.014 | 0.007 | 0.013 | 0.015 | 0.025 | 0.014 | 0.008 |
| 12 | 0.008 | 0.010 | 0.013 | 0.009 | 0.007 | 0.012 | 0.010 | 0.011 | 0.010 | 0.010 |
| 13 | 0.007 | 0.008 | 0.010 | 0.007 | 0.009 | 0.006 | 0.008 | 0.014 | 0.006 | 0.011 |
| 14 | 0.006 | 0.004 | 0.007 | 0.006 | 0.006 | 0.007 | 0.004 | 0.005 | 0.005 | 0.006 |
| 15 | 0.006 | 0.004 | 0.010 | 0.005 | 0.005 | 0.007 | 0.004 | 0.008 | 0.004 | 0.006 |
| 16 | 0.004 | 0.004 | 0.005 | 0.004 | 0.003 | 0.005 | 0.004 | 0.005 | 0.004 | 0.004 |
| 17 | 0.005 | 0.004 | 0.009 | 0.004 | 0.004 | 0.005 | 0.003 | 0.006 | 0.003 | 0.005 |
| 18 | 0.004 | 0.004 | 0.004 | 0.004 | 0.003 | 0.005 | 0.004 | 0.005 | 0.004 | 0.004 |
| 19 | 0.003 | 0.004 | 0.005 | 0.003 | 0.003 | 0.004 | 0.003 | 0.005 | 0.003 | 0.004 |
| 20 | 0.003 | 0.002 | 0.004 | 0.003 | 0.003 | 0.004 | 0.002 | 0.006 | 0.003 | 0.003 |

*Proportion of imaging-related covariance explained by each PLS component for each architecture–state-specific transcriptomic model. S1 and S2 represent architecture1 and architecture2. PLS1 represents the first latent component and accounts for the largest proportion of explained covariance across architecture–state combinations.*

***Supplementary Table S10. Spatial correlations between ROI activity patterns and PLS1 scores***

|  | *r* | *p* |
| --- | --- | --- |
| S1_state1_ | 0.563111 | 8.16E-12 |
| S1_state2_ | -0.46985 | 3.24E-08 |
| S1_state3_ | 0.497928 | 3.46E-09 |
| S1_state4_ | 0.476184 | 1.99E-08 |
| S1_state5_ | -0.60158 | 1.18E-13 |
| S2_state1_ | 0.51391 | 8.82E-10 |
| S2_state2_ | -0.45245 | 1.18E-07 |
| S2_state3_ | -0.49522 | 4.33E-09 |
| S2_state4_ | 0.455526 | 9.41E-08 |
| S2_state5_ | -0.58042 | 1.30E-12 |

*Pearson correlation analyses between state-specific regional imaging patterns (ROI activity) and corresponding PLS1 spatial scores derived from architecture–state-specific transcriptomic models. S1 and S2 represent architecture1 and architecture2.*

***Supplementary Table S11.*** ***Enrichment analysis results of transcriptomic Family 3***

| *Pathway* | *Count* | *%* | *Log10(P)* | *Log10(q)* | *-Log10(p)* | *-Log10(q)* |
| --- | --- | --- | --- | --- | --- | --- |
| GO: trans-synaptic signaling | 7 | 11.11 | -4.37 | -0.09 | 4.37 | 0.09 |
| GO: postsynapse | 8 | 12.7 | -3.72 | 0 | 3.72 | 0 |
| GO: regulation of calcium-mediated signaling | 3 | 4.76 | -2.93 | 0 | 2.93 | 0 |
| GO: regulation of lymphocyte differentiation | 4 | 6.35 | -2.8 | 0 | 2.8 | 0 |
| GO: small GTPase-mediated signal transduction | 4 | 6.35 | -2.59 | 0 | 2.59 | 0 |
| GO: sex differentiation | 4 | 6.35 | -2.54 | 0 | 2.54 | 0 |
| GO: signaling adaptor activity | 3 | 4.76 | -2.52 | 0 | 2.52 | 0 |
| GO: calcium ion binding | 6 | 9.52 | -2.45 | 0 | 2.45 | 0 |
| GO: basal plasma membrane | 4 | 6.35 | -2.43 | 0 | 2.43 | 0 |
| GO: cellular response to steroid hormone stimulus | 3 | 4.76 | -2.36 | 0 | 2.36 | 0 |
| GO: peptidase complex | 3 | 4.76 | -2.33 | 0 | 2.33 | 0 |
| GO: extrinsic component of membrane | 3 | 4.76 | -2.29 | 0 | 2.29 | 0 |
| GO: protein kinase activity | 5 | 7.94 | -2.15 | 0 | 2.15 | 0 |
| GO: regulation of reproductive process | 3 | 4.76 | -2.11 | 0 | 2.11 | 0 |
| GO: glycoprotein biosynthetic process | 3 | 4.76 | -1.71 | 0 | 1.71 | 0 |
| GO: DNA metabolic process | 5 | 7.94 | -1.64 | 0 | 1.64 | 0 |
| GO: motile cilium | 3 | 4.76 | -1.5 | 0 | 1.5 | 0 |
| GO: small molecule catabolic process | 3 | 4.76 | -1.43 | 0 | 1.43 | 0 |
| GO: positive regulation of cell projection organization | 3 | 4.76 | -1.41 | 0 | 1.41 | 0 |
| GO: protein heterodimerization activity | 3 | 4.76 | -1.37 | 0 | 1.37 | 0 |

*The table lists the top 20 ranked pathways based on nominal enrichment significance. Count, percentage of overlapping genes, nominal P values, and FDR-adjusted q values are provided. No pathway reached significance after false discovery rate correction.*

***Supplementary Table S12.*** ***Jaccard similarity of PLS1-associated genes across architecture–state combinations***

|  | S1_CCS | S1_SPS | S2_CCS | S2_SPS | S1_AVS | S2_AVS | S1_SMS | S2_SMS | S1_THS | S2_THS |
| --- | --- | --- | --- | --- | --- | --- | --- | --- | --- | --- |
| S1_CCS | 1.00 | 0.82 | 0.74 | 0.83 | 0.09 | 0.09 | 0.12 | 0.10 | 0.01 | 0.02 |
| S1_SPS | 0.82 | 1.00 | 0.62 | 0.85 | 0.07 | 0.08 | 0.11 | 0.09 | 0.01 | 0.02 |
| S2_CCS | 0.74 | 0.62 | 1.00 | 0.69 | 0.09 | 0.09 | 0.11 | 0.10 | 0.00 | 0.01 |
| S2_SPS | 0.83 | 0.85 | 0.69 | 1.00 | 0.07 | 0.07 | 0.10 | 0.09 | 0.01 | 0.02 |
| S1_AVS | 0.09 | 0.07 | 0.09 | 0.07 | 1.00 | 0.77 | 0.64 | 0.67 | 0.02 | 0.01 |
| S1_SMS | 0.12 | 0.11 | 0.11 | 0.10 | 0.64 | 0.65 | 1.00 | 0.78 | 0.01 | 0.00 |
| S2_AVS | 0.09 | 0.08 | 0.09 | 0.07 | 0.77 | 1.00 | 0.65 | 0.79 | 0.02 | 0.00 |
| S2_SMS | 0.10 | 0.09 | 0.10 | 0.09 | 0.67 | 0.79 | 0.78 | 1.00 | 0.01 | 0.00 |
| S1_THS | 0.01 | 0.01 | 0.00 | 0.01 | 0.02 | 0.02 | 0.01 | 0.01 | 1.00 | 0.38 |
| S2_THS | 0.02 | 0.02 | 0.01 | 0.02 | 0.01 | 0.00 | 0.00 | 0.00 | 0.38 | 1.00 |

*The Jaccard coefficient was defined as the size of the intersection divided by the size of the union between two gene sets, reflecting the proportion of shared genes relative to the total number of unique genes. Higher values indicate greater overlap of PLS1-associated genes between transcriptomic models.*

***Supplementary Table S13.*** ***Overlap coefficient of PLS1-associated genes across architecture–state combinations***

|  | S1_CCS | S1_SPS | S2_CCS | S2_SPS | S1_AVS | S2_AVS | S1_SMS | S2_SMS | S1_THS | S2_THS |
| --- | --- | --- | --- | --- | --- | --- | --- | --- | --- | --- |
| S1_CCS | 1.00 | 0.98 | 0.93 | 0.96 | 0.42 | 0.47 | 0.46 | 0.48 | 0.08 | 0.18 |
| S1_SPS | 0.98 | 1.00 | 0.90 | 0.93 | 0.42 | 0.48 | 0.45 | 0.49 | 0.11 | 0.26 |
| S2_CCS | 0.93 | 0.90 | 1.00 | 0.95 | 0.37 | 0.41 | 0.41 | 0.42 | 0.04 | 0.11 |
| S2_SPS | 0.96 | 0.93 | 0.95 | 1.00 | 0.40 | 0.45 | 0.43 | 0.46 | 0.13 | 0.21 |
| S1_AVS | 0.42 | 0.42 | 0.37 | 0.40 | 1.00 | 0.88 | 0.90 | 0.81 | 0.08 | 0.02 |
| S1_SMS | 0.46 | 0.45 | 0.41 | 0.43 | 0.90 | 0.92 | 1.00 | 0.92 | 0.07 | 0.02 |
| S2_AVS | 0.47 | 0.48 | 0.41 | 0.45 | 0.88 | 1.00 | 0.92 | 0.98 | 0.06 | 0.02 |
| S2_SMS | 0.48 | 0.49 | 0.42 | 0.46 | 0.81 | 0.98 | 0.92 | 1.00 | 0.06 | 0.02 |
| S1_THS | 0.08 | 0.11 | 0.04 | 0.13 | 0.08 | 0.06 | 0.07 | 0.06 | 1.00 | 0.64 |
| S2_THS | 0.18 | 0.26 | 0.11 | 0.21 | 0.02 | 0.02 | 0.02 | 0.02 | 0.64 | 1.00 |

*The overlap coefficient was defined as the size of the intersection divided by the size of the smaller gene set, quantifying the degree to which genes from one model were contained within another model. Higher values indicate greater inclusion of shared PLS1-associated genes between transcriptomic models.*

***Supplementary Table S14. Sparse principal component loadings of brain-state features***

|  | sPC1 | sPC2 | sPC3 | sPC4 | sPC5 |
| --- | --- | --- | --- | --- | --- |
| state1-FO | -0.33414 | 0.000467 | 0 | 0 | 0 |
| state2-FO | 0.367829 | 0 | 0 | 0 | 0 |
| state3-FO | -0.00495 | -0.12018 | 0 | 0 | 0 |
| state4-FO | 0.370141 | 0 | 0 | 0 | 0 |
| state5-FO | -0.3497 | 0 | 0 | 0 | 0 |
| state1-SR | -0.13683 | 0 | 0 | 0 | 0 |
| state2-SR | 0.224489 | 0 | 0 | 0 | 0 |
| state3-SR | 0.041529 | 0 | 0 | 0 | 0 |
| state4-SR | 0.192932 | 0 | 0 | 0 | 0 |
| state5-SR | -0.15703 | 0 | 0 | 0 | 0 |
| state1-LT | -0.23619 | 0.069055 | 0 | 0 | 0 |
| state2-LT | 0.274598 | 0 | 0 | 0 | 0 |
| state3-LT | -0.04081 | -0.13477 | 0 | 0 | 0 |
| state4-LT | 0.302969 | 0 | 0 | 0 | 0 |
| state5-LT | -0.24729 | 0 | -0.01964 | 0 | 0 |
| state1-somatomotorHand | 0 | -0.40219 | 0 | 0 | 0 |
| state1-somatomotorMouth | 0 | -0.38759 | 0 | 0 | 0 |
| state1-cinguloopercular | 0 | -0.06234 | -0.28106 | 0 | 0 |
| state1-auditory | 0 | -0.29835 | -0.04898 | 0 | 0 |
| state1-default | 0 | 0 | -0.16538 | -0.19202 | 0 |
| state1-memoryRetrieval | 0 | 0.123513 | 0 | 0 | 0 |
| state1-ventralAttention | 0 | 0 | -0.03904 | 0 | 0 |
| state1-visual | 0 | 0 | 0.302791 | -0.11332 | 0 |
| state1-frontoparietal | 0 | 0.128762 | 0 | 0.185691 | 0 |
| state1-salience | -0.01363 | 0.061226 | -0.28513 | 0 | 0 |
| state1-subcortical | 0 | 0 | -0.35205 | 0 | 0 |
| state1-dorsalAttention | 0 | 0 | 0.305542 | 0.009556 | 0 |
| state2-somatomotorHand | 0 | 0 | 0 | 0 | -0.37284 |
| state2-somatomotorMouth | 0 | 0 | 0 | 0 | -0.39004 |
| state2-cinguloopercular | 0 | 0 | 0 | 0.389287 | -0.14902 |
| state2-auditory | 0 | 0 | 0 | 0.106659 | -0.38914 |
| state2-default | -0.18455 | -0.14832 | 0.037145 | 0 | 0 |
| state2-memoryRetrieval | 0 | -0.0275 | 0 | 0 | 0 |
| state2-ventralAttention | 0 | 0 | 0 | 0.046805 | 0 |
| state2-visual | 0 | 0.097671 | -0.02854 | 0 | 0 |
| state2-frontoparietal | 0 | 0 | 0 | 0 | 0.155646 |
| state2-salience | 0 | 0 | 0 | 0.392737 | 0 |
| state2-subcortical | 0 | 0 | 0 | 0.31869 | 0 |
| state2-dorsalAttention | 0.101273 | 0 | 0 | 0 | 0 |
| state3-somatomotorHand | 0 | 0 | 0 | 0 | 0 |
| state3-somatomotorMouth | 0 | 0 | 0 | 0 | 0 |
| state3-cinguloopercular | 0 | 0 | 0 | 0 | 0 |
| state3-auditory | 0 | 0 | 0 | 0 | 0 |
| state3-default | 0 | 0 | 0 | 0 | 0 |
| state3-memoryRetrieval | 0 | 0 | 0 | 0 | 0 |
| state3-ventralAttention | 0 | 0 | 0 | 0 | 0 |
| state3-visual | 0 | 0 | 0 | 0 | 0 |
| state3-frontoparietal | 0 | 0 | 0 | 0 | 0 |
| state3-salience | 0 | 0 | 0 | 0 | 0 |
| state3-subcortical | 0 | 0 | 0 | 0 | 0 |
| state3-dorsalAttention | 0 | 0 | 0 | 0 | 0 |
| state4-somatomotorHand | 0 | 0 | 0 | 0 | 0.404316 |
| state4-somatomotorMouth | 0 | 0 | 0 | 0 | 0.390544 |
| state4-cinguloopercular | 0 | 0 | 0 | -0.39193 | 0.15568 |
| state4-auditory | 0 | 0 | 0 | -0.13938 | 0.381129 |
| state4-default | 0.163144 | 0.134522 | -0.06511 | 0 | 0 |
| state4-memoryRetrieval | 0 | 0.011388 | -0.00781 | 0 | 0 |
| state4-ventralAttention | 0 | 0 | 0 | -0.056 | 0 |
| state4-visual | 0 | -0.04062 | 0.13735 | 0 | 0 |
| state4-frontoparietal | 0 | 0 | 0 | 0 | -0.15972 |
| state4-salience | 0 | 0 | 0 | -0.38217 | 0 |
| state4-subcortical | 0 | 0 | -0.02328 | -0.31362 | 0 |
| state4-dorsalAttention | -0.06401 | 0 | 0 | 0 | 0 |
| state5-somatomotorHand | 0 | 0.411905 | 0 | 0 | 0 |
| state5-somatomotorMouth | 0 | 0.400298 | 0 | 0 | 0 |
| state5-cinguloopercular | 0 | 0.0814 | 0.267365 | 0 | 0 |
| state5-auditory | 0 | 0.308122 | 0.037388 | 0 | 0 |
| state5-default | 0 | 0 | 0.190989 | 0.190521 | 0 |
| state5-memoryRetrieval | 0 | -0.10971 | 0 | 0 | 0 |
| state5-ventralAttention | 0 | 0.010293 | 0.010891 | 0 | 0 |
| state5-visual | 0.000819 | 0 | -0.26155 | 0.132673 | 0 |
| state5-frontoparietal | 0 | -0.15049 | 0 | -0.14253 | 0 |
| state5-salience | 0 | -0.06104 | 0.258195 | 0 | 0 |
| state5-subcortical | 0 | 0 | 0.367384 | 0 | 0 |
| state5-dorsalAttention | 0 | 0 | -0.30603 | 0 | 0 |

*This table presents the loading coefficients of the five extracted sparse components (sPC1–sPC5). Non-zero loading values indicate features contributing to the corresponding sparse component, whereas zero values indicate features excluded by the sparsity constraint.*

***Supplementary Table S15. Silhouette coefficients across candidate cluster numbers***

| *K* | *silhouette* |
| --- | --- |
| K=2 | 0.2111 |
| K=3 | 0.1110 |
| K=4 | 0.0933 |
| K=5 | 0.0928 |
| K=6 | 0.0934 |
| K=7 | 0.0933 |
| K=8 | 0.0697 |
| K=9 | 0.0651 |

***Supplementary Table S16. Distribution of neurodynamic architecture composition across sites***

| ***Site*** | ***A1, n (M/F)*** | ***A2, n (M/F)*** | ***A1 Age*** | ***A2 Age*** | ***A1 HAMD*** | ***A2 HAMD*** |
| --- | --- | --- | --- | --- | --- | --- |
| S1 | 21 (10/11) | 50 (20/30) | 30.52 ± 7.37 | 32.14 ± 8.20 | 23.86 ± 4.82 | 25.04 ± 4.76 |
| S2 | 9 (1/8) | 20 (4/16) | 40.11 ± 8.10 | 44.70 ± 14.33 | 22.22 ± 2.82 | 23.90 ± 3.18 |
| S3 | 18 (9/9) | 9 (3/6) | 20.61 ± 1.97 | 19.89 ± 1.54 | NA | NA |
| S5 | 0 (0/0) | 3 (2/1) | NA | 30.33 ± 5.13 | NA | NA |
| S6 | 3 (0/3) | 11 (5/6) | 26.67 ± 4.04 | 31.18 ± 12.34 | 22.67 ± 4.16 | 23.45 ± 7.12 |
| S7 | 6 (2/4) | 26 (10/16) | 42.00 ± 15.05 | 42.19 ± 11.62 | 24.00 ± 5.02 | 21.88 ± 2.96 |
| S8 | 16 (8/8) | 37 (11/26) | 30.06 ± 10.83 | 28.03 ± 11.07 | 22.12 ± 10.45 | 23.08 ± 7.41 |
| S9 | 23 (14/9) | 27 (10/17) | 27.00 ± 8.15 | 29.04 ± 9.42 | NA | NA |
| S10 | 19 (9/10) | 29 (12/17) | 32.26 ± 9.89 | 33.93 ± 11.40 | 19.63 ± 4.21 | 21.52 ± 3.43 |
| S11 | 3 (0/3) | 27 (13/14) | 22.33 ± 0.58 | 30.19 ± 10.40 | NA | 23.38 ± 4.37 |
| S12 | 8 (2/6) | 24 (5/19) | 31.38 ± 7.07 | 35.58 ± 10.33 | 24.75 ± 6.48 | 24.08 ± 4.07 |
| S13 | 16 (7/9) | 9 (4/5) | 33.75 ± 10.55 | 29.67 ± 6.24 | 26.12 ± 3.81 | 23.67 ± 3.74 |
| S14 | 41 (16/25) | 21 (7/14) | 29.56 ± 7.11 | 31.90 ± 7.03 | 20.93 ± 3.11 | 22.05 ± 4.07 |
| S15 | 14 (5/9) | 29 (11/18) | 49.64 ± 10.60 | 44.48 ± 15.73 | 25.14 ± 4.07 | 27.72 ± 5.59 |
| S16 | 9 (4/5) | 20 (13/7) | 34.00 ± 10.46 | 29.45 ± 7.96 | 22.44 ± 2.88 | 21.10 ± 3.40 |
| S17 | 10 (7/3) | 31 (7/24) | 20.40 ± 3.03 | 21.39 ± 3.66 | 23.00 ± 5.58 | 19.68 ± 5.56 |
| S18 | 10 (5/5) | 11 (2/9) | 31.00 ± 7.92 | 31.82 ± 6.37 | 23.80 ± 5.61 | 25.64 ± 5.61 |
| S20 | 65 (22/43) | 213 (77/136) | 39.68 ± 15.69 | 38.30 ± 12.90 | 21.21 ± 5.21 | 20.87 ± 5.73 |
| S21 | 18 (11/7) | 67 (27/40) | 35.94 ± 10.75 | 34.54 ± 13.17 | 14.50 ± 8.74 | 14.41 ± 7.92 |
| S22 | 11 (5/6) | 18 (9/9) | 33.55 ± 9.69 | 34.44 ± 10.46 | 24.18 ± 6.10 | 21.89 ± 4.27 |
| S23 | 7 (4/3) | 25 (12/13) | 30.57 ± 11.33 | 23.48 ± 7.26 | 22.57 ± 5.13 | 17.44 ± 8.87 |
| S24 | 16 (4/12) | 16 (4/12) | 31.81 ± 9.42 | 31.38 ± 7.74 | 25.06 ± 3.89 | 23.19 ± 3.89 |
| S25 | 48 (12/36) | 40 (11/29) | 65.77 ± 6.82 | 65.12 ± 6.62 | 5.77 ± 4.36 | 6.05 ± 5.11 |

*Values statistics were calculated using available data; NA indicates that HAMD data were unavailable. A1: architecture1; A2:architecture2;*

***Supplementary Table S17. Comparison of two clustering strategies in the validation cohort***

| Independent reclustering | Centroid-based Architecture 1 | Centroid-based Architecture 2 | Total |
| --- | --- | --- | --- |
| Architecture 1 | 18 (100.0%) | 0 (0.0%) | 18 |
| Architecture 2 | 23 (48.9%) | 24 (51.1%) | 47 |
| Total | 41 | 24 | 65 |

*Agreement was asymmetric: all 18 participants assigned to Architecture 1 by independent re-clustering retained this assignment under centroid-based transfer, whereas the 47 independently re-clustered Architecture 2 participants were divided between Architecture 1 (n = 23) and Architecture 2 (n = 24) under centroid-based assignment.*
